# A virus co-opts vector’s m^6^A machinery for adaptive compensation

**DOI:** 10.64898/2026.08.27.747426

**Authors:** Buli Fu, Jinyu Hu, Qun Liu, Rong Zhang, Xiaojie Wu, Xinfeng Guo, Shaonan Liu, Mingjiao Huang, Peipan Gong, Xuegao Wei, Jing Yang, Qimei Tan, Jinjin Liang, Chao He, Xuguo Zhou, Ralf Nauen, Xiaobin Shi, Youjun Zhang, Chris Bass, Xin Yang

## Abstract

Insect vectors harbor diverse viruses that threaten human health and agricultural production, yet the genetic basis of mutualistic virus-vector interactions remain elusive. This underscores a long-standing paradox in pest management: viruses that cause destructive diseases also facilitate vector adaptation in ways that benefit both parties. Here we show that the globally devastating tomato yellow leaf curl virus (TYLCV) acts as a “Cooperative Partner”, co-opting its vector’s m^6^A epitranscriptomic machinery to offset the reproductive fitness costs of neonicotinoid resistance in the whitefly. In resistant vectors, TYLCV infection alleviates reproductive deficits via the m^6^A-dependent pathway, where the methyltransferase METTL14 and demethylase ALKBH4 coordinately stabilize *vitellogenin* (Vg) transcripts to boost female fecundity. Mechanistically, the viral C2 and CP proteins directly interact with METTL14 and ALKBH4, respectively, remodeling their RNA binding affinity to facilitate m^6^A modification on *Vg*. Disrupting either of these viral-vector protein interactions restores a reproductive cost phenotype in viruliferous resistant vectors, offering novel targets for sustainable pest control. Our findings uncover a fundamental role of epigenetic marks in adaptive trait, reshaping beneficial virus-vector relationships and providing new insights for innovative public health and crop protection strategies to address global challenges.

## Introduction

Insect-borne viruses annually cause diverse diseases in humans and crops, threatening global health and food security (1–4). Decades of research have revealed that virus–insect vector interactions span a spectrum from beneficial mutualism to detrimental parasitism (5–7). A key unresolved question lies at the heart of this field: why do some insect vectors not only tolerate but also benefit from the viruses they harbor? This question underscores a growing paradox: viruses that cause devastating diseases also enhance vector adaptation in ways that benefit both parties (8–11). Resolving this paradox challenges a century of pest control dogma and uncovers an unexpected alliance hidden in plain sight. Yet a critical knowledge gap persists: what genetic mechanisms enable the evolution and persistence of such mutually beneficial relationships?

Epigenetic modifications, which can rapidly respond to environmental cues to fine-tune gene expression, have emerged as key regulators of adaptive trait evolution (12). Among these, N⁶-methyladenosine (m^6^A)—the most prevalent internal mRNA modification—governs RNA stability, splicing, and translation, and is dynamically regulated by methyltransferases (writers), demethylases (erasers), and m^6^A-binding proteins (readers) (13–15). Growing studies have implicated m^6^A in vector adaptation (16–18) and viral replication (19–22), yet direct evidence linking m^6^A as a central hub in the mutualistic interactions between viruses and their vectors has been conspicuously absent—until now. This knowledge gap carries profound agricultural implications, as approximately 80% of plant viruses are transmitted by insect vectors (23–25). Over the past century, intensive insecticide use has driven rapid resistance evolution in vectors (26–30). Insecticide resistance often incurs fitness costs (31–33), while some insects can acquire endogenous modifiers to mitigate these costs during long-term evolution (34, 35). Understanding the evolutionary logic of such fitness trade-offs is central to sustainable pest management (36, 37). However, whether and how plant viruses influence insecticide resistance and the associated fitness costs remains completely unknown.

Here, we show that tomato yellow leaf curl virus (TYLCV) (38–45), a globally devastating plant virus, serves as an intimate partner to rescue the reproductive costs associated with neonicotinoids resistance in its vector *Bemisia tabaci* (46–51) through employing the vector’s m^6^A epitranscriptomic machinery. Our findings not only resolve the long-standing paradox of virus-vector mutualism but also provide a framework for developing innovative strategies to protect human health and crop production.

## Results

### Field-evolved neonicotinoids resistance correlates with TYLCV infection in the vector B. tabaci

Outbreaks of *B. tabaci* and TYLCV have currently caused widespread damage to a wide host range of agricultural crops (Fig. S1). This outcome might be attributed to the evolution of vector resistance to insecticides and TYLCV spread. We thus investigated neonicotinoid resistance and TYLCV infection in the field-sampled vectors of *B. tabaci* MED from 2023 to 2025 (Table S1). All field populations developed high level of resistance to neonicotinoid imidacloprid and thiamethoxam compared to the susceptible reference strain (Fig. S2; Table S2 and S3). Strikingly, we found a strong positive correlation between resistance level and TYLCV infection prevalence: 100% of the extremely high-resistant whiteflies were infected with TYLCV, while the infection rate declined progressively in lower-resistant populations (Fig. 1; Fig. S2; Table S2 and S3). These data reveal a close link between viral infection and insecticide resistance, prompting us to investigate the underlying mechanism.

**Fig. 1.**
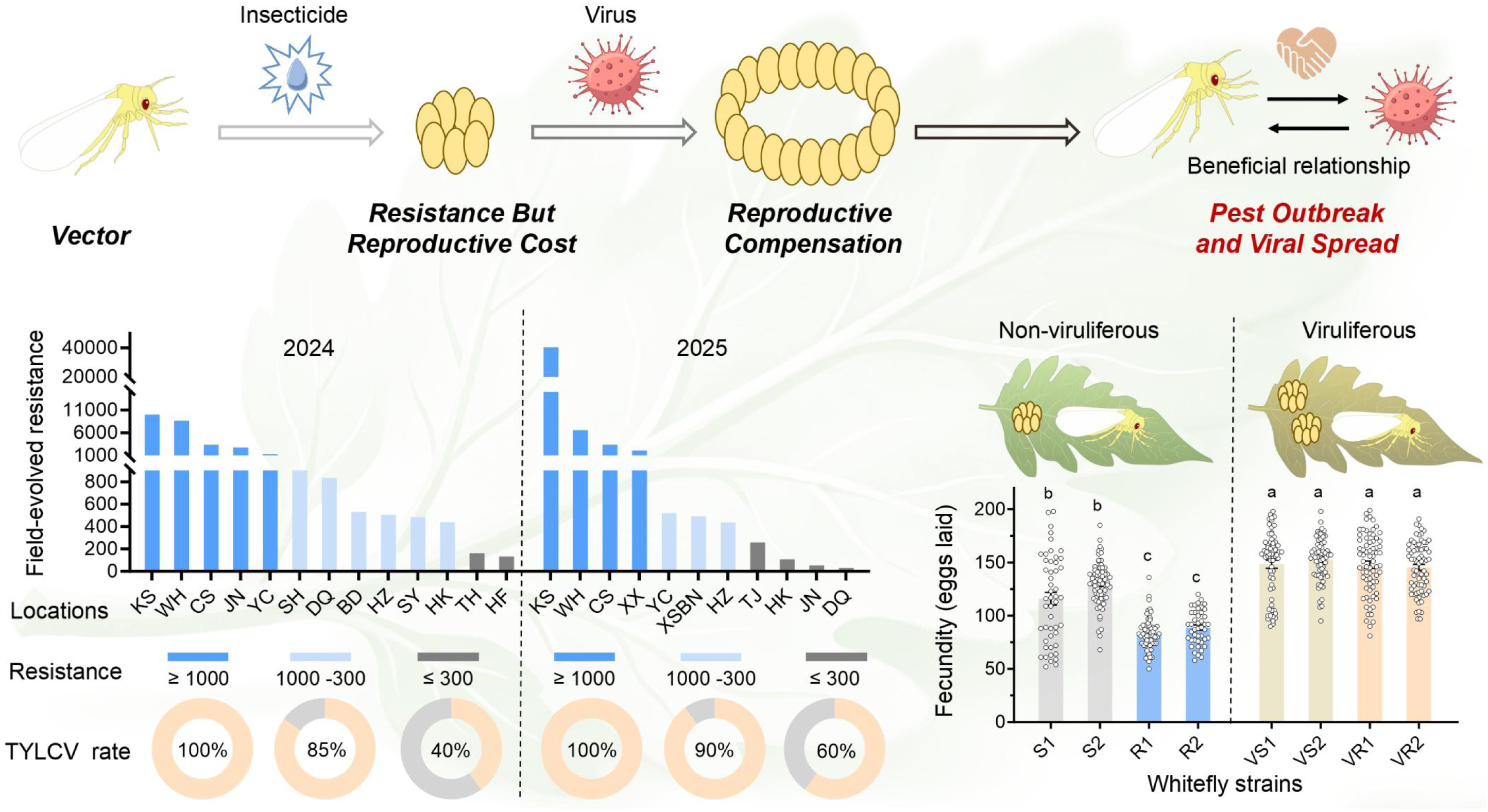
The plant virus TYLCV rescues reproductive cost of neonicotinoid resistance in its vector *B. tabaci* to reshape their beneficial relationship. Widespread of neonicotinoid resistance in field vectors is highly infected by TYLCV with varying proportions depending on the resistance level. Notably, the evolution of neonicotinoid resistance comes with significant reproductive cost, whereas this phenotype is recovered by TYLCV infection in the viruliferous neonicotinoid-resistant vectors.

### TYLCV infection rescues fitness costs of neonicotinoid resistance in its vector B. tabaci

To dissect the interplay between plant virus and insecticide resistance (Fig. S3A), we designed a TYLCV infection experiment using indoor neonicotinoid-resistant (R1 and R2) and susceptible (S1 and S2) strains of *B. tabaci* MED (Table S4). Insecticide bioassays (Fig. S3B) confirmed that viral infection reduced the sensitivity of both susceptible (Fig. S3C, D) and resistant (Fig. S4A, B) whiteflies to neonicotinoids imidacloprid and thiamethoxam (Table S5). We next performed life-table analysis to quantify fitness costs associated with neonicotinoids resistance. Non-viruliferous resistant strains (R1, R2) exhibited longer development, lower survival, and reduced reproduction, compared with susceptible counterparts (S1, S2) (Figs. S5 and S6; Tables S6 and S7). Notably, these deficits were fully recovered by TYLCV infection: viruliferous resistant strains (VR1, VR2) had extended oviposition periods, longer adult lifespans, and higher fecundity than their non-viruliferous counterparts (R1, R2) (Figs. S5 and S6; Tables S6 and S7). The relative fitness (*Rf*), quantified using the intrinsic rate of natural increase (*r*), approached unity in both VR strains (VR1: 1.0009; VR2: 0.9757) when normalized to the susceptible S1 reference (Table S8). Collectively, these data demonstrate that TYLCV infection completely offsets the fitness costs associated with neonicotinoid resistance in *B. tabaci* vectors.

### TYLCV infection specifically alleviates resistance-associated reproductive costs

Our previous work has demonstrated that neonicotinoid resistance imposes severe reproductive burdens in non-viruliferous *B. tabaci* (33). We therefore investigated whether TYLCV infection counteracts such deficits via reproductive compensation. Based on net reproductive rate (*R_0_*), non-viruliferous resistant strains showed markedly suppressed relative fitness (R1: 0.5904; R2: 0.4559) compared with the susceptible S1 reference. In contrast, viral infection substantially restored reproductive fitness in resistant whiteflies (VR1: 1.2363; VR2: 1.3272) (Table S8). Fecundity phenotyping further validated this pattern: viruliferous resistant strains (VR1, VR2) produced significantly more eggs (Fig. 1) and exhibited higher net maternity (*lₓmₓ*) than their non-viruliferous counterparts (R1, R2) (Fig. S6A). Consequently, TYLCV infection specifically mitigates the reproductive cost.

To solidify this reproductive phenotype, we quantified female fecundity under paired (1♀:1♂), single (virgin, 1♀), and group (15♀:15♂) conditions (Fig. S7A). In both paired and single settings, VR1 and VR2 females laid significantly more eggs throughout adulthood than non-viruliferous resistant R1 and R2 females (Fig. S7B, C). Under group housing, viral infection consistently boosted egg output across early, middle and late adult stages (Fig. S7D). Taken together, these data demonstrate that TYLCV specifically counteracts the reproductive cost of insecticide resistance, thereby establishing a mutualistic relationship between the virus and its vector (Fig. 1).

### Vitellogenin upregulation underlies reproductive compensation

To investigate the molecular mediator of this reproductive compensation, we examined four key oogenesis genes (Fig. S8A) previously implicated in this cost (33). Among these, Vg showed elevated expression in viruliferous resistant (VR1, VR2) females compared with non-viruliferous resistant (R1, R2) females, as determined by qPCR and western blot (Fig. 2A; Fig. S8B, C). Additionally, TYLCV acquisition also increased Vg protein levels in non-viruliferous R1 and R2 strains over time (Fig. S8D). Immunofluorescence (IF) assays further confirmed stronger Vg protein signal in oocytes of VR1 and VR2 relative to R1 and R2 (Fig. S8E). These findings suggest that TYLCV-induced upregulation of Vg correlates to reproductive compensation.

**Fig. 2.**
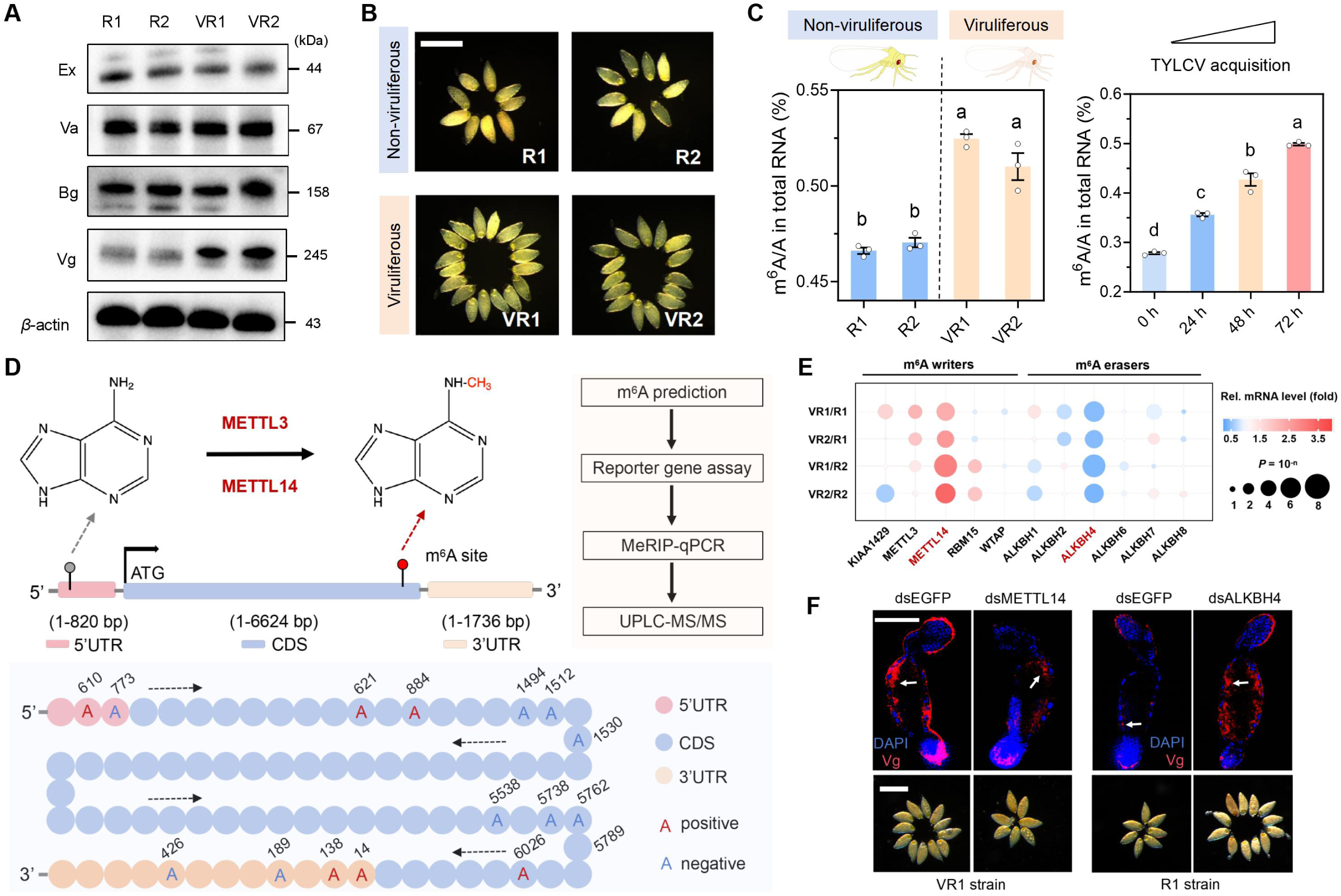
TYLCV-mediated vitellogein (Vg) upregulation compensates the reproductive cost of neonicotinoids resistance via m^6^A modification. (**A**) Western blot analysis of key oogenesis genes involved in reproductive compensation in female vectors. (**B**) Representative images displaying mature oocytes in female vectors. Scale bars, 200 μm. (**C**) UPLC-MS/MS analysis of the global m^6^A level of RNA samples extracted from female vectors (*P* < 0.05, Tukey’s HSD multiple comparisons test). (**D**) Identification of functional m^6^A sites of *Vg* gene. (**E**) qRT-PCR analysis of relative gene expression of key m^6^A writers and erasers in female vectors. (**F**) Functional validation of METTL14 and ALKBH4 in regulation of Vg and phenotypic observation following RNAi experiments. Scar bar, 50 μm for immunofluorescence image, 200 μm for oocyte image.

Vg is indispensable for oogenesis in oviparous animals (52, 53), providing nutrients for maturing oocytes (Fig. S8F). We thus explored whether TYLCV impacts oocyte development to offset reproductive cost. Ovarian dissection revealed that mature oocyte number was increased by day 12 post-eclosion and declined thereafter (Fig. S8G). Resistant strains (R1, R2) harbored fewer mature oocytes than susceptible strains (S1 and S2) at all time points, whereas TYLCV infection significantly restored oocyte maturation in VR1 and VR2 strains (Fig. 2B; Fig. S8G-I). To test causality, we performed RNAi-mediated *Vg* knockdown in VR1 and VR2 (Fig. S9A). Silencing of *Vg* reduced egg production and the number of mature oocytes compared with dsEGFP controls (Fig. S9B-D). Consequently, TYLCV-induced Vg upregulation is responsible for the reproductive compensation.

### m^6^A modification correlates with TYLCV infection

Given our previous work on epigenetic regulation of insecticide resistance (36, 54), we next investigated whether key RNA modifications (m^6^A, m¹A, m⁵C) epigenetically modulate TYLCV-mediated Vg upregulation. Dot blot assays with methylation-specific antibodies showed higher m^6^A signals in viruliferous resistant (VR1, VR2) than non-viruliferous counterparts (R1, R2), with no differences in m¹A or m⁵C (Fig. S10A). UPLC-MS/MS corroborated this result with the m^6^A abundance of total RNA found to be significantly higher in VR1 and VR2 strains compared to R1 and R2 (Fig. 2C; Fig. S10B). To further confirm TYLCV-driven m^6^A modification, we conducted TYLCV acquisition tests in the non-viruliferous R1 and R2 strain. Exposure to TYLCV for 24, 48 and 72 h significantly increased m^6^A level over time compared to the 0 h control (Fig. 2C; Fig. S10C, D). Collectively, TYLCV-induced m^6^A upregulation is associated with Vg upregulation leading to reproductive compensation.

### Mapping the m^6^A landscape of Vg gene

To test whether *Vg* is epigenetically regulated by m^6^A, we first performed Rapid Amplification of cDNA Ends (RACE) technique to generate full-length cDNAs (Fig. S10E). SRAMP prediction identified 83 putative m^6^A sites with conserved RRACH motifs (Table S14). These sites were distributed in UTRs and throughout the CDS (including exon–exon junctions), with enrichment near long internal exons, start/stop codons, and three 100 bp-window m^6^A clusters deposited within the CDS (Fig. S10F-H), suggesting potential *Vg* regulatory roles. To experimentally validate this (Fig. 2D), we first mutated all 83 adenosine (A) sites to guanine (G) and compared mutants (M) with wild-type (WT) Vg coding sequence in dual-luciferase reporter assays and qPCR using Drosophila S2 cells (Fig. S11A). A total of 53 sites (63.85%) showed no difference in reporter activity or transcription level between WT and M, indicating no regulatory function (Fig. S11B, C). Of the 30 functional sites, 11 positively and 19 negatively regulated *Vg* (Fig. S11B, C), revealing the regulation of *Vg* via these m^6^A sites.

m^6^A modification is often catalyzed by key methyltransferases METTL3 and METTL14 (36, 54). MeRIP-qPCR assays on VR1 adult females identified 13 of the 30 functional sites enriched with m^6^A antibody relative to IgG control (Fig. S12A). All 13 sites except A6360s were highly enriched for METTL3- and METTL14-specific antibodies relative to IgG (Fig. S12B), indicating binding of METTL3 and METTL14 to these m^6^A sites. To confirm methylation effects on these sites, we generated 15 bp single-stranded RNAs (ssRNAs) covering each m^6^A site and examined methylation capacity using UPLC-MS/MS. Co-incubation of recombinant METTL3 or METTL14 with these ssRNAs increased methylation levels compared with controls (without enzyme) in all cases (Fig. S12C). Altogether, these *in vivo* and *in vitro* data identify functional *Vg* m^6^A sites targeted by METTL3 and METTL14 (Fig. 2D; Fig. S12D).

### METTL14 and ALKBH4 coordinate to enhance Vg overexpression

Given TYLCV-induced m^6^A elevation in *B. tabaci* (Fig. 2C), we examined if this phenotype stems from altered expression of m^6^A writers or erasers. qRT-PCR analysis of 11 writer/eraser genes from the *B. tabaci* genome (55, 56) showed that, viruliferous resistant strains (VR1, VR2) exhibited upregulation of the writer *METTL14* and downregulation of the eraser *ALKBH4* compared with non-viruliferous resistant strains (R1, R2) (Fig. 2E). These results were confirmed by western blot (Fig. S13A), implicating METTL14 and ALKBH4 in TYLCV-induced m^6^A upregulation. Additionally, TYLCV acquisition increased METTL14 and decreased ALKBH4 expression in non-viruliferous R1 and R2 strain (Fig. S13B), further confirming their role in m^6^A elevation.

To determine whether METTL14 or ALKBH4 regulates *Vg*, we performed RNAi to knockdown these two genes in the adult females of VR1 strain. Silencing of *METTL14* decreased Vg protein levels in VR1 strain, whereas knockdown of *ALKBH4* increased Vg expression in R1 strain (Fig. S13C). IF assays further confirmed reduced Vg signal in mature oocytes after silencing of *METTL14* and increased signal after knockdown of *ALKBH4* (Fig. 2F). To elucidate if the action of METTL14 or ALKBH4 on *Vg* influences reproductive phenotype, we then assessed female fecundity and oocyte maturation following RNAi knockdown of each gene in VR1 females. Depletion of *METTL14* reduced fecundity and the number of mature oocytes at days 5 and 10 compared with dsEGFP (Fig. S13D, E), whereas silencing of ALKBH4 had an opposite effect (Fig. S13F, G). Collectively, METTL14 and ALKBH4 coordinate to enhance Vg expression, thereby mediating reproductive compensation in viruliferous neonicotinoid resistant *B. tabaci*.

### METTL14 and ALKBH4 selectively target Vg m^6^A sites to enhance its mRNA stability

To elucidate how METTL14 and ALKBH4 modulate Vg via m^6^A modification, we performed MeRIP-qPCR targeting the functional m^6^A sites mapped previously (Fig. 2D; Fig. S12D). Compared with non-viruliferous resistant strains (R1, R2), RNA immunoprecipitated with the m^6^A antibody was highly enriched at eight m^6^A sites in the VR1 and VR2 strain (Fig. S14A). Building on this, RNA immunoprecipitated with the METTL14 antibody revealed enrichment at four of the eight sites, while ALKBH4 antibody showed significant depletion at the same four sites (3′UTR: A14, A138; CDS: A621, A884) in VR1 and VR2 relative to R1 and R2 (Fig. S14A). Since the four sites positively regulate *Vg* (Fig. 2D), we conclude that METTL14 and ALKBH4 selectively target these m^6^A sites to drive Vg overexpression (Fig. 3A). We then applied a single-base elongation-and ligation-based qPCR amplification method (SELECT) to verify the four m^6^A sites at single-base resolution (57, 58). In VR1 strain, each candidate adenine (A site) exhibited higher amplification curves (C_T_ values) than surrounding non-m^6^A sites (N sites), confirming m^6^A modification on these sites (Fig. S14B). SELECT assays further revealed higher C_T_ values at these four sites in equivalent RNA samples of VR1 and VR2 strains compared to R1 and R2 (Fig. 3B; Fig. S15A), indicating TYLCV-induced m^6^A elevation at these sites.

**Fig. 3.**
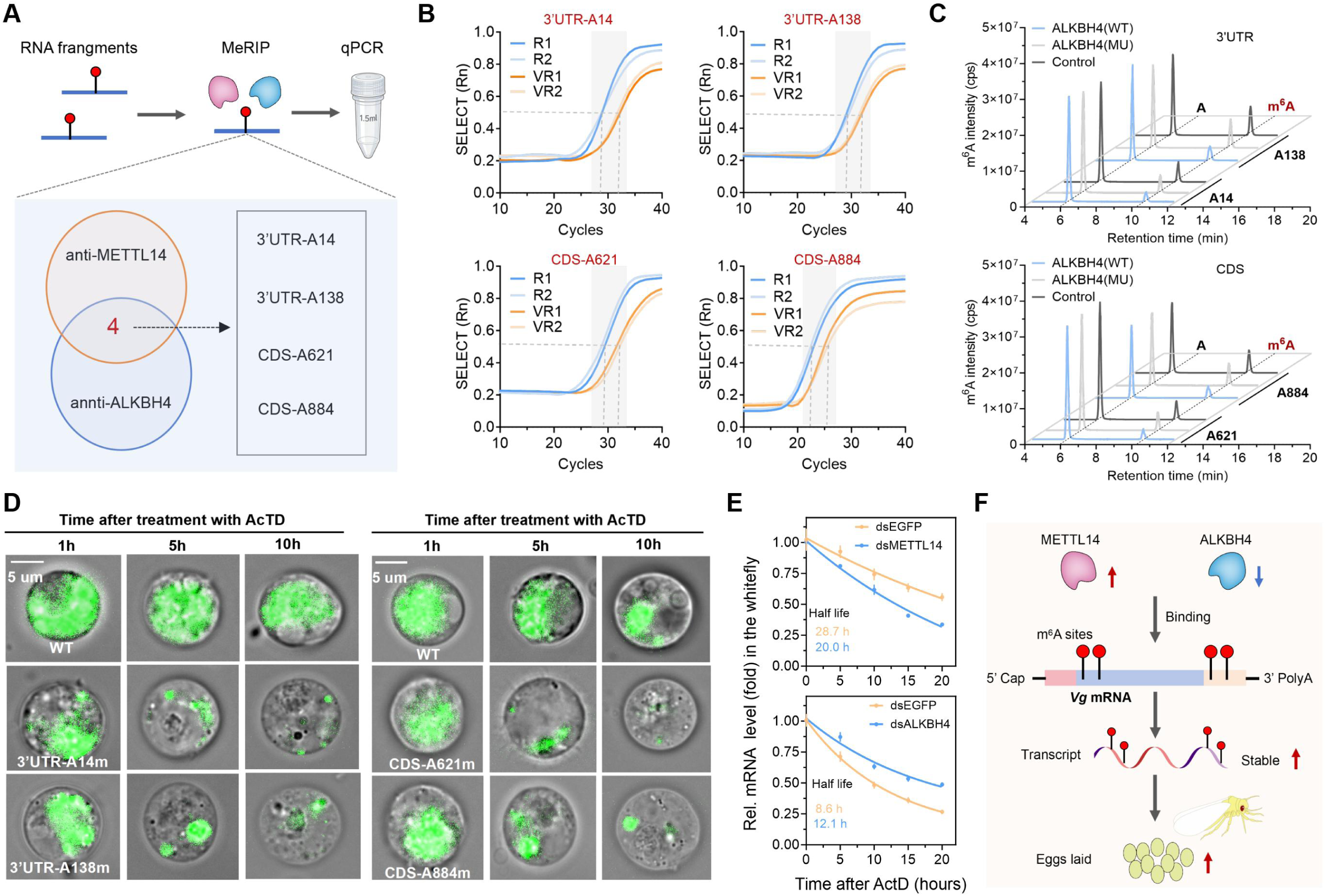
METTL14 and ALKBH4 coordinate to enhance *Vg* mRNA stability through selectively targeting functional m^6^A sites. (**A**) Schematic diagram of screening functional m^6^A sites of *Vg* by MeRIP-qPCR analysis. (**B**) The SELECT results for comparison of m^6^A modification on the functional m^6^A sites of *Vg* at single-base resolution in the vector. (**C**) UPLC-MS/MS analysis of demethylation activity of recombinant ALKBH4 on the functional m^6^A sites of *Vg*. (**D**) mRNA stability of *Vg* following transfection of wide-type (WT) and mutant (m) constructs in to Drosophila S2 cells. (**E**) mRNA stability of *Vg* post RNAi knockdown of METTL14 and ALKBH4 in the vector. (**F**) A working model for the coordinate role of METTL14 and ALKBH4 in enhancing *Vg* mRNA stability that contributes to compensating the reproductive cost of neonicotinoids resistance in the viruliferous vector.

To confirm site-specific m^6^A regulation by METTL14 and ALKBH4, we performed SELECT assays on RNA co-incubated with recombinant METTL14 or ALKBH4. METTL14 reduced the final elongated-ligated product relative to controls (no enzyme or mutant METTL14), whereas ALKBH4 had the opposite effect versus mutant ALKBH4 or EDTA-treated controls (Fig. S16A, B), demonstrating their antagonistic action at the four m^6^A sites. We have recently demonstrated that ALKBH4 functions as a key m^6^A demethylase in the whitefly, catalyzing the oxidative reversal of mRNA m6A modifications both in vitro and in vivo (56). UPLC-MS/MS was then used to examine ALKBH4-mediated demethylation of these four sites. Co-incubation of recombinant wild-type (WT) ALKBH4 with m^6^A-tagged ssRNAs containing each site reduced methylation levels by ∼50% relative to no-enzyme controls (Fig. 3C). Conversely, functional loss-of-mutation (MU) of ALKBH4 increased methylation (Fig. 3C), confirming ALKBH4 demethylates these Vg m^6^A sites. To functionally test the direct binding of METTL14 or ALKBH4 to the four m^6^A sites, we performed electrophoretic mobility shift assay (EMSA) using 25 bp ssRNA sequences containing each site as a probe. Regarding METTL14, binding was confirmed by a shifted band that disappeared with unlabeled competitor or mutated probes (Fig. S17A). EMSA also demonstrated direct binding of recombinant ALKBH4 to these sites when probes tagged m^6^A at the adenosine, whereas no shift was observed with non-m^6^A modified probes (Fig. S17B).

To quantify Vg regulation by these four sites, we constructed Vg-coding sequence constructs containing WT m^6^A sites, A-to-G mutant (MU) m^6^A sites, or m^6^A-tagged at adenosine sites, which were cloned into the pAC5.1b-G3-X2-EGFP vector (Fig. S15B) and transfected into Drosophila S2 cells. WT cells showed the highest GFP fluorescence, followed by m^6^A-tagged cells, with MU cells the lowest (Fig. S15C)—consistent with *Vg* mRNA levels (Fig. S15D). These results indicate that m^6^A modification at these sites enhances Vg expression. Given that m^6^A modification primarily regulates mRNA stability (10, 13), we hypothesized METTL14 andALKBH4 could influence *Vg* mRNA stability via these four sites. mRNA decay assays showed that MU cells had significantly reduced *Vg* mRNA half-life compared to WT (Fig. S17C), corroborated by GFP fluorescence of *Vg* reporter constructs (Fig. 3D) following Actinomycin D (ActD) treatment. To clarify this result, we performed SELECT assays after RNAi knockdown of METTL14 or ALKBH4 in VR1 strains. Silencing of METTL14 decreased C_T_ values at all four m^6^A sites compared to dsEGFP controls, whereas knockdown of ALKBH4 showed an opposite effect (Fig. S18A, B), confirming the importance of METTL14 and ALKBH4 in site-specific m^6^A modification. Additionally, METTL14 depletion reduced Vg transcript half-life, while ALKBH4 knockdown increased it relative to dsEGFP, as determined by ActD-based mRNA decay assays (Fig. 3E). Conclusively, the above findings demonstrate that METTL14- and ALKBH4-mediated m^6^A modification at the four functional sites (3’UTR: A14 and A138; CDS: A621 and A884) enhances *Vg* mRNA stability, thereby contributing to reproductive compensation in neonicotinoid resistant viruliferous *B. tabaci* (Fig. 3F).

### TYLCV proteins C2 and CP interact specifically with METTL14 and ALKBH4 of B. tabaci, respectively

How does TYLCV infection influence m^6^A modification in *B. tabaci*To answer this question, we hypothesized that one or more TYLCV structural proteins (59–61) might interact with vector’s METTL14 or ALKBH4 (Fig. S19A). Yeast two-hybrid (Y2H) assays identified a strong interaction between METTL14 and C2, as well as between ALKBH4 and CP, C2, and C3 (Fig. S19B). Immunoprecipitation (IP) assay confirmed direct interaction between METTL14 and C2, and between ALKBH4 and CP in VR1 strain (Fig. 4A), whereas ALKBH4 did not interact with C2 or C3 (Fig. S19C). IF staining assays revealed co-localization of METTL14-C2 and ALKBH4-CP interaction in mature oocytes of VR1 and VR2 strains (Fig. 4B), which was further validated by GST pull-down assays (Fig. S19D). Taken together, these *in vitro* and *in vivo* data establish that the viral C2 and CP proteins interact specifically with *B. tabaci* METTL14 and ALKBH4, respectively.

**Fig. 4.**
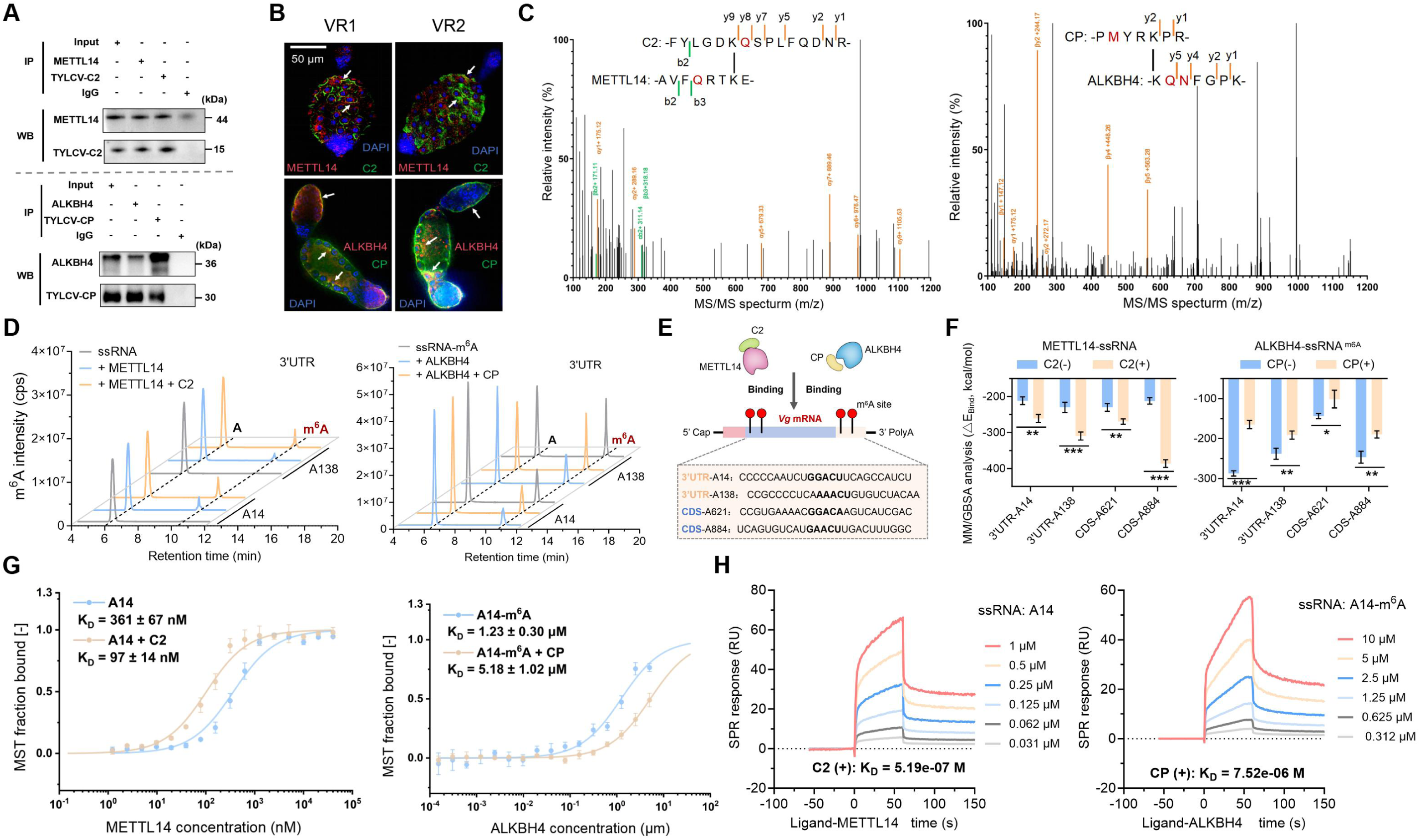
The binding of TYLCV protein C2 and CP to vector’s METTL14 and ALKBH4 enhances m^6^A modification on Vg. (**A**) Immunoprecipitation analysis of METTL14-C2 and ALKBH4-CP interaction in the vector. (**B**) Immunofluorescence staining assays of the colocalization of METTL14 with C2, and ALKBH4 with CP in oocytes of the vector. (**C**) The cross-linking mass spectrometry (XL-MS) analysis of binding interfaces of METTL14-C2 or ALKBH4-CP complex. (**D**) UPLC-MS/MS analysis of the effect of recombinant C2 or CP on m^6^A modification of *Vg* by coincubation with METTL14 or ALKBH4. (**E**) Schematic diagram for testing the impact of METTL14-C2 or ALKBH4-CP complex on binding capacity to *Vg* m^6^A sites. (**F**) The simulated binding energy of METTL14 or ALKBH4 to the ssRNAs in the presence or absence of C2 or CP by molecular docking and molecular dynamics simulations. (**G**) The MST assay for detecting binding affinities of recombinant METTL14 and ALKBH4 to substrate ssRNA in the presence of viral C2 or CP protein. (**H**) The SPR assay for detecting binding affinities of recombinant METTL14 and ALKBH4 to substrate ssRNA in the presence of viral C2 or CP protein.

To characterize METTL14-C2 and ALKBH4-CP interactions, we attempted to experimentally resolve the crystal structure of these proteins but were unsuccessful. Instead, we performed structural predictions using AlphaFold3 (Fig. S19E). Molecular docking identified a series of amino acids as potentially involved in the interaction between METTL14 (eight residues) and C2 (ten residues) or between ALKBH4 (ten residues) and CP (nine residues), which were formed by hydrogen, π–π and salt bridge bonding interactions (Tables S16 and S17). Subsequently, 50ns molecular dynamic simulations were performed to relax the protein-ligand binding complex. The resulting RMSD, RMSF and RG identified an energetically favourable binding conformation (△E_Bind_) (METTL14-C2 complex: -149.15 kcal/mol, ALKBH4-CP complex: -143.42 kcal/mol) (Tables S16 and S17). The binding interfaces of METTL14-C2 or ALKBH4-CP complex were then built based on cross-linking mass spectrometry (XL-MS) (Fig. S20). We identified an unique inter-linked peptide pairs in each complex, mapping the binding epitope to the METTL14 residues [AVFQRTKE (aa 243-250)] and the C2 residues [FYLGDKQSPLFQDNR (aa 68-81)], and the ALKBH4 residues [KQNFGPK(aa 106-112)] and CP residues [PMYRKPR(49–55)] (Fig. 4C). These XL-MS data corroborate the overall fold predicted by AlphaFold3 predictions while defining their interaction interface at residue-level resolution (Fig. S20A, B). To validate these binding residues, we generated mutant constructs (BD-METTL14, AD-C2, BD-ALKBH4, AD-CP) and performed Y2H assays. Unlike wild-type constructs, co-transformed mutant cells failed to grow on quadruple dropout media (Fig. S21). These findings confirm that TYLCV C2 and CP proteins directly interact with *B. tabaci* METTL14 and ALKBH4, respectively.

### The viral proteins C2 and CP facilitate m^6^A modification of Vg via interactions with METTL14 and ALKBH4

Given the established METTL14-C2 and ALKBH4-CP interactions, we next investigated whether C2 and CP regulate m^6^A modification at the four functional *Vg* m^6^A sites. UPLC-MS/MS analyses showed that co-incubation of recombinant C2 with METTL14, or recombinant CP with ALKBH4, significantly increased m^6^A levels at these sites compared to incubation with METTL14 or ALKBH4 alone (Fig. 4D; Fig. S22). To further validate this, we generated recombinant mutants (MU) with disrupted binding residues (predicted earlier) to abrogate protein-protein interactions. Relative to wild-type (WT) controls, these MU proteins reduced m^6^A levels at all four Vg m^6^A sites (Figs. S23-S24). Collectively, these *in vitro* data demonstrate that METTL14-C2 and ALKBH4-CP interactions facilitate *Vg* m^6^A modification. To confirm the *in vivo* relevance of these findings, we performed SELECT assays after oral ingestion of anti-C2 or anti-CP antibodies (to block METTL14-C2 and ALKBH4-CP interactions, respectively) in VR1 whiteflies. Feeding on anti-C2 or anti-CP antibodies reduced CT values at these sites compared to preimmune serum controls in VR1 whiteflies (Fig. S25). Together, these *in vivo* results indicate that the viral protein C2 and CP facilitate *Vg* m^6^A modification through their interactions with METTL14 and ALKBH4.

We next explored the mechanism by which C2 and CP enhance *Vg* m^6^A modification. Given that METTL14-C2 and ALKBH4-CP interactions alter the conformational states of METTL14 and ALKBH4 (Tables S16 and S17), we hypothesized that these complexes might impact enzymatic activity. Using an ELISA-like assay, we found that METTL14 methylase activity increased with recombinant METTL14 concentration but was unchanged by the presence of C2 at various concentrations (Fig. S26A). Likewise, no changes in m^6^A demethylase activity when C2 was co-incubated with ALKBH4 (Fig. S26B). Thus, METTL14-C2 and ALKBH4-CP interactions do not affect the enzymatic activities of METTL14 or ALKBH4, indicating an alternative mechanism underlying their facilitation of *Vg* m^6^A modification.

### The viral protein C2 enhances the binding affinity of METTL14 to Vg m^6^A sites

Since binding specificity of m^6^A writers to substrate RNA is a key determinant of m^6^A modification, we investigated whether the METTL14-C2 interaction affects METTL14 binding to the four *Vg* m^6^A sites (Fig. 4E). To this end, we performed molecular docking and 50 ns molecular dynamics simulations to visualize conformational changes in of METTL14-ssRNA binding in the presence or absence of C2 (Fig. S26C). In each METTL14-ssRNA complex, a subset of METTL14 residues directly contacted the ssRNA, while this binding mode was altered by the presence of C2 in the ternary METTL14-ssRNA-C2 complex (Table S18). The presence of C2 exhibited significantly lower binding energy (△E_Bind_) than the METTL14-ssRNA binary complex (Fig. 4F; Fig. S27A). These predictions indicate that C2 enhances RNA binding capacity of METTL14 to *Vg* m^6^A sites, thus facilitating m^6^A modification of *Vg*. We then performed microscale thermophoresis (MST) assay to experimentally test the binding affinity of target proteins to substrate RNAs. The recombinant C2 reduced dissociation constant (*K_D_*) values at each *Vg* m^6^A site upon co-incubation with recombinant METTL14 (Fig. 4G; Figs. S28and S29; Table S24), demonstrating an enhanced binding affinity of METTL14 to *Vg* m^6^A sites in the presence of C2. This result was further confirmed by surface plasmon resonance (SPR) analysis on two representative *Vg* m^6^A site (3UTR-A14 and CDS-A621), with a lower binding constant (*K_D_*) value when co-incubation with C2 and METTL14 compared to METTLA4 alone (Fig. 4H; Fig. S32 and Table S26). To further validate this, we mutated key C2 binding residues (MU) involved in hydrogen bonds, π–π interactions, and salt bridges within the METTL14-ssRNAs-C2 complex. Using two representative ssRNAs spanning m^6^A sites (3′UTR-A14, A138), EMSA assays showed that increasing concentrations of wild-type (WT) C2 enhanced the binding of METTL14 to both ssRNAs, whereas the mutant (MU) C2 markedly reduced this binding shift (Fig. S34A, B). Furthermore, HPLC-MS/MS analysis revealed that, the mutant C2 reduced m^6^A levels at each *Vg* m^6^A site upon co-incubation with METTL14 compared to wild-type C2 (WT) (Fig. S34C). Collectively, these data provide evidence that the viral protein C2 strengthens the RNA binding affinity of METTL14, thereby promoting *Vg* m^6^A modification.

### The viral protein CP blocks the binding affinity of ALKBH4 to Vg m^6^A sites

Given the robust ALKBH4-CP interaction (Fig. 4E), we investigated whether CP could impact ALKBH4 binding to *Vg* m^6^A sites. We performed molecular docking and 50 ns molecular dynamics simulations to visualize conformational changes in ALKBH4-ssRNA binding in the presence or absence of CP. Notably, the ALKBH4-ssRNA-CP ternary complex exhibited significantly higher binding energy (ΔE_bind_) than the ALKBH4-ssRNA binary complex (Fig. 27B). This was attributed to CP-induced alterations in the binding state of the ALKBH4-ssRNA complex (Table S19), indicating an inhibited binding of ALKBH4 to substrate ssRNAs.

Given these predictions, we next conducted MST assay to experimentally illuminate the role of CP in binding affinity of ALKBH4 to substrate RNAs. For each *Vg* m^6^A sites, the presence of recombinant CP increased *K_D_* values compared to without CP upon co-incubation with recombinant ALKBH4 (Fig. 4G; Figs. S30 and S31; Table S25), revealing a reduced binding affinity of ALKBH4 to *Vg* m^6^A sites. Based on these, we then performed SPR assay to investigate whether CP influence binding affinity of ALKBH4 to *Vg* m^6^A sites. The *K_D_* values were found to be increased in the presence of CP compared to without (Fig. 4G; Fig. S33 and Table S26), demonstrating a decreased binding affinity of ALKBH4 to *Vg* m^6^A sites and implying an increased m^6^A levels on these sites. To further validate this hypothesis, we mutated key CP residues involved in the ALKBH4-ssRNA-CP complex. The resulting EMSA showed that the ALKBH4-ssRNA binding shift diminished with increasing concentrations of wild-type CP (CP) but was restored when key CP binding residues were mutated (MU) (Fig. S34D, E). HPLC-MS/MS analysis further revealed that the mutant CP (MU) dramatically increased m^6^A levels at each *Vg* m^6^A site upon co-incubation with ALKBH4 compared with wide-type CP (WT) (Fig. S34F). Together, these findings demonstrate that the viral protein CP blocks ALKBH4 binding to *Vg* m^6^A sites, thereby enhancing *Vg* m^6^A modification.

### Inhibition of METTL14-C2 or ALKBH4-CP binding interactions restores a reproductive fitness cost of neonicotinoid resistance

Building on the findings above, we postulated that disrupting METTL14-C2 or ALKBH4-CP interactions could reverse the reproductive compensation phenotype—offering an alternative strategy for pest control. Focusing on the METTL14-C2 interaction, we first used virus-induced gene silencing (VIGS) to generate pTRV2-METTL14 and pTRV2-EGFP (control) vectors (Fig. S35A). After confirming silencing fragments in *Nicotiana benthamiana* (Fig. S35B), viruliferous VR1 whiteflies were fed on these plants for 21 days. METTL14 and Vg expression were markedly decreased at days 7 and 14 in females fed on pTRV2-METTL14 plants compared to controls (Fig. S35C, D), with significantly reduced egg production and fewer mature oocyte numbers (Fig. S35E-G). To gain more evidences, we also performed anti-C2 and anti-CP antibody feeding (blocking METTL14-C2 and ALKBH4-CP interactions, respectively) to examine these effect on reproductive fitness in VR1 strain. Oral ingestion of either anti-C2 (Fig. S36A-E) or anti-CP (Fig. S36F-J) antibody reduced Vg expression, egg production and mature oocytes relative the respective preimmune serum controls, revealing a reproductive cost phenotype. Consequently, inhibition of either METTL14-C2 or ALKBH4-CP interaction restores a reproductive cost of neonicotinoids resistance in the viruliferous whitefly, supporting our control opinion for sustainable pest control.

## Discussion

We uncover a mechanism by which a plant virus acts as an intimate partner, co-opting its vector’s m^6^A machinery to offset the reproductive cost of insecticide resistance in a global pest (Fig. S37). This work unravels a fundamental role for epigenetic regulation in cooperative virus–vector relationships, showing that viruses exploit vector’s epigenetic marks for evolutionary gain. Disrupting these m^6^A-mediated interactions may open new avenues for pest surveillance and control.

Plant viruses often boost vector performance to promote viral spread (62–64). We show that TYLCV restores the reproductive cost of neonicotinoid resistance in *B. tabaci*, consistent with reports that TYLCV increases its vector fitness (65, 66). Thus, this compensatory evolution can offset fitness costs of insecticide resistance through cooperative virus–vector interactions—challenging the conventional view that such costs are inevitable (30). Endogenous modifiers such as *Scalloped wings* in *Lucilia cuprina* (37) and peroxiredoxin in *Nilaparvata lugens* (35) drive similar compensation. We further show that TYLCV-mediated m^6^A modification of *Vg* compensates this reproductive cost, and that m^6^A also regulates juvenile hormone to maintain Bt resistance fitness in *P. xylostella* (36). Hence, epigenetic m^6^A serves both as a key modifier of insecticide resistance fitness costs and as a regulatory hub in virus–vector interactions.

Epigenetic marks change rapidly in response to environmental cues, enabling adaptive trait inheritance (67–69). DNA and RNA methylation enhance plant adaptability under various environmental stress (70–72). In animals, m^6^A modulates aggregation behavior, development and reproduction in several model insects (73). We recently demonstrated that the m^6^A writer METTL3 epigenetically regulates CYP4C64 in whitefly, conferring thiamethoxam resistance (54). Here, we show that METTL14 and ALKBH4 coordinately promote fitness in neonicotinoid-resistant whitefly upon TYLCV infection. Thus, m^6^A emerges as a key epigenetic regulator of adaptive evolution across taxa, and our work provides a framework to dissect genetic changes within the m^6^A pathway (writers– erasers–readers) that govern resistance fitness trade-offs and how they are controlled by other molecular modifications.

Vitellogenin (Vg) is essential for oocyte maturation and reproductive capacity in oviparous animals (52, 53). Beyond this, insect Vg also mediates viral transmission and plant immune defense (74–76). Here, we mapped the m^6^A landscape of whitefly *Vg*, providing a framework for future studies of its multifaceted, m^6^A-dependent functions. Intriguingly, downregulation of Vg underpins reproductive cost in non-viruliferous *B. tabaci* (33); however, TYLCV offsets this cost via METTL14 and ALKBH4, which act on specific m^6^A sites of *Vg* to stabilize its transcripts. Mechanistically, METTL14 binds viral C2 and ALKBH4 binds viral CP, differentially modulating their binding to Vg m^6^A sites—identifying a new mechanism by which C2 and CP facilitate *Vg* m^6^A modification. These findings link m^6^A to adaptive traits in a virus-responding insect vector, highlighting its role in virus–vector interactions. Whether and how changes in the m^6^A pathway in turn benefit the virus merits future investigations.

Compensatory evolution in insecticide-resistant vectors may drive pest resurgence and viral outbreaks, threatening global crop protection. Given the global prevalence of TYLCV and neonicotinoid resistance, disrupting these m^6^A-mediated interactions with virus may establish the m^6^A pathway as a promising target for RNAi-based insecticides and transgenic crops. Beyond agriculture, we unravel a fundamental principle: viruses can co-opt vector epigenetic machinery to offset fitness costs of resistance (Fig. S37), offering a new paradigm for virus-vector coevolution. These insights have broad implications for vector-borne disease control and may inform innovative strategies to safeguard human health and food security.

## Author contributions

Conceptualization: Y.J, Z; X, Y; C. B; Project administration: Y.J, Z; X,Y; C. B; B.L, F; Supervision: Y.J, Z; X, Y; Methodology: B.L, F; J.Y, H; R. Z; X.J, W; X.F, G; X. B, S; Investigation: B.L, F; J.Y, H; R. Z; X.J, W; X.F, G; Validation: B.L, F; J.Y, H; S.N, L; M.J, H; P.P, G; Y.J, Y; Visualization: B.L, F; R. Z; Q.M, T; J.J, L; X.G, W; C, H; Writing-original draft: B.L, F; X. Y; Q, L; Writing-review & editing: B.L, F; Y.J, Z; X. Y; Q, L; C. B; X.G, Z.

## Data and materials availability

All data are available in the main text or the supplementary materials.

## SUPPLEMENTARY MATERIALS

Materials and Methods Figs. S1 to S37

References and notes (*1-83*)

Tables S1 to S26 Datasets S1 to S3

