## Supporting Information for "A virus co-opts vector’s m^6^A machinery for adaptive compensation"

*Running title：Virus as a cooperative partner*

Buli Fu[1,2,3,§], Jinyu Hu[1,§], Qun Liu[4,§], Rong Zhang[1,§], Xiaojie Wu[1,§], Xinfeng Guo[1], Shaonan Liu[1], Mingjiao Huang[1], Peipan Gong[3], Xuegao Wei[1], Jing Yang[1], Qimei Tan[1,7], Jinjin Liang[1], Chao He[1], Xuguo Zhou[5], Ralf Nauen[6], Xiaobin Shi[7], Youjun Zhang[1], Chris Bass[8]*, Xin Yang[1]*

***To whom correspondence may be addressed.**

 (C.B) or (X.Y)

#### This file includes:

Materials and Methods

Figs. S1 to S37

Tables S1 to S26

References (*1-83*)

#### Other Supplementary Material for this manuscript includes the following:

Dataset S1

Dataset S2

Dataset S3

### Materials and Methods

**RNA extraction and cDNA preparation.** Total RNA of samples was isolated using standard TRIzol reagent (Invitrogen). After detection of the RNA concentration with a spectrophotometer (NanoDrop 2000c, USA), the extracted RNA sample was then converted to cDNA using the PrimeScript RT reagent Kit (Takara) and the PrimeScript II 1st strand cDNA Synthesis Kit (Takara) according to the provided manuals.

**Quantitative real-time PCR (qPCR).** qPCR analysis was performed to detect the transcript levels of whitefly genes of interest using the Power SYBR Green PCR Master Mix Kit (Tiangen). qPCR analysis was performed in a QuantStudio 3 Real-Time PCR System (Applied Biosystems) via a two-step protocol as follows: an initial denaturation at 95°C for 1 min, followed by 40 cycles of denaturation at 95°C for 5 s, and annealing and extension at 60°C for 15 s. The relative expression of target gene, normalized to two reference genes (EF1α and RPL29) was calculated by the $2^{-\Delta\Delta Ct}$ method (77). Each treatment contained three biological replicates with four technical replicates. The gene-specific primers (Table S9) of target genes used for qRT-PCR analysis were designed using the Primer 5.0 software (Primer Biosoft).

**Western blot.** Approximately 300 adult whiteflies for each treatment were prepared for extraction of proteins using the ProteinExt Mammalian Total Protein Extraction Kit (Beyotime, China) following the manufacturer's instructions. The protein concentration was quantified using the BCA Protein Assay Kit (Beyotime, China). Protein extracts supplemented with SDS-PAGE loading buffer (CWBIO) were boiled for 10 min in 100 °C water. Then, protein samples were separated with a 4-20% SDS-PAGE precast gel (GenScript Young PAGE, USA) and transferred to a PVDF membrane (Merck Millipore, Germany). Following blocking in the Block Buffer Kit (Beyotime, China), the membrane was incubated against the primary rabbit-polyclonal antibodies of the target genes at 4°C overnight, followed by incubation with goat anti-rabbit IgG-HRP (CWBIO, 1:5000) secondary antibody at room temperature for 1 h. The SuperSignal West Pico Chemiluminescent Substrate (Thermo Fisher, USA) was used to visualize the protein bands. Images were acquired by the Tanon-5200 Chemiluminescent Imaging System (Tanon). Information of antibodies were available in supporting information (Table S10).

**Immunofluorescence (IF) assays and Multi-SIM super-resolution microscopy.** As reported previously (33), IF assay was performed to visualize protein signal located in the oocytes of whitefly ovary. First, whitefly ovaries were dissected from female adults as needed under a Leica microscope (Leica, Germany). The ovary samples were then transferred into 75% alcohol for 30 min permeation, and the specimens were subsequently fixed in 4% paraformaldehyde (PFA) for 4 h at room temperature. After blocking in Immunol Staining Blocking Buffer Kit (Beyotime, China) at 4 °C for 5 h, the ovary specimens were incubated with the primary antibodies against Vg (1:400) proteins lasted at 4 °C overnight, followed by incubation against the secondary antibody Alexa Fluor® 555 goat anti-rabbit IgG (1:500) (Abcam) for 2 h at room temperature in a black box. The specimens were placed on an adhesion microslide and stained with DAPI (Abcam) at room temperature. Fluorescence images were acquired using a Multi-SIM (Multimodality Structured Illumination Microscopy) imaging system (Beijing NanoInsights-Tech Co., Ltd.) equipped with an Objective Plan-Apochromat 63x/1.40 Oil M27 (ZEISS), semiconductor lasers (405 nm) /solid-state lasers (488nm, 561nm, 640nm) and a sCMOS (Complementary Metal-Oxide-Semiconductor) camera (Photometrics Kinetix). SIM image stacks were reconstructed using SI-Recon 2.23.3 (NanoInsights) with the following settings: pixel size 30.6 nm; channel-specific optical transfer functions; Wiener filter constant 0.01 for 2D mode and 0.005 for 3D mode; discard Negative Intensities background. Then the reconstructed SIM image was denoised with total variation (TV) constraint.

**RNAi experiments.** dsRNAs were synthesized using the T7 RiboMAX$^{TM}$ Express RNAi System (Promega, WI, USA) according to the manufacturer's instructions. All primers used for producing dsRNA are listed in Table S11. dsRNAs were fed to newly-emerged adult whiteflies (1 day after eclosion). Following dsRNA treatment, only females were collected for RNA extraction and cDNA preparation. The relative expression of genes of interest was then examined using qPCR described as above. After identifying the RNAi interference efficiency, those females fed with dsRNAs were used for further investigation of female fecundity, ovarian development, qPCR and western blot as needed. The dsEGFP served as a non-specific negative control.

**5'UTR and 3'UTR RACE cloning.** Rapid Amplification of cDNA Ends (RACE) was performed to generate full-length cDNAs of 5'UTR and 3'UTR of *Vg* gene by using the SMARTer RACE 5'/3' Kit (Takara). After isolation and purity of total RNA, the whitefly RNA samples were then used to prepare the 5'- and 3'-RACE-Ready cDNA synthesis (the First-Strand cDNA Synthesis) according to the protocols of synthesis reactions. Subsequently, 5'-RACE and 3'-RACE PCR reactions were conducted to generate the 5' and 3' cDNA fragments. Reactions of 50 μL total volume were composed of 2.5 μL RACE-Ready cDNA samples, 5 μL Universal Primer A Mix (UPM), 1 μL gene-specific primers (GSPs) and 41.5 μL Matser Mix. RACE products were then characterized by NucleoSpin Gel and PCR Clean-Up Kit (Takara) to confirm the desired cDNA products, and sequenced by Sanger with the T7 priming sites closed to the 5'- and 3'-cloing sites to ensure complete coverage in the sequencing trace. All primers used in RACE were summarized in the supporting information (Table S12).

**Dot blot.** Total RNA of 200 adult *B. tabaci* MED was isolated using standard TRIzol reagent (Invitrogen). The total RNA samples were dissolved to subgroups of 40 and 10 ng/μL. Then, the RNA subgroups were denatured at 95 °C for 5 min and cooled on ice for 5 min. A total of 10 μL RNA subgroups were spotted on a PVDF membrane (Millipore), and dried at 90 °C for 20 minutes. Two membranes for each assay were used to detect the $m^6A$ modification level and total amount of input RNA. After UV cross-link for 120s at 2000 joules, one membrane was stained with 0.02% methylene blue (Solarbio), and another was blocked in the Block Buffer Kit (Beyotime, China) followed by incubation with the $m^6A$ primary antibody (1:10000, Synaptic Systems). Subsequently, the membrane was further incubated with HRP-conjugated goat anti-rabbit IgG (1:5000, CWBIO) secondary antibody at room temperature for 1 h. The SuperSignal West Pico Chemiluminescent Substrate (Thermo Fisher, USA) was used to visualize the dot blots. Images were acquired by the Tanon-5200 Chemiluminescent Imaging System (Tanon).

**UPLC-MS/MS analysis of $m^6A$ level in whitefly.** Total RNA of 200 adult *B. tabaci* MED was isolated using standard TRIzol reagent (Invitrogen). A total of 1μg RNA samples were heated at 100 °C for 5 min, and chilled on ice immediately for 5 min. The RNA samples were then digested to generate single nucleosides by adding 18 μL reaction buffer (pH 5.3) containing 25 mM sodium acetate (Sigma Aldrich), 2.5 mM zinc chloride (Sigma Aldrich), sodium chloride and 2 μL (1 U) nuclease P1 (Sigma Aldrich) at 50 °C for 2h, followed by adding 5 μL of 1M ammonium bicarbonate and 1 μL (1 U) of alkaline phosphatase (Sigma Aldrich) at 37 °C for 4 h. Subsequently, the samples were dephosphotylated by bacterial alkaline phosphatase (10 U) (Invitrogen) and heated at 100°C for 5 min. UPLC-MS/MS was used to examine $m^6A$/A ratio of the samples via a program as follows: solvent A was maintained at 99% for 0.5 min, 70% for 2.5 min, 0% for 4.1 min, and finally 99% for 1.9 min. The mobile phase A was water containing 0.1% formic acid (Fisher Chemicals), and mobile phase B was acetonitrile containing 0.1% formic acid (Fisher Chemicals). UPLC–MS/MS analysis was performed with a Waters ACQUITY UPLC I-Class/Xevo TQ-S Micro (Waters, MA, USA) equipped with a BEH C18 1.7 μm 2.1 × 50 mm column (Waters, MA, USA). The column and FTN sample manager temperatures were set at 20°C and 4°C, respectively. Data were analyzed by MassLynx V4.1 (Waters, MA, USA). The experimental UPLC–MS/MS parameters were summarized in Table S13.

**Identification of functional $m^6A$ sites of *Vg* gene.** According to the conserved $m^6A$ motifs DRACH (D = C/A/U/G; R = A/G; H = A/C/U), the SRAMP prediction server (http://www.cuilab.cn/sramp/) provides a useful tool to predict $m^6A$ modification sites on the RNA sequences of interests. The potential $m^6A$ motifs of *Vg* gene including 5’UTR, CDS and 3’UTR were predicted through SRAMP. All the predicted $m^6A$ sites (Table S14) and their corresponding mutants were then arranged into dual-luciferase reporter assays, UPLC-MS/MS analysis and MeRIP-qPCR assay to identify the putative $m^6A$ sites.

**MeRIP-qPCR.** The total RNA of female whiteflies was used to perform Methylated RNA Immunoprecipitation (MeRIP) assays according to the protocols of Magna MeRIP $m^6A$ kit (Merck Millipore). Briefly, a total of 50 μg RNA sample was sheared to 100 nt in length by the fragmentation buffer. After centrifugation of the sonicated lysate, 5% of the supernatant was removed as input. The RNA supernatant was then transferred into Magna ChIP Protein A/G Magnetic Beads, followed by incubation with 8 μg IgG/$m^6A$/METTL3/METTL14 /ALKBH4 antibody at 25 °C for 45 min. After purification, the protein-RNA complexes were introduced into 100 μL of 5X immunoprecipitation (IP) buffer containing 5 μL of RNase inhibitor and rotated overnight at 4 °C. The $m^6A$-modified RNAs were reverse transcribed and converted to cDNA using the PrimeScript RT Reagent Kit (TaKaRa) according to the manuals. Subsequently, qPCR was conducted to quantify the level of IgG/$m^6A$/METTL3/METTL14/ALKBH4 binding to *Vg* gene by using the specific primers (Table S15) encompassing the $m^6A$ motifs. The relative $m^6A$/METTL3/METTL14/ ALKBH4 enrichment of *Vg* gene in the susceptible and resistant populations was calculated by normalizing to a reference gene *EF1α*.

**Recombinant protein preparation.** The full-length METTL3, METTL14 and ALKBH4 of the whitefly *B. tabaci* MED were cloned into pET28a vector and induced with 0.25 mM IPTG for 24 h at 16 °C. All these recombinant proteins and their mutants (Table S20) were then expressed in *E. coli* strain BL21 cell (Tsingke, China) and purified using the His-tag Protein Purification Kit (Beyotime, China) according to the manufacture’s protocols. The recombinant proteins were then verified by a 4-20% SDS-PAGE precast gel (GenScript Young PAGE, USA), followed by InstanBlue Coomassie Protein Stain (Abcam) or western blots. These recombinant proteins were used in $m^6A$ detection by UPLC-MS/MS and EMSA assays as needed.

**Biochemical analysis of methylation or demethylation effect on *Vg* $m^6A$ sites.** All the single-stranded RNAs (ssRNAs) (Table S21) containing the putative *Vg* $m^6A$ motifs were prepared by gene synthesis (Tsingke, China). Methylation or demethylation effect was assayed as previously reported (58, 78), with minor modifications. The reaction mixtures (100 μL) consisted of 50 μM of the recombinant METTL3 or METTL14 proteins, 5 μM ssRNA, 5 mM SAM, 10 mM HEPES, 5 mM DTT, 50 mM NaCl, 1 mM $MgCl_2$, followed by incubation at 25 °C for 2 h and then heating at 95 °C for 5 min. DNA/RNA was digested with 2 µL (1 U) nuclease P1 (Sigma Aldrich) in 50 µL of buffer (20 mM NaOAc, 5 mM ZnCl2, 50 mM NaCl) for 2 h at 50 °C, followed by the addition of 5 µL (1 M) of fresh NH4HCO3 and 1 µL (1 U) of alkaline phosphatase (Sigma Aldrich). The mixture was incubated for an additional 4 h at 37°C. The mixture was subsequently diluted 100-fold with 80% methanol and m6A and A nucleosides quantified by UPLC–MS/MS as described above. Methanol was used as the mobile phase A and water as mobile phase B. For demethylation activity assays, the ssRNAs tagged with $m^6A$ on the A nucleotide were used for incubation with ALKBH4-recombinant proteins in 100 µL reaction mixtures for 3 h, prior to $m^6A$ detection by UPLC–MS/MS analysis.

**Cell lines and transfection.** The *Drosophila* S2 cells were maintained in Hyclone SFX-insect medium (Thermo Fisher) at 27 °C and continuously sub-cultured in fresh medium every 2 days. Prior to transfection, the S2 cells were seeded in a 24-well plate with 300 µL SFX-insect medium at 27 °C for 48h. All plasmids were diluted to a same concentration of 100 ng/µL, followed by transfecting into the S2 cells with Lipofectamine 2000 transfection reagent (Invitrogen). The transfection lipid was then transferred into the 300 µL SFX-insect medium containing S2 cells described as above. The transfection mixtures were maintained at 27 °C and used for dual-luciferase reporter assays.

**Dual-luciferase reporter assays of the function of $m^6A$ site in 3′UTR or 5'UTR.** The dual luciferase assays were used to validate the functional role of the $m^6A$ site of *Vg* gene in S2 cells. Wild-type (WT) fragment containing CDS and 3′UTR or 5'UTR of *Vg* gene and mutant-type (MU) fragment was made by replacing the adenosine bases (A) with cytosine (G). The fragments of 3'UTR were attached to the pmirGLO vector by InFusion HD Cloning Kit (TaKaRa) according to the instructions, while the 5'UTR-fragments were cloned into the pGL4.26 reporter plasmid containing a mini-promoter. The WT and MU plasmid (600 ng) together with the reference reporter plasmid pGL4.73 (100 ng) were co-transfected into the S2 cells using Lipofectamine 2000 described as above. After 48 h, the S2 cells were lysed, and the fluorescence activity was detected using the Dual-Glo Luciferase Assay System (Promega). The relative luciferase activity (Firefly luciferase activity/Renilla luciferase activity) of each construct was normalized to that of the control group. Each reporter assay was performed with at least three independent replicates.

**qPCR analysis of the function of $m^6A$ site in CDS.** The full-length wild-type coding sequence (CDS) of *Vg* gene and mutated versions (A mutant to G) of the CDS were cloned into the pAC5.1b/V5/His B expression vector (Invitrogen) using the In-Fusion HD Cloning Kit (TaKaRa) according to the instructions. The plasmids (600 ng) were then transfected into S2 cells using Lipofectamine 2000 described as above. Following transfection for 48 h, the S2 cells were harvested and extracted total RNA using TRIzol reagent (Invitrogen) for qRT-PCR analysis. The relative expression levels of *Vg* gene were calculated by normalizing to a reference gene RPL32 of *Drosophila melanogaster* with at least three independent replicates.

**SELECT assays for detection specific m$^6$A sites and m$^6$A abundance at single-base resolution.** As previously reported (56, 57), single-base elongation- and ligation-based qPCR amplification (SELECT) was performed to verify the specific m$^6$A sites of *Vg* gene in whitefly according to the protocols of Epi-SELECT$^{TM}$ m$^6$A site identification kit (EPI-BIOTEK, China). Briefly, a total of 17 μL reaction system contains 1.5 μg sample RNA, 0.5 μM Up Probe, 0.5 μM Down Probe, dNTP and 10X-Reaction Buffer. The reaction system was then annealed under a temperature gradient process: 90 °C for 1 min, 80 °C for 1 min, 70 °C for 1 min, 60 °C for 1 min, 50 °C for 1 min, 40 °C for 6 min, and finally a hold at 4°C. Subsequently, 3 μL of a mixture containing 0.3 μL SELECT$^{TM}$ DNA polymerase, 0.47 μL SELECT$^{TM}$ ligase, and 2.23 μL ATP was added to the reaction mixture for single-base elongation under a temperature process: 40 °C for 20 min, denatured at 80 °C for 20 min, and kept at 4 °C. The amount of final elongated and ligated products were quantified by RT-qPCR analysis using a 10 μL reaction system: 1 μL cDNA product, 5 μL ChamQ Universal SYBR qPCR Master Mix (Vazyme), 0.2 μL Select F primer (10 μM), 0.2 μL Select R primer (10 μM) and 3.6 μL RNase-free Water. For identification of a specific m$^6$A site, 'A site' denotes the m$^6$A sites, while 'N site' represents a non-modification site around 'A site' (-6 to +2) served as a control. Notably, a qPCR assay with gene specific primers covering each m$^6$A site (Table S15) was carried out to confirm equivalent amount of total RNA for the control and treatment sample, prior to SELECT-qPCR assay for comparing m$^6$A abundance. The SELECT probes used are listed in Table S22.

**ALKBH4-associated SELECT for detection of m$^6$A modification at single-base resolution.** The ALKBH4-associated SELECT assay (16) was conducted to detect m$^6$A modification on the functional m$^6$A sites of *Vg* gene in whitefly using an Epi-SELECT$^{TM}$ m$^6$A site identification (with FTO-assisted step) kit, with some modifications. A total of 50 μL demethylation reaction mix containing 2 μg total RNA, 20 μg recombinant ALKBH4 protein, 10 μL reaction buffer (5X), 1 μL RI and RNase-free Water was incubated at 37 °C for 30 minutes, inactivated at 95 °C for 5 minutes and hold at 4 °C. The reaction was stopped by adding 4 μL EDTA (0.1M). For the control group, EDTA was added before the demethylation reaction. Next, a total of 17 μL reaction mix containing 2 ug demethylated RNA was mixed with 1.6 μL Up Probe (1 μM), 1.6 μL Down Probe (1 μM) and 2 μL

Reaction buffer (10X) was incubated at 90 °C for 1 min, 80 °C for 1 min, 70 °C for 1 min, 60 °C for 1 min, 50 °C for 1 min, 40 °C for 6 min, and finally a hold at 4°C. Then, 0.3 μL SELECT$^{TM}$ DNA polymerase, 0.47 μL SELECT$^{TM}$ ligase, and 2.23 μL ATP was added to the 17 μL reaction mixture for single-base elongation under a temperature process: 40 °C for 20 min, denatured at 80 °C for 20 min, and kept at 4 °C. SELECT-qPCR analysis was performed to detect the amount of final elongated and ligated products through a 10 μL reaction system: 1 μL cDNA product, 5 μL ChamQ Universal SYBR qPCR Master Mix (Vazyme), 0.2 μL Select F primer (10 μM), 0.2 μL Select R primer (10 μM) and 3.6 μL RNase-free Water.

**mRNA stability assays.** The wild-type and mutant-type *Vg* fragments containing 5'UTR, CDS and 3'UTR were cloned into the pAC5.1b/V5/His B vector (Invitrogen) by using the In-Fusion HD Cloning Kit (TaKaRa) according to the instructions. The plasmids (600 ng) were transfected into S2 cells using Lipofectamine 2000 described as above. After 36 h, the transfection mixtures were treated with 10 mg. $L^{-1}$ actinomycin D (ActD, Sigma). Following 30 min of incubation, the S2 cells were collected with an interval of 2 h and extracted total RNA using TRlzol reagent (Invitrogen). The cDNA was prepared using the PrimeScript RT reagent Kit (TaKaRa), and the relative expression levels of *Vg* were detected by qPCR analysis normalizing to a reference gene RPL32 of *Drosophila melanogaster* with at least three independent replicates.

**Electrophoretic mobility shift assay (EMSA).** EMSA were used for detecting direct binding of METTL3/METTL14/ALKBH4 to the $m^6A$ motifs using the Electrophoretic Mobility Shift Assay Kit (Invitrogen) according to the manufacture's introductions. First, oligonucleotide probes (Table S23) for the wide-type or mutant $m^6A$ motifs that were labeled with biotin at the 5'-terminus and with CY5 at 3'-terminus were prepared by gene synthesis (Tsingke, China). The ssRNA probes and recombinant proteins were incubated together in the EMSA/GEL-Shift binding buffer for 60 min at 25 °C. The RNA-protein complex was then electro-transferred by polyacrylamide gel electropheresis (PAGE). The PAGE gel was then immediately placed into the Tanon 5200Multi Chemiluminescent Imaging System (Tanon, China), and images were acquired with a Fluorescence Imaging System by laser trigger of CY5. Subsequently, the PAGE gel containing the RNA-protein mixture was then electro-transferred into a nylon membrane (Beyotime, China). After conjugate-blocking and substrate equilibration, the nylon membrane was imaged using the Tanon-5200 Chemiluminescent Imaging System (Tanon, China). Regarding the binding of ALKBH4 to the $m^6A$ motifs, note that the ssRNA probes tagged with $m^6A$ modification were used.

**Immunoprecipitation (IP) assays.** The whitefly protein from VR1 population was extracted with the ProteinExt Mammalian Total Protein Extraction Kit (Beyotime, China) following the manufacturer's protocols. IP assays were then performed to examine the direct binding interaction of interested proteins in the whitefly using the Immunoprecipitation Kit (BEAVER Biomedical Engineering, China). Specifically, equal total protein amounts extracted from female whiteflies were incubated with protein A/G-magnetic beads at 25 °C for 4 h, followed by incubation with the corresponding protein antibodies overnight at 4 °C. The beads were neutralized by washing two times with lysis buffer and pooled with eluate. The interaction of target proteins were detected by western blot analysis described as above. The IgG antibody was used as a negative control.

**Y2H assays.** The Y2H assays were performed to test the protein-protein interactions *in vitro*. The coding sequence of TYLCV structural protein CP, C2, C3, C4, V2 and Rep were separately attached to the pGADT7(AD) vector to produce the prey constructs. Meanwhile, the coding sequence of the whitefly METTL14 and ALKBH4 were separately cloned into pGBKT7 (BD) vector to produce the bait constructs. Various construct combinations (the bait and prey constructs) were co-transformed into yeast strain Y2HGold. Subsequently, the co-transformed yeast clones were serially diluted (1:10, 1:100 and 1:1000) and spotted on the selective medium (lacking Ade, His, Leu and Trp) for growth. Images were captured on day 5 following incubation at 30 °C. pGADT7-T and pGBKT7-53 were used as a positive control, and pGBKT7-lam and pGADT7-T were served as a negative control. Following confirmation of binding status of METTL14-C2 and ALKBH4-CP interaction by molecular docking, the binding residues was mutated for each protein and were made to produce the bait or prey constructs. Further Y2H assays were then conducted to verify the binding interaction of METTL14-C2 and ALKBH4-CP described above.

**GST-pull down assays.** GST pull-down assays were conducted to detect the direct interaction of target proteins of interest *in vitro* using the Pierce$^{TM}$ GST Protein Interaction Pull-Down Kit (Thermo Scientific). Briefly, the whitefly METTL14/ALKBH4 gene was separately cloned into pET28a vector and induced with 0.25 mM IPTG for 24 h at 16 °C, followed by purification as His-tagged fusion protein (the prey protein). Meanwhile, the TYLCV-CP/C2 gene was cloned into the pGEX-4T-1 vector and induced with 1 mM IPTG for 24 h at 20 °C, followed by purification as GST-tagged fusion protein (the bait protein). The bait TYLCV-CP/C2 protein were then immobilized on an equilibrated Glutathione Agarose Resin at 4 °C for 6 h with gentle rocking motion on a rotating platform. After washing with TBS containing Pull Down lysis buffer, the immobilized GST-tagged bait protein was incubated with the prey protein at 4 °C overnight with gentle rocking motion on a rotating platform. Following prey protein capture, the bait-prey complex was prepared for elution with 10 mM Glutathione Elution Buffer. Then, the protein samples were analyzed by western blot using the anti-6X His-Tag antibody (Abcam) and anti-6X GST-Tag antibody (Abcam) as needed. Protein-protein interaction was visualized using the SuperSignal West Pico Chemiluminescent Substrate (Thermo Fisher). Images were acquired by the Tanon-5200 Chemiluminescent Imaging System (Tanon).

**AlphaFold3 prediction and molecular docking.** The full-length amino acid sequence of METTL14, ALKBH4, TYLCV-CP and TYLCV-C2 was used for protein structure prediction by AlphaFold3. The single strain RNA (ssRNA) from *Vg* gene was also used for RNA structure prediction by AlphaFold3. Parameters -m is model 1, model 2, model 3, model 4, model 5, and -g False. The models with the highest confidence level 'ranked_0.pdb' were selected as the predicted structure. The predicted structure was then subjected to energy optimization using the Rosetta Relax module, and the resulting structure was used for molecular docking. Briefly, rigid docking was first performed using HDOCK to adjust the initial conformation of protein-protein/RNA complexes. Upon the results of rigid docking, flexible docking was accomplished by RosettaDock. The visualization of the docked complexes was accomplished through PyMOL version 2.1. The top-ranked model was output for subsequent molecular dynamics simulations to further investigate the binding behavior of protein-protein complexes.

**Molecular dynamics (MD) simulations.** The docked complexes obtained from docking calculations were used for molecular dynamics simulations. Molecular dynamics simulations were performed using Gromacs 2023.3 software. The solvent was predefined using the TIP3P parameters. The solution simulated was a 0.145 M NaCl solution. NaCl was used to neutralize the charge of the system at a distance of 10 Å. The simulations were carried out from 0k to 310.15K. The simulations were performed using Desmond's standard NPT relaxation protocol. A total of three sets of 50 ns computational simulations were carried out. Throughout the simulation, all involved hydrogen bonds were constrained using the LINCS algorithm with a 2 fs integration time step. Electrostatic interactions were calculated through the Particle-mesh Ewald (PME) method with a cutoff value of 1.2 nm. The cutoff for nonbonded interactions was set to 10 Å and updated every 10 simulation steps. Trajectory data were periodically recorded for subsequent analysis, including root-mean-square deviation (RMSD), root-mean-square fluctuation (RMSF), radius of gyration (Rg), and hydrogen bond count. Finally, binding free energies between the protein and small molecule ligands were calculated using the MM/GBSA module.

**Chemical cross-linking and mass spectrometry (XL-MS) analysis.** The XL-MS analysis was performed to characterize the intermolecular and intramolecular interactions of protein complex. Purified METTL14-C2 or ALKBH4-CP complex was incubated in reaction buffer and cross-linked with 0.5 mM DSS at room temperature for 60 min, with continuous shaking at 500 rpm for 1 h using a Thermo Mixer. The reaction was quenched with 20 mM ammonium bicarbonate (Sigma). The cross-linked proteins were precipitated with pre-cooled acetone and vacuum-dried. The resulting pellet was resuspended in denaturing buffer (8 M urea, 100 mM Tris-HCl, pH 8.5), followed by TCEP reduction and iodoacetamide (Sigma)-mediated cysteine alkylation. Proteins were then digested overnight with trypsin (Promega) at 37 °C at a protein-to-enzyme ratio of 50:1 (w/w). Tryptic peptides were then desalted using Pierce C18 spin columns (GL Sciences) and eluted with methanol-formic acid buffer (99.9% methanol, 0.1% formic acid). Eluted peptides were vacuum-dried, reconstituted in solvent A (0.1% formic acid in water), and analyzed via nanoLC-tandem MS. LC separation was conducted on an EASY-nLC 1200 UPLC system coupled to an Orbitrap Exploris 480 mass spectrometer (Thermo Fisher Scientific, Bremen, Germany) equipped with a nano-electrospray ionization source. A 2 µL peptide aliquot was loaded onto a C18 trap column (100 µm × 2 cm, Thermo Scientific Acclaim PepMap) at 10 µL/min for 3 min, then separated on a C18 analytical column (75 µm × 50 cm) using a segmented linear gradient of solvent B (80% ACN, 0.1% formic acid): 5–35% over 108 min, 30–50% over 6 min, ramped to 100% within 1 min, and maintained at 100% for 5 min. The column was re-equilibrated for 10 min after each run. Chromatography was performed at a constant flow rate of 300 nL/min and column temperature of 60 °C, with an electrospray ionization voltage of 2.3 kV. The mass spectrometer was operated in data-dependent acquisition (DDA) mode. Full MS scans (m/z 350–1800) were acquired at 60 K resolution, followed by 15 sequential HCD MS/MS scans at 15 K resolution. The MS AGC target was set to 1 × 10 with a 50 ms maximum injection time, while MS/MS parameters included an AGC target of 1 × 10, intensity threshold of 13,000, and 80 ms maximum injection time. Spectra were recorded with a 30 s dynamic exclusion window, a fixed MS/MS first mass of 110 m/z, and a normalized HCD collision energy of 30%. Raw MS data were analyzed using pLink v2.3.11 for cross-linked peptide identification, with searches performed against target and reversed decoy protein databases. The search parameters were defined as: trypsin digestion (≤3 missed cleavages), peptide mass of 600–6000 Da, peptide length of 6–60 amino acids, ±20 ppm precursor and fragment tolerances, DSS cross-linking, fixed cysteine carbamidomethylation, and variable methionine oxidation. PSM-level results were filtered at 5% FDR with a refined mass tolerance of ±10 ppm. Validated cross-linking data were finally visualized via the xiNET online platform (https://crosslinkviewer.org).

**Microscale thermophoresis (MST) assay.** The MST assays for the binding affinities of target proteins to nucleic acids were run on a MONOLITH NT.115 instrument (NanoTemper Technologies). In brief, recombinant protein of interest was served as target and labeled with fluorescent by Monolith Protein Labeling Kit RED-NHS (MO-L011, NanoTemper Technologies). The substrate ssRNAs (labeled with CY5) were employed as the ligands (Table s23). For each assay, the labelled protein (20 nM) was incubated with the varying concentrations (100 μM to 0.00305 μM) of the ligand in the MST assay buffer (10 mmol/L Tris-HCl; 150 mmol/L NaCl; 0.05% Tween 20; pH=7.8-8.2) at room temperature for 15 min. All samples were loaded into MST NT.115 standard glass capillaries and measurement was performed at 20% excitation power to control the fluorescence value between 300 and 400 using the MO control software (v.1.6.1). The thermophoresis time (t) was 23 s, and the experimental temperature (T) was 25 °C with 40% MST power (medium). At least three independent experiments were repeated and then the dissociation constant ($K_D$) values were calculated using MO Affinity Analysis software (v.2.3) of NanoTemper. The cold and hot fluorescence were measured at −1–0 s and 4–5 s to avoid thermally induced protein configuration change, respectively.

**Surface plasmon resonance (SPR) analysis.** SPR analysis was conducted with a Biacore 1K system (Cytiva) to determine the binding affinities. The Series S Sensor Chip CM5 was firstly installed on the SPR instrument in accordance with the standard procedure. Next, the ligand protein (METTL14 or ALKBH4) was immobilized to the CM5 Chip via covalent bonds to the amino acid residues in immobilization buffer. Subsequently, different concentrations of viral protein (C2 or CP) and the substrate ssRNAs (labeled with CY5, Table S23) were together diluted in the analyte buffer and were injected into the flowing channel to allow their interaction with the ligand protein. The interacting phase included 120 s of association phase and 300 s of dissociation phase. The kinetic parameters of the binding reactions were calculated and analyzed by using Biacore 1K Evaluation Software (Cytiva).

**Virus-induced gene silencing (VIGS) experiment.** VIGS assays were conducted to examine the effect of continuous interference with the *METTL14* gene on reproductive performance of the whitefly. The experiment protocol has previously been described (46), with slight modifications. First, a 332 bp fragment of *METTL14* and a 334 bp fragment of EGFP were separately cloned from *B. tabaci* MED using specific primers, and then the PCR product was then cloned into XbaI-SacI-cut pTRV2 to construct pTRV2-METTL14 and pTRV2-EGFP. The pTRV1, pTRV2-METTL14 and pTRV2-EGFP vectors were then transferred into *Agrobacterium tumefaciens* GV3101 by electroporation, and the bacteria were selected on LB agar plates containing 100 mg/ml of rifampicin and 50 mg/ml of kanamycin. The *A. tumefaciens* carrying pTRV1 and target-gene pTRV2 were grown in 5 mL LB medium with 100 mg/L rifampicin and 50 mg/L kanamycin for 18 h at 30 °C with shaking at 200 rpm. Two milliliters of starter culture was diluted into 48 mL identical LB medium supplemented with 200 μM acetosyringone and incubated for an additional 18 h under matching shaking and temperature conditions. Cells were harvested by centrifugation (3000×g, 10 min), washed once with infiltration buffer (10 mM $MgCl_2$, 10 mM MES, 200 μM acetosyringone), and resuspended in fresh buffer to an $OD_{600}$ of 0.4. pTRV1 and pTRV2 bacterial suspensions were pooled at a 1:1 volume ratio, then infiltrated into the two largest mature true leaves of tobacco plants (*Nicotiana benthamiana*) using a 1 mL needleless syringe. Treated plants were covered and incubated overnight following infiltration and kept in an intelligent greenhouse at 25 °C for 20 days. The successfully infected tobacco plants (Fig. S33A, B) were used for further investigations. To characterize VIGS-mediated effects on *B. tabaci* reproductive fitness, a clip cage was secured to a single leaf of each of 12 METTL14-VIGS silenced tobacco plants; 10 newly emerged (<24 h) female whiteflies were confined per cage. Every 7 days, egg numbers on caged leaves were counted. Twelve EGFP-VIGS plants served as negative controls (Fig. S33A, B).

**Insect vector, virus and plants.** From 2023 to 2025, a series of field vectors *B. tabaci* MED were sampled on numerous agricultural crops across China (Table S1). Each of these field population was used for detecting neonicotinoids resistance level and TYLCV infection rate (Tables S2 and S3). The laboratory source of the whitefly *B. tabaci* MED was described in our previous work (33). These nonviruliferous samples comprised two susceptible strain ($S^{\#1}$, $S^{\#2}$) and two resistant strains ($R^{\#1}$, $R^{\#2}$), which were reared with cotton plants. In 2023, a total of 400 individuals (mixed sex) from each of the four strains were separately transferred to healthy tomato plants and TYLCV-infected tomato plants to prepare healthy strains (S1, S2, R1, R2) and viruliferous strains (VS1, VS2, VR1, VR2), respectively. Further details on these strains are provided in the supporting information (Table S4). All strains were kept in nylon cages and maintained in an intelligent-controlled greenhouse at 25 °C and 70% relative humidity (RH) under a 14:10 (L:D) photoperiod. To maintain the level of insecticide resistance, the resistant strains were arranged into imidacloprid and thiamethoxam selection at a concentration of 100 mg/L through root-irrigation of plants with an interval of 45 days. Infectious clones of TYLCV isolate SH2 (GenBank accession no. AM282874) were agro-inoculated into true leaf stage tomato seedlings. TYLCV-infected tomato plants could be used on 20 days following virus inoculation. All plants were grown in insect-proof greenhouses under a controlled temperature of 25 °C and natural lighting.

**Insecticide bioassays.** Four commonly used neonicotinoids (Table S5) were used in insecticide bioassays on adult of *B. tabaci* MED as previously reported (33). Specifically, each insecticide bioassay included seven concentrations and contained four replicates with approximately 20 individual whiteflies for each replicate. After checking the mortality for each treatment, bioassay data were used to determine the lethal concentration required to kill 50% of the population ($LC_{50}$) and associated 95% confidence limits (95% CL) using the POLO program PC PoloPlus (LeOra Software, Berkeley, CA, 2003). The resistance ratio (RR) was measured by dividing the $LC_{50}$ value of a strain by the corresponding $LC_{50}$ value of the susceptible reference strain.

**'Age stage-two sex' life table study**. The 'age stage-two sex' life table study (79) for the whitefly *B. tabaci* MED has been reported in our previous works (80). For life-history cycle and biological traits of *B. tabaci* MED has been characterized in our previous works (79-81). Briefly, at least 200 eggs for each whitefly population were reared on various tomato plants with approximately 10 eggs remained on a tomato leaf. The eggs were identified as creamy and ovoid and gradually turned dark brown over time. Once hatched, the feeding point of each nymph was marked with a fine-tip nontoxic Sharpie marker. The 1st-(N1), 2nd-(N2), 3rd-(N3) and 4th-instar (N4) nymph were distinguished depending on their relative body size after a molt compared to that in the previous stage. The developmental time and survival at each immature stage for all marked individuals were recorded daily. After eclosion, newly emerged male and female adults were collected daily. A pair of male and female was then transferred into a 5 mL centrifuge tube to allow mating and reared with a piece of tomato leaf as food and egg laying. The food for each treatment was replaced every day. The number of eggs laid and adult survival was recorded every day until all adults died.

**Evaluation of life-table parameters and relative fitness.** Biological raw data on fitness parameters were analyzed via the TWO-SEX-MS program (82). The age-stage-specific survival rate (*sxj*) (where *x* = age and *j* = stage), age-specific survival rate (*lx*), age-specific fecundity (*mx*) and life-table parameters (*r*, the intrinsic rate of increase; *λ*, the finite rate of increase; $R_0$, the net reproduction rate; and *T*, the mean generation time) were calculated accordingly. The standard errors (SEs) of all life table parameters, including *r*, *λ*, $R_0$, *T*, adult longevity, and fecundity, were estimated by a bootstrapping procedure with 100,000 bootstraps. A paired bootstrap test was used to detect differences among strains based on the confidence interval of the differences. The relative fitness (*Rf*) values were calculated by *r* and $R_0$ (33, 80, 81).

**Fecundity measurement.** Newly emerged (1-day-old) females and males of *B. tabaci* MED were collected before experiments. Female fecundity of *B. tabaci* MED was assessed by counting eggs on cotton or tomato leaf as described previously (33). Briefly, a single female was introduced into a centrifuge tube (5 mL) for rearing with a piece of fresh plant leaf. The number of eggs laid by the female was counted daily under a microscope at 20X magnification, and the plant leaf was refreshed every day. A total of 80 females arranged into four groups (replicates) with 20 females for each replicate were measured. Additionally, a 'cage experiment' was also carried out to investigate female fecundity on alive plants. Approximately five females and five males were transferred into a small clip, which was then clamped on a leaf of plant seedling. After removal of the alive females and males to a new leaf, the number of total eggs laid on the previous leaf was recorded every seven days within 30 days. Female fecundity was divided into early (1-7 days), middle (8-14 days) and late (15-21 days) stage by the female age. This experiment contained at least 12 replicates (clips) for each treatment.

**Light microscopy.** As reported in our previous work (33), ovarian development of *B. tabaci* MED was determined by the number of matured oocytes contained in the female ovary. After dissection of the ovary, the number of matured oocytes was counted and photographed at 60X magnification using a Leica microscope (Leica, Germany). Images were acquired at 120X magnification with a Leica Application Suite X. A total of 80 females arranged into four groups (replicates) with 20 females for each replicate were measured.

**Statistical analysis and data visualization.** Unless otherwise stated, all quantitative data are presented as the mean ± SEM of at least three independent experiments. The statistical significance of differences among treatments was determined using Student's *t*-test or one-way analysis of variance (ANOVA) with Tukey's HSD test by IBM SPSS Statistics 23.0. All graphs were created through GraphPad Prism 8.3, SigmaPlot 12.5, Adobe Illustrator 2022 and Microsoft Office 2024.

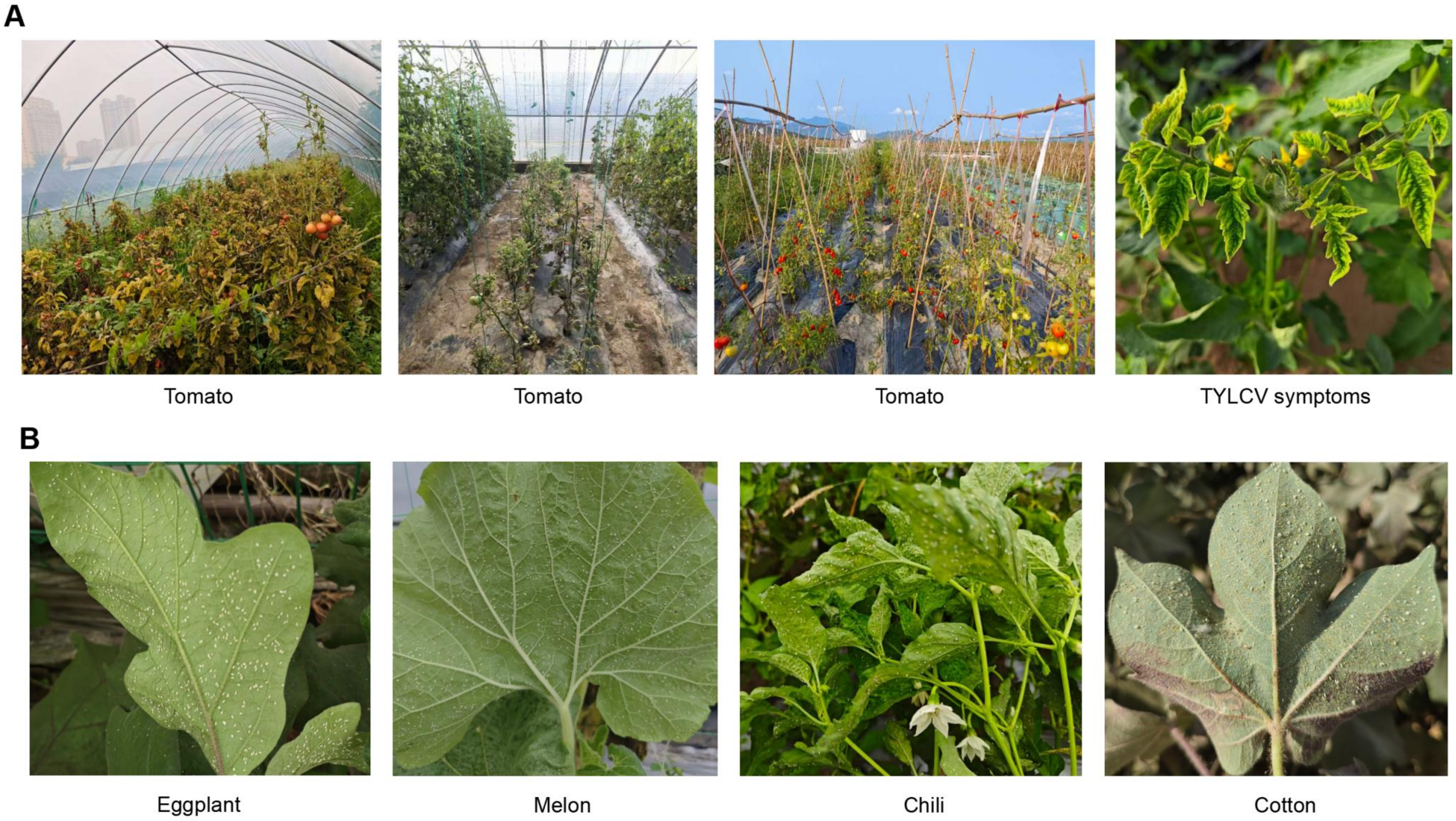
A
Tomato
Tomato
Tomato
TYLCV symptoms
B
Eggplant
Melon
Chili
Cotton

**Fig. S1 Outbreaks of TYLCV and its vector *B. tabaci* on agricultural crops across China.** (**A**) Representative images showing current outbreaks of TYLCV in various tomato plantations. (**B**) Representative images showing current outbreaks of *B. tabaci* on numerous agricultural crops.

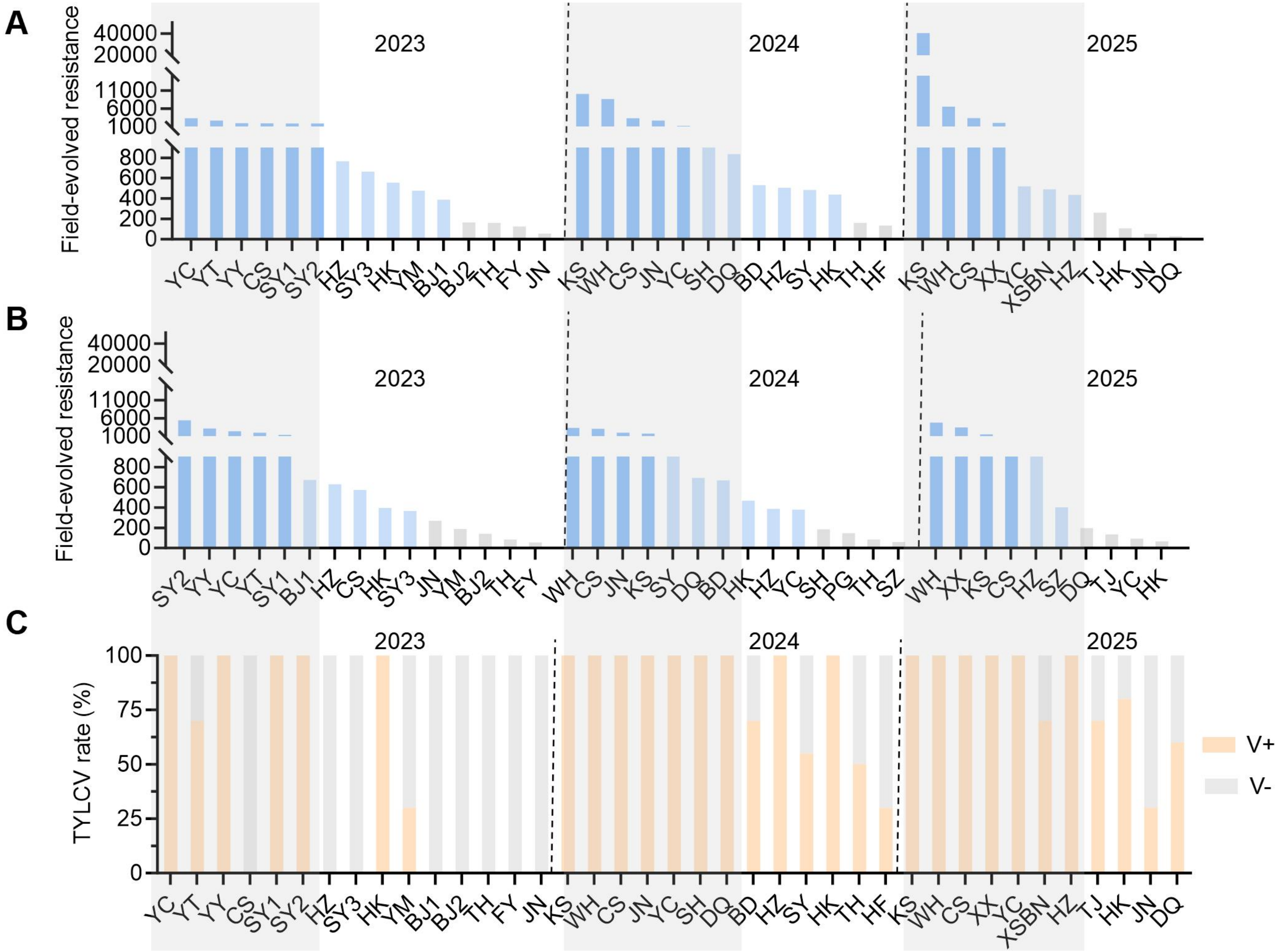

A
B
C
Field-evolved resistance
TYLCV rate (%)
2023
2024
2025
40000
20000
11000
6000
1000
800
600
400
200
0
100
75
50
25
YC YT YY CS SY1 SY2 HZ SY3 HK YM BJ1 BJ2 TH FY JN
KS WH CS JN YC SH DQ BD HZ SY HK TH HF
KS WH CS XX YC XSBN HZ TJ HK JN DQ
SY2 YY YC YT SY1 BJ1 HZ CS HK SY3 JN YM BJ2 TH FY
WH CS JN KS SY DQ BD HK HZ YC SH PG TH SZ
WH XX KS CS HZ SZ DQ TJ YC HK
V+
V-

**Fig. S2 Field-evolved neonicotinoid resistance and TYLCV infection in the vector *B. tabaci* MED.** (**A**, **B**) Resistance level of imidacloprid (**A**) and thiamethoxam (**B**) in filed vectors sampled from 2023 to 2025. (**C**) Infection rate of TYLCV in the field vectors from 2023 to 2025. For details see Table S1-S3.

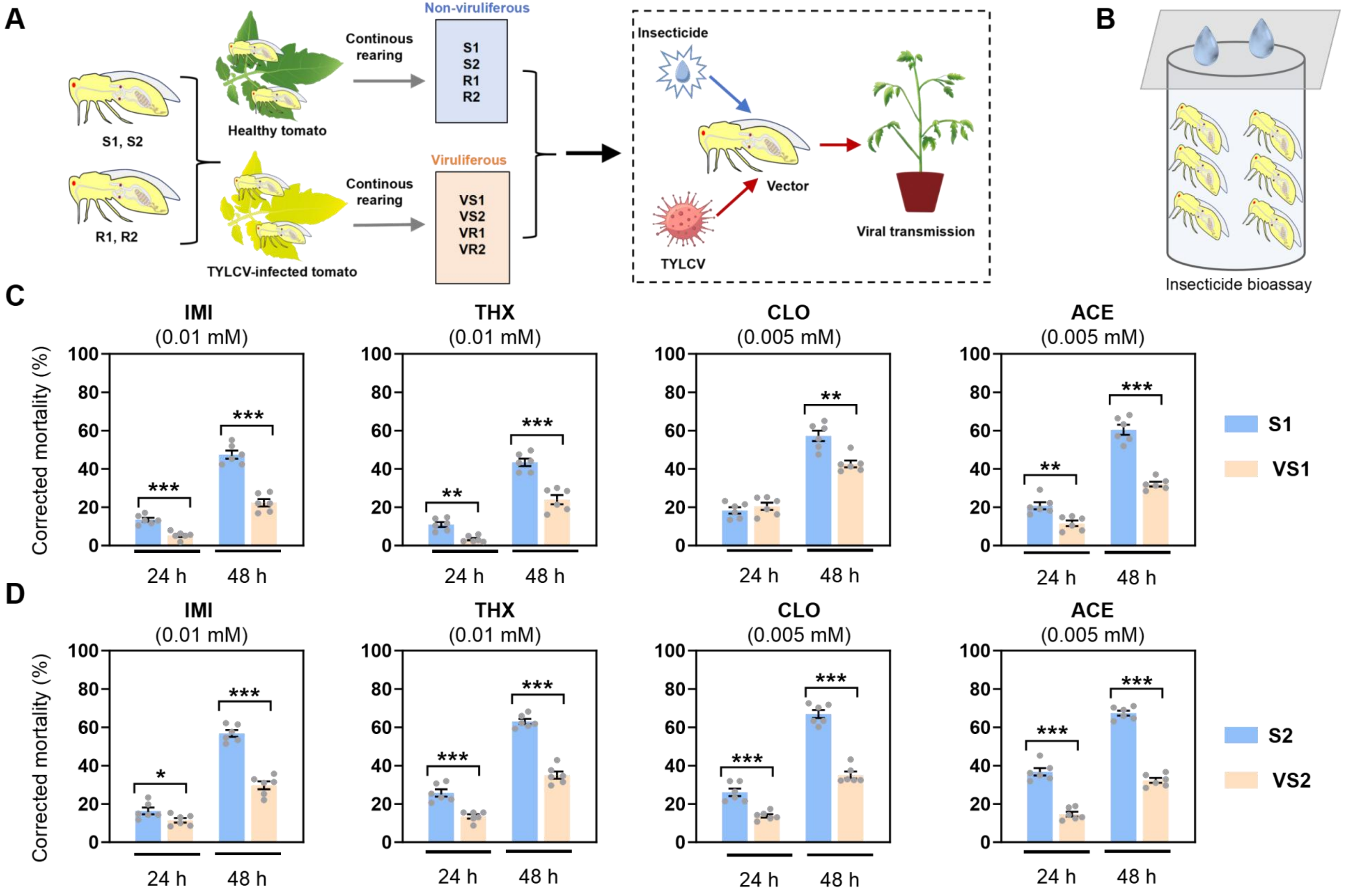
A
S1, S2
R1, R2
Healthy tomato
TYLCV-infected tomato
Continous rearing
Continous rearing
Non-viruliferous
S1
S2
R1
R2
Viruliferous
VS1
VS2
VR1
VR2
Insecticide
Vector
TYLCV
Viral transmission
B
Insecticide bioassay
C
IMI (0.01 mM)
THX (0.01 mM)
CLO (0.005 mM)
ACE (0.005 mM)
Corrected mortality (%)
24 h
48 h
S1
VS1
D
IMI (0.01 mM)
THX (0.01 mM)
CLO (0.005 mM)
ACE (0.005 mM)
Corrected mortality (%)
24 h
48 h
S2
VS2

**Fig. S3 TYLCV infection decreased neonicotinoids sensitivity of the susceptible vector *B. tabaci* MED.** (**A**) A flow diagram for screening stably heritable strains to test effect of TYLCV infection on insecticide resistance in the laboratory. (**B**) Schematic diagram of insecticide bioassay for adult whitefly. (**C**, **D**) Corrected mortality of the susceptible whitefly from viruliferous strain (VS1 and VS2) to neonicotinoids compared to their non-viruliferous counterparts (S1 and S2) (n = 7, Student's *t* test: *$P < 0.05$, **$P < 0.01$, ***$P < 0.001$). Neonicotinoid insecticides include imidacloprid (IMI), thiamethoxam (THX), clothianidin (CLO) and acetamiprid (ACE).

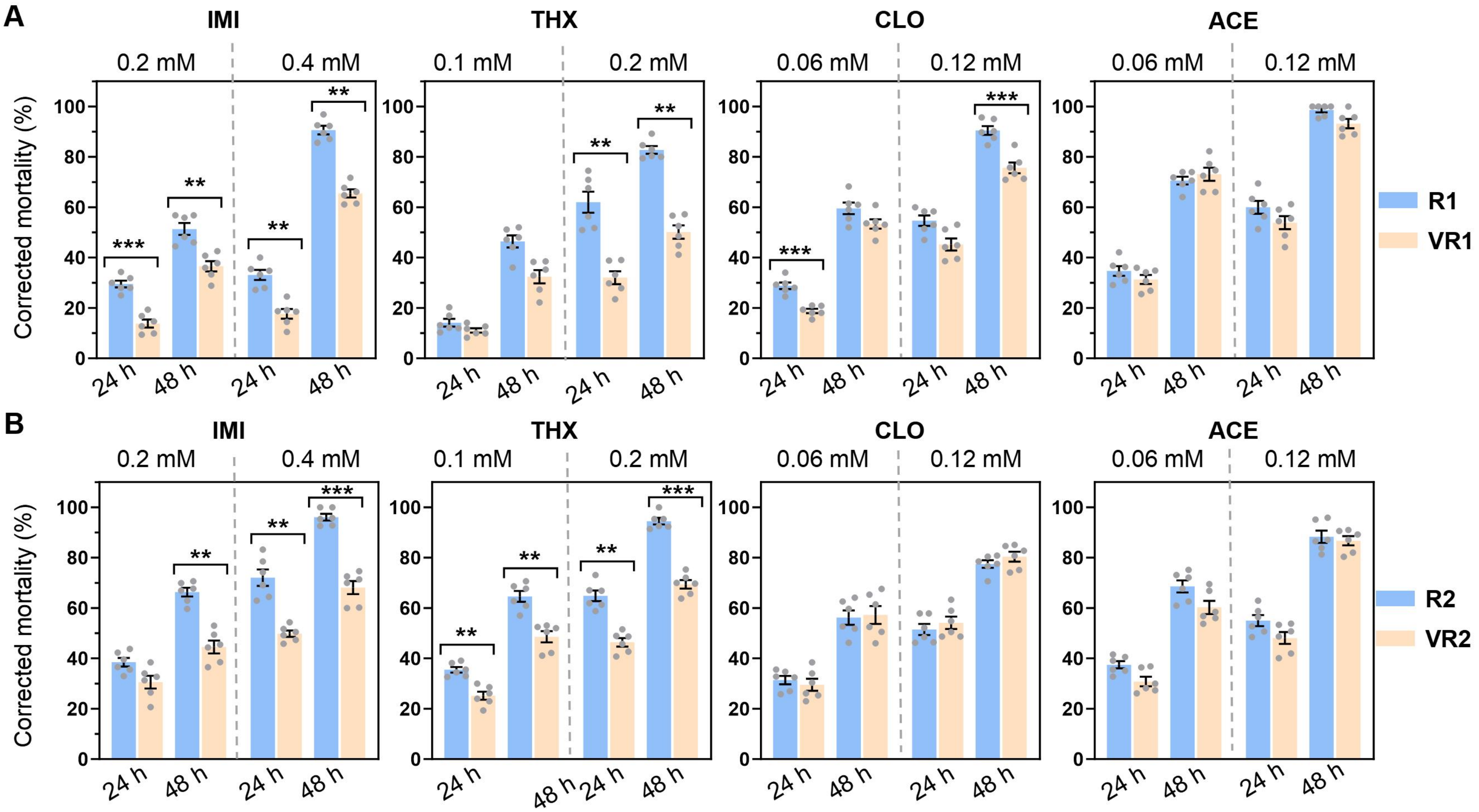

A
B
IMI
THX
CLO
ACE
0.2 mM
0.4 mM
0.1 mM
0.2 mM
0.06 mM
0.12 mM
Corrected mortality (%)
24 h
48 h
R1
VR1
R2
VR2

**Fig. S4 TYLCV infection enhance imidacloprid and thiamethoxam resistance in the resistant vector *B. tabaci* MED.** (**A**, **B**) Corrected mortality of the resistant whitefly from viruliferous strain (VR1 and VR2) to neonicotinoids compared to their non-viruliferous counterparts (S1 and S2) ($n = 7$, Student's *t* test: $^{*}P < 0.05$, $^{**}P < 0.01$, $^{***}P < 0.001$). Neonicotinoid insecticides include imidacloprid (IMI), thiamethoxam (THX), clothianidin (CLO) and acetamiprid (ACE).

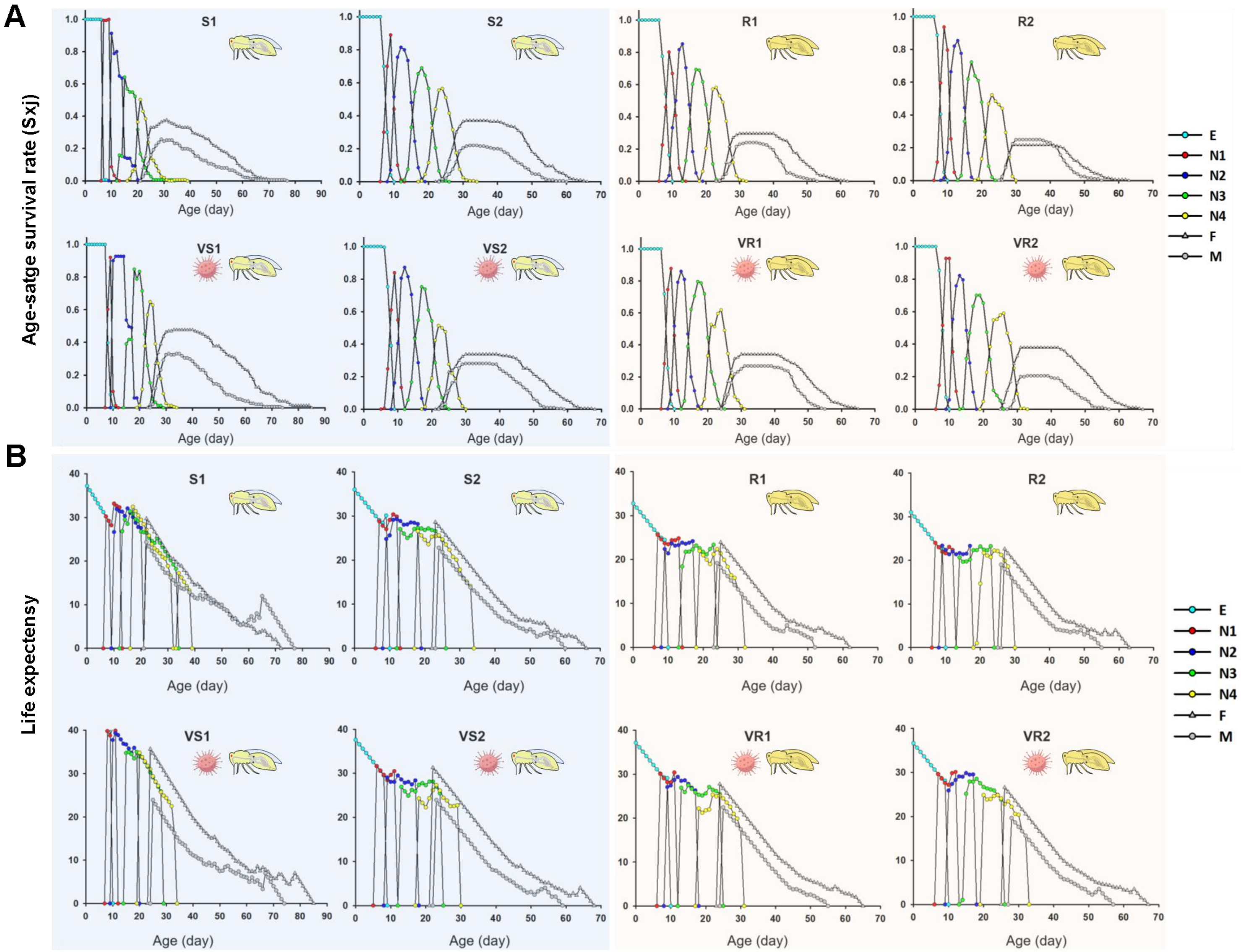
A
B
S1
S2
R1
R2
VS1
VS2
VR1
VR2
Age-satge survival rate (Sxj)
Life expectensy
Age (day)
E
N1
N2
N3
N4
F
M

**Fig. S5 Life-table analysis of survival rate and life expectancy of experimental strains of the vector *B. tabaci* MED.** (**A**) Age-stage survival rate (*Sxj*). (**B**) Life expectancy (*Exj*). The whitefly stages examined` include egg (E), 1st-insatr nymph (N1), 2nd-instar nymph (N2), 3rd-instar nymph (N3), 4th-instar nymph (N4), adult female (F) and adult male (M).

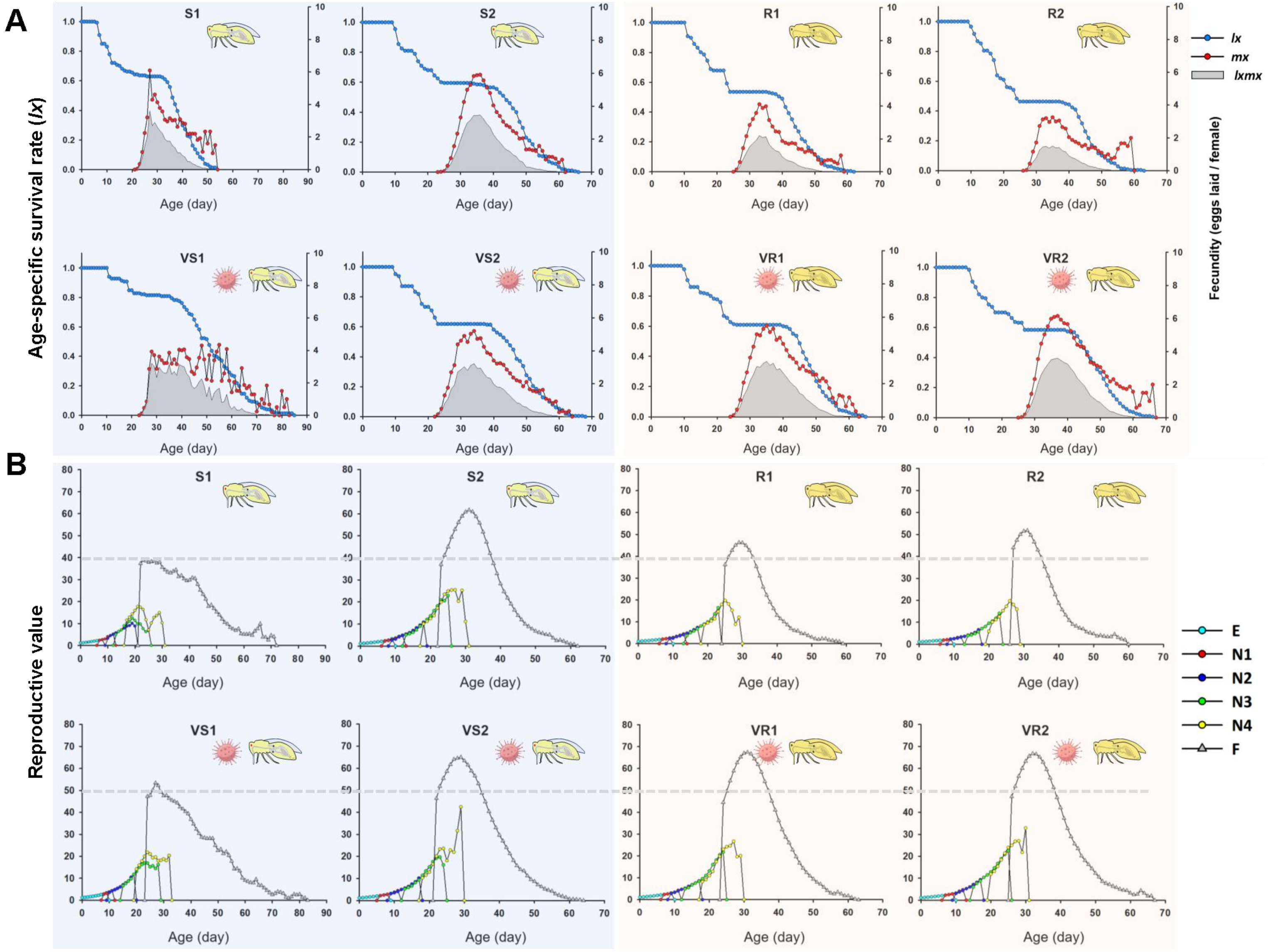
A
B
S1
S2
R1
R2
VS1
VS2
VR1
VR2
lx
mx
lxmx
Age-specific survival rate (lx)
Fecundity (eggs laid / female)
Age (day)
Reproductive value
E
N1
N2
N3
N4
F

**Fig. S6 Effect of TYLCV infection on reproductive capacity of experimental strains of the vector *B. tabaci* MED.** (**A**) Age-specific survival rate ($l_x$), fecundity ($m_x$) and net maternity ($l_x m_x$). (**B**) Reproductive value. The whitefly stages examined` include egg (E), 1st-insatr nymph (N1), 2nd-instar nymph (N2), 3rd-instar nymph (N3), 4th-instar nymph (N4), adult female (F) and adult male (M).

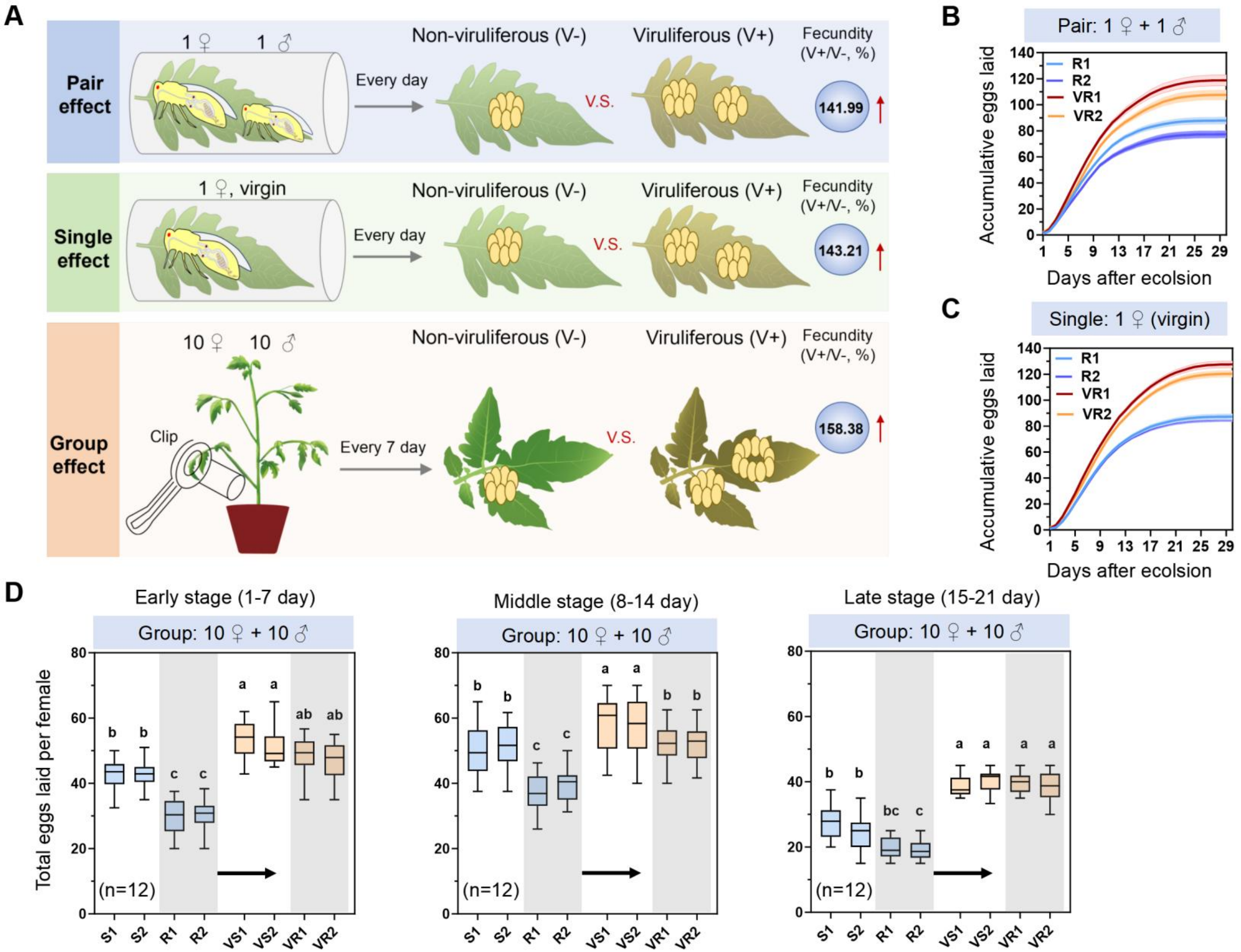

A
Pair effect
1 ♀ 1 ♂
Every day
Non-viruliferous (V-)
V.S.
Viruliferous (V+)
Fecundity (V+/V-, %)
141.99
Single effect
1 ♀, virgin
Every day
Non-viruliferous (V-)
V.S.
Viruliferous (V+)
Fecundity (V+/V-, %)
143.21
Group effect
10 ♀ 10 ♂
Clip
Every 7 day
Non-viruliferous (V-)
V.S.
Viruliferous (V+)
Fecundity (V+/V-, %)
158.38
B
Pair: 1 ♀ + 1 ♂
Accumulative eggs laid
R1
R2
VR1
VR2
Days after ecolsion
C
Single: 1 ♀ (virgin)
Accumulative eggs laid
R1
R2
VR1
VR2
Days after ecolsion
D
Early stage (1-7 day)
Group: 10 ♀ + 10 ♂
Middle stage (8-14 day)
Group: 10 ♀ + 10 ♂
Late stage (15-21 day)
Group: 10 ♀ + 10 ♂
Total eggs laid per female
(n=12)
S1 S2 R1 R2 VS1 VS2 VR1 VR2

**Fig. S7 Validation of reproductive fitness by determining pair, single and group effect on female fecundity in the vector *B. tabaci* MED.** (**A**) Schematic diagram for testing the effect of pair, single and group on female fecundity of the vector strains. (**B**-**D**) Investigation of pair (**B**), single (**C**) and group (**D**) effect on female fecundity of whitefly experimental strains (Tukey's HSD multiple comparisons test: $P < 0.05$).

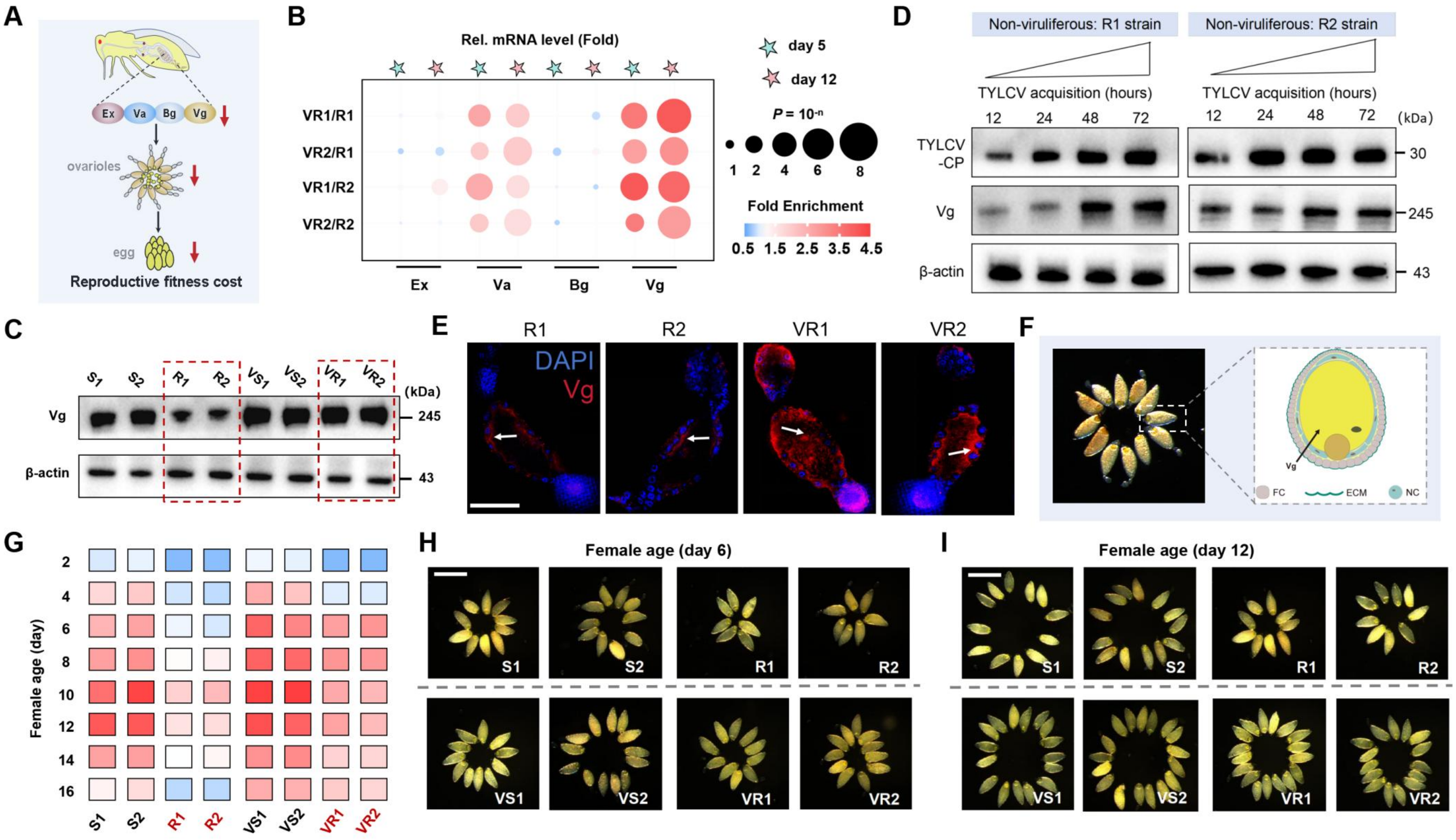

A
Ex
Va
Bg
Vg
ovarioles
egg
Reproductive fitness cost
B
Rel. mRNA level (Fold)
day 5
day 12
VR1/R1
VR2/R1
VR1/R2
VR2/R2
Ex
Va
Bg
Vg
P = 10-n
1 2 4 6 8
Fold Enrichment
0.5 1.5 2.5 3.5 4.5
C
S1 S2 R1 R2 VS1 VS2 VR1 VR2
(kDa)
Vg
245
β-actin
43
D
Non-viruliferous: R1 strain
Non-viruliferous: R2 strain
TYLCV acquisition (hours)
12 24 48 72
(kDa)
TYLCV -CP
30
Vg
245
β-actin
43
E
R1
R2
VR1
VR2
DAPI
Vg
F
FC
ECM
NC
G
Female age (day)
2 4 6 8 10 12 14 16
S1 S2 R1 R2 VS1 VS2 VR1 VR2
H
Female age (day 6)
S1 S2 R1 R2
VS1 VS2 VR1 VR2
I
Female age (day 12)
S1 S2 R1 R2
VS1 VS2 VR1 VR2

**Fig. S8 TYLCV-mediated vitellogein (Vg) overexpresssion enhances the reproductive fitness of neonicotinoids resistance in the whitefly *B. tabaci* MED.** (**A**) Schematic diagram showing whitefly oogenesis genes underlying reproductive cost with neonicotinoid resistance in our recently published work (33). (**B**) qRT-PCR analysis of the *Ex*, *Va*, *Bg*, and *Vg* mRNA level in female adults of the experimental strains sampled on days 5 and 12 after eclosion (n = 3, Student's *t* test). (**C**) Western blot analysis of the Vg protein level in female adults of the experimental strains sampled on day 10 after eclosion. (**D**) Vg protein level after TYLCV acquisition at various time points. (**E**) Immunofluorescence assays of Vg signal in female oocytes of the experimental strains. Scar bar, 50 μm. **F**) Schematic diagram for the role of Vg in maturing oocytes in whitefly. (**G**) The average number of mature oocytes of in the experimental strains after eclosion (n ≥ 60). (**H**, **I**) Representative images displaying mature oocytes of female whitefly sampled on day 6 (**H**) and 12 (**I**) after eclosion. Scale bars, 200 μm.

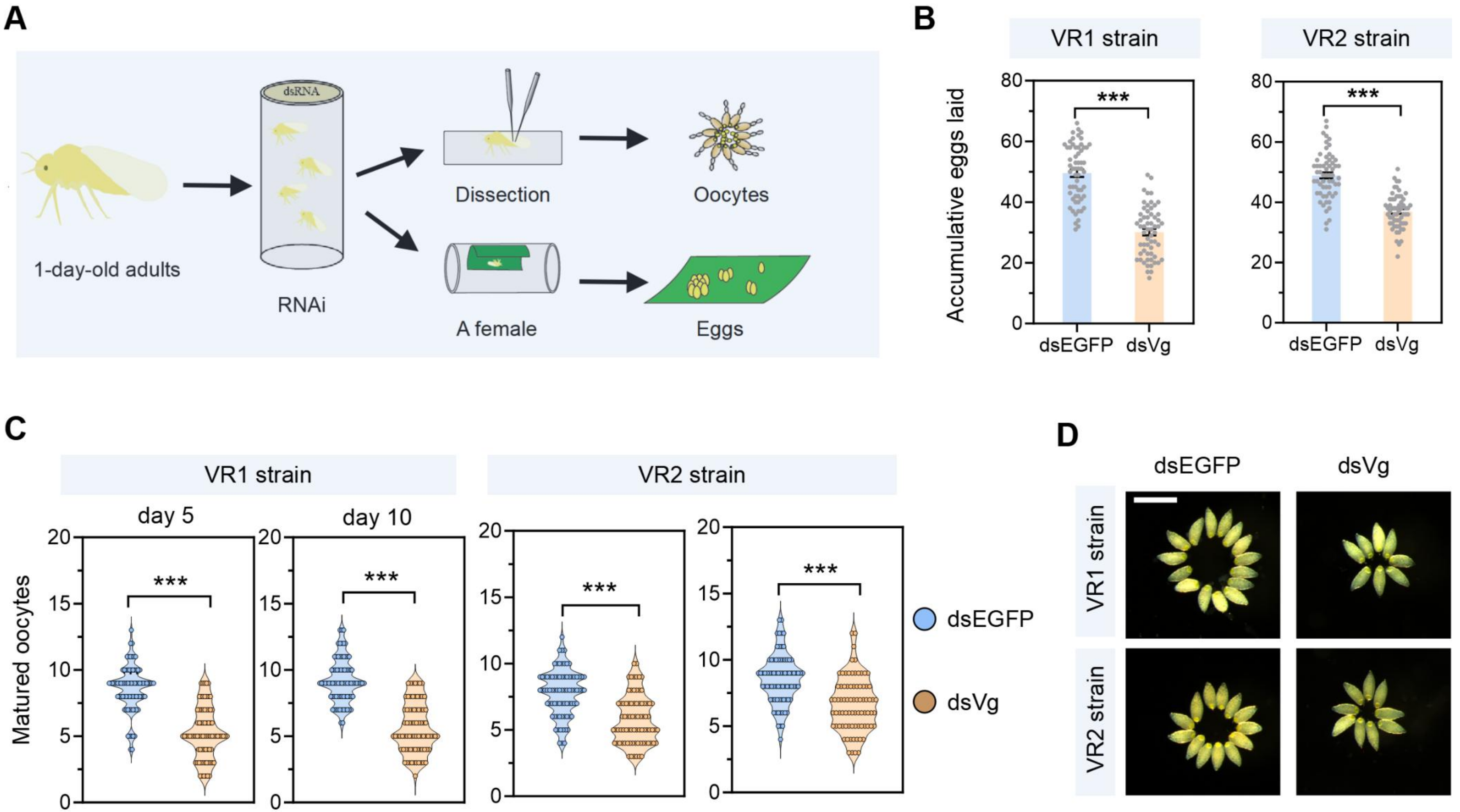
A
dsRNA
1-day-old adults
RNAi
Dissection
Oocytes
A female
Eggs
B
VR1 strain
VR2 strain
Accumulative eggs laid
***
dsEGFP
dsVg
C
VR1 strain
VR2 strain
day 5
day 10
Matured oocytes
***
dsEGFP
dsVg
D
dsEGFP
dsVg
VR1 strain
VR2 strain

**Fig. S9 RNAi-mediated validation of the role of Vg in conferring reproductive compensation in the neonicotinoids resistant vector *B. tabaci* MED.** (**A**) A schematic of the experimental process used for RNAi. (**B**, **C**) Functional study of the cumulative eggs laid (**B**) and the number of mature oocytes (**C**) per female after RNAi knockdown of *Vg* in viruliferous strains VR1 and VR2 (n ≥ 60, Student' s *t* test: ***$P$ < 0.001). (**D**) Representative images showing the mature oocytes in adult female sampled on day 10 after RNAi. Scar bar, 200 μm.

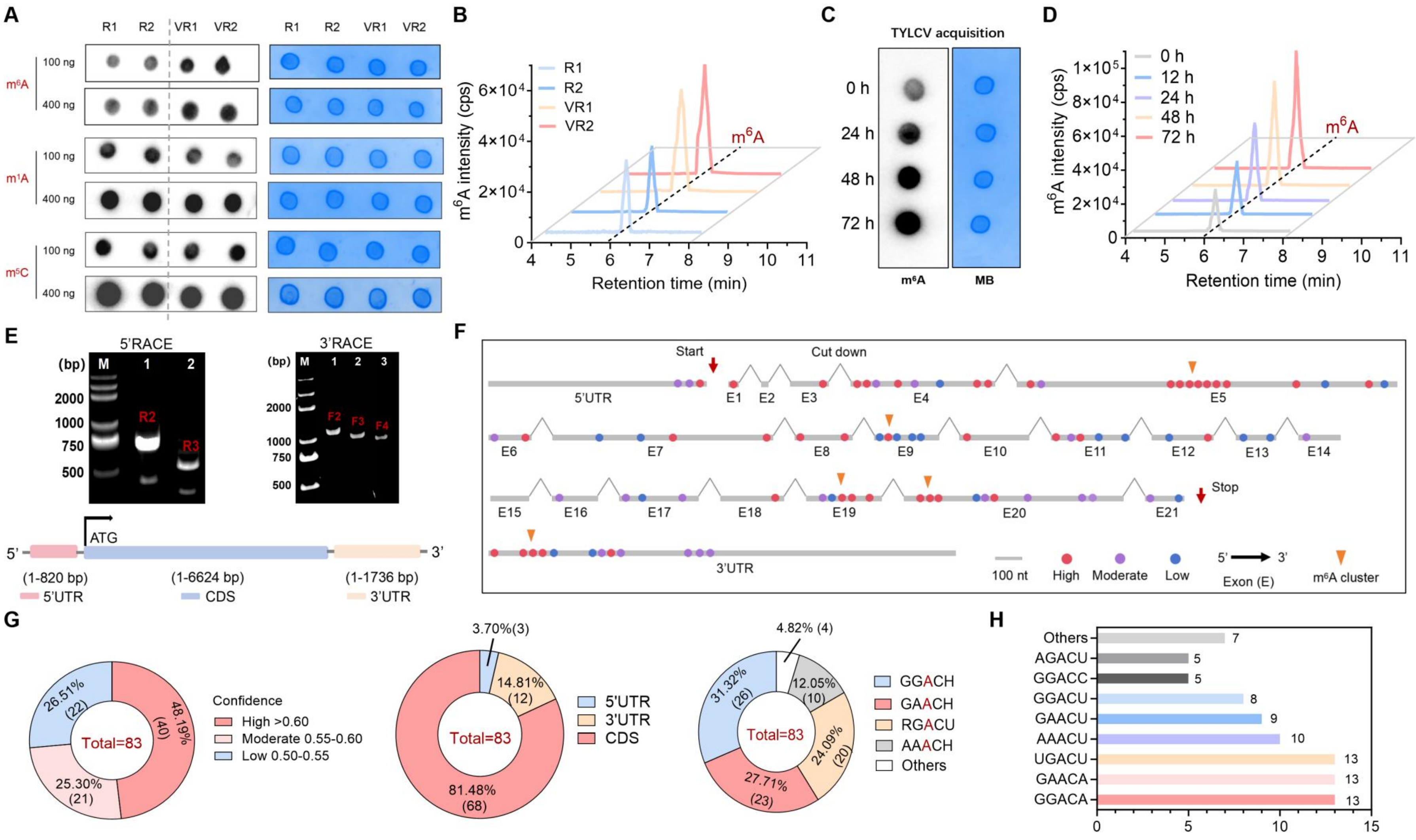

A
R1 R2 VR1 VR2
m⁶A
m¹A
m⁵C
100 ng
400 ng
B
m⁶A intensity (cps)
Retention time (min)
R1
R2
VR1
VR2
m⁶A
C
TYLCV acquisition
0 h
24 h
48 h
72 h
m⁶A
MB
D
m⁶A intensity (cps)
Retention time (min)
0 h
12 h
24 h
48 h
72 h
E
5'RACE
3'RACE
(bp)
M 1 2 3
R2 R3
F2 F3 F4
ATG
(1-820 bp)
(1-6624 bp)
(1-1736 bp)
5'UTR
CDS
3'UTR
F
Start
Cut down
Stop
5'UTR
3'UTR
100 nt
High
Moderate
Low
Exon (E)
m⁶A cluster
G
Confidence
High >0.60
Moderate 0.55-0.60
Low 0.50-0.55
Total=83
48.19% (40)
25.30% (21)
26.51% (22)
3.70%(3)
14.81% (12)
81.48% (68)
5'UTR
3'UTR
CDS
4.82% (4)
12.05% (10)
24.09% (20)
27.71% (23)
31.32% (26)
GGACH
GAACH
RGACU
AAACH
Others
H
Others 7
AGACU 5
GGACC 5
GGACU 8
GAACU 9
AAACU 10
UGACU 13
GAACA 13
GGACA 13

**Fig. S10 Mapping m$^6$A landscape of *Vg* gene in the vector *B. tabaci* MED.** (**A**) Dot blot analysis of the key RNA modifications (m$^6$A, m$^1$A and m$^5$C) in female adults of the experimental strains. Methylene blue staining (right blots) was used as a loading control to detect input RNA. (**B**) UPLC-MS/MS analysis of the global m$^6$A modification level of RNA samples extracted from female adults of the experimental strains. (**C**, **D**) m$^6$A level in female adults of non-viruliferous strains R1 following TYLCV acquisition at various time points through dot blot (**C**) and UPLC-MS/MS (**D**) analysis. (**E**) Rapid Amplification of cDNA Ends (RACE) was performed to generate full-length cDNAs of 5'UTR and 3'UTR of *Vg* gene. (**F**) A schematic of predicted m$^6$A sites on whole *Vg* gege through SRAMP server. (**G**, **H**) Distribution of *Vg* m$^6$A sites: proportion of m$^6$A confidence, m$^6$A sits in 5'UTR, CDS and 3'UTR, key m$^6$A motifs and TOP ranking list of m$^6$A motifs.

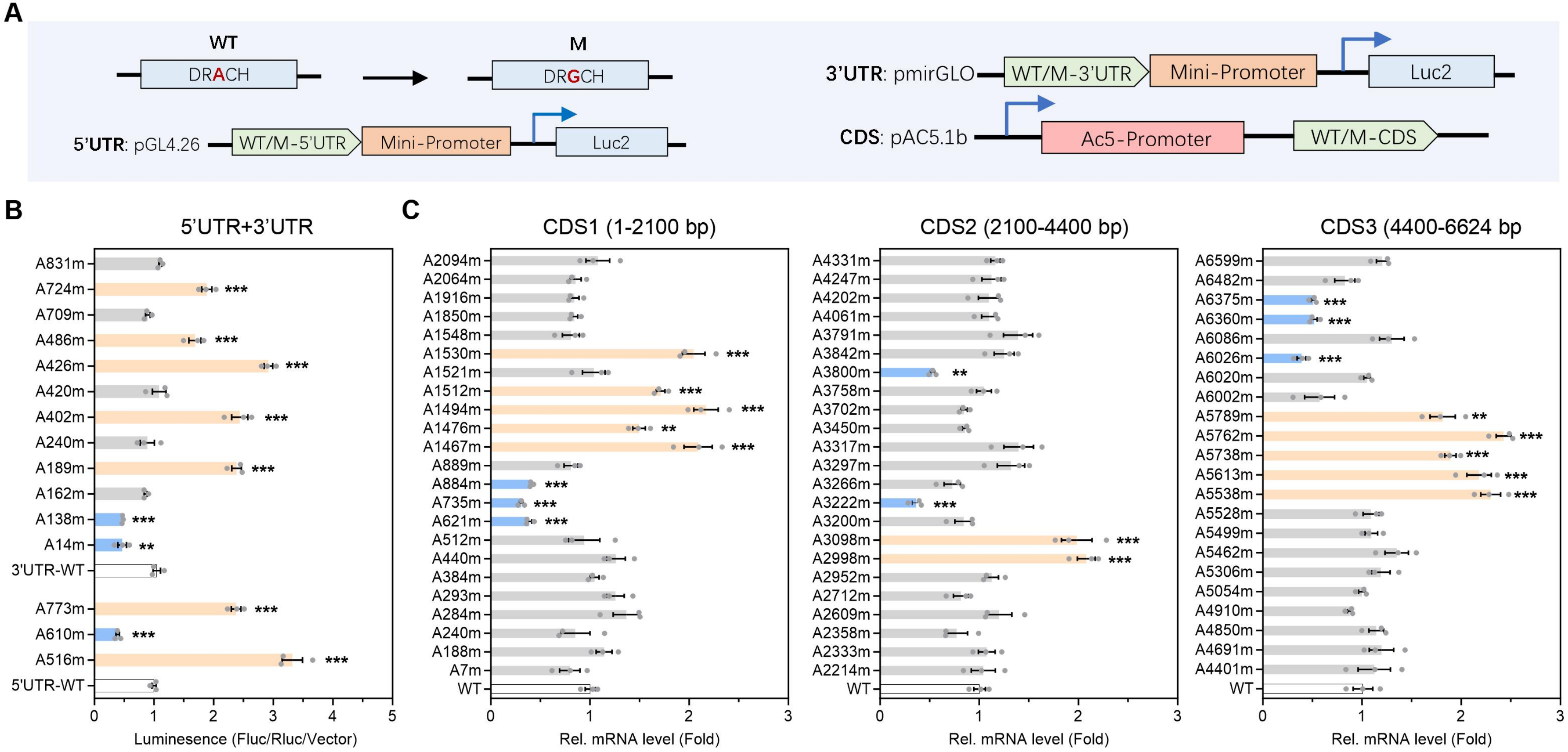

A
WT
DRACH
M
DRGCH
5'UTR: pGL4.26
WT/M-5'UTR
Mini-Promoter
Luc2
3'UTR: pmirGLO
WT/M-3'UTR
Mini-Promoter
Luc2
CDS: pAC5.1b
Ac5-Promoter
WT/M-CDS
B
5'UTR+3'UTR
Luminesence (Fluc/Rluc/Vector)
C
CDS1 (1-2100 bp)
CDS2 (2100-4400 bp)
CDS3 (4400-6624 bp
Rel. mRNA level (Fold)

**Fig. S11 Identifying the role of predicted $m^6A$ sites in regulating *Vg*.** (**A**) Schematic diagrams of various vectors containing PCR fragments of the *Vg* gene used to functionally characterize $m^6A$ sites. (**B**) Dual-luciferase reporter gene assay of the functional activity of predicted $m^6A$ sites in the 5'UTR and 3'UTR of *Vg* constructs after transfection into Drosophila S2 cells (n = 3, Student's *t* test: ****P* < 0.001). (**C**) qRT-PCR analysis of relative gene expression of *Vg* following transfection of the wide-type *Vg* CDS and the corresponding mutant constructs into Drosophila S2 cells (n = 3, Student 's *t* test: ****P* < 0.001).

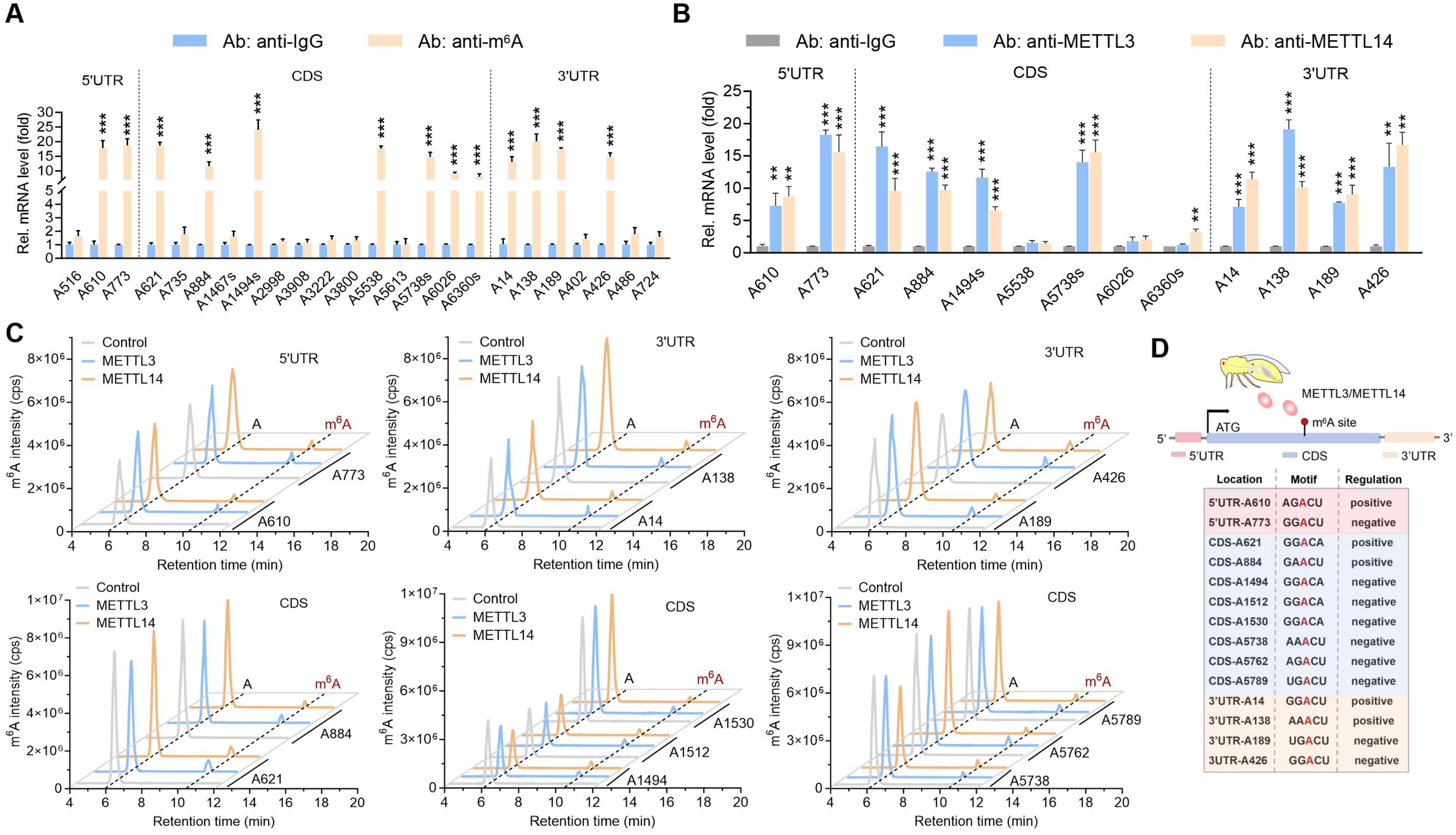
A
Ab: anti-IgG
Ab: anti-m6A
5'UTR
CDS
3'UTR
Rel. mRNA level (fold)
A516
A610
A773
A621
A735
A884
A1467s
A1494s
A2998
A3908
A3222
A3800
A5538
A5613
A5738s
A6026
A6360s
A14
A138
A189
A402
A426
A486
A724
B
Ab: anti-IgG
Ab: anti-METTL3
Ab: anti-METTL14
C
Control
METTL3
METTL14
m6A intensity (cps)
Retention time (min)
A
m6A
A610
A773
A14
A138
A189
A426
A621
A884
A1494
A1512
A1530
A5738
A5762
A5789
D
METTL3/METTL14
ATG
m6A site
5'UTR
CDS
3'UTR
Location
Motif
Regulation
5'UTR-A610 AGACU positive
5'UTR-A773 GGACU negative
CDS-A621 GGACA positive
CDS-A884 GAACU positive
CDS-A1494 GGACA negative
CDS-A1512 GGACA negative
CDS-A1530 GGACA negative
CDS-A5738 AAACU negative
CDS-A5762 AGACU negative
CDS-A5789 UGACU negative
3'UTR-A14 GGACU positive
3'UTR-A138 AAACU positive
3'UTR-A189 UGACU negative
3UTR-A426 GGACU negative

**Fig. S12 The key methyltransferases METTL3 and METTL14 bind to the putative sites of *Vg* for $m^6A$ modification.** (**A**) MeRIP analysis of binding enrichment of $m^6A$ to the putative $m^6A$ sites of *Vg* in VR1 strain (n = 3, Student's *t* test: $^{**}P < 0.01$, $^{***}P < 0.001$). (**B**) MeRIP analysis of binding enrichment of METTL3 and METTL14 protein to the putative $m^6A$ sites of *Vg* mRNA in VR1 strain (n = 3, Student's *t* test: $^{**}P < 0.01$, $^{***}P < 0.001$). (**C**) UPLC-MS/MS analysis of $m^6A$ modification using recombinant METTL3 and METTL14. A 15bp-length single strain RNA (ssRNA) containing the *Vg* $m^6A$ sites was used as a substrate of METTL3 and METTL14. (**D**) Schematic diagram of the functional $m^6A$ sites and their role in epigenetic regulation of *Vg*.

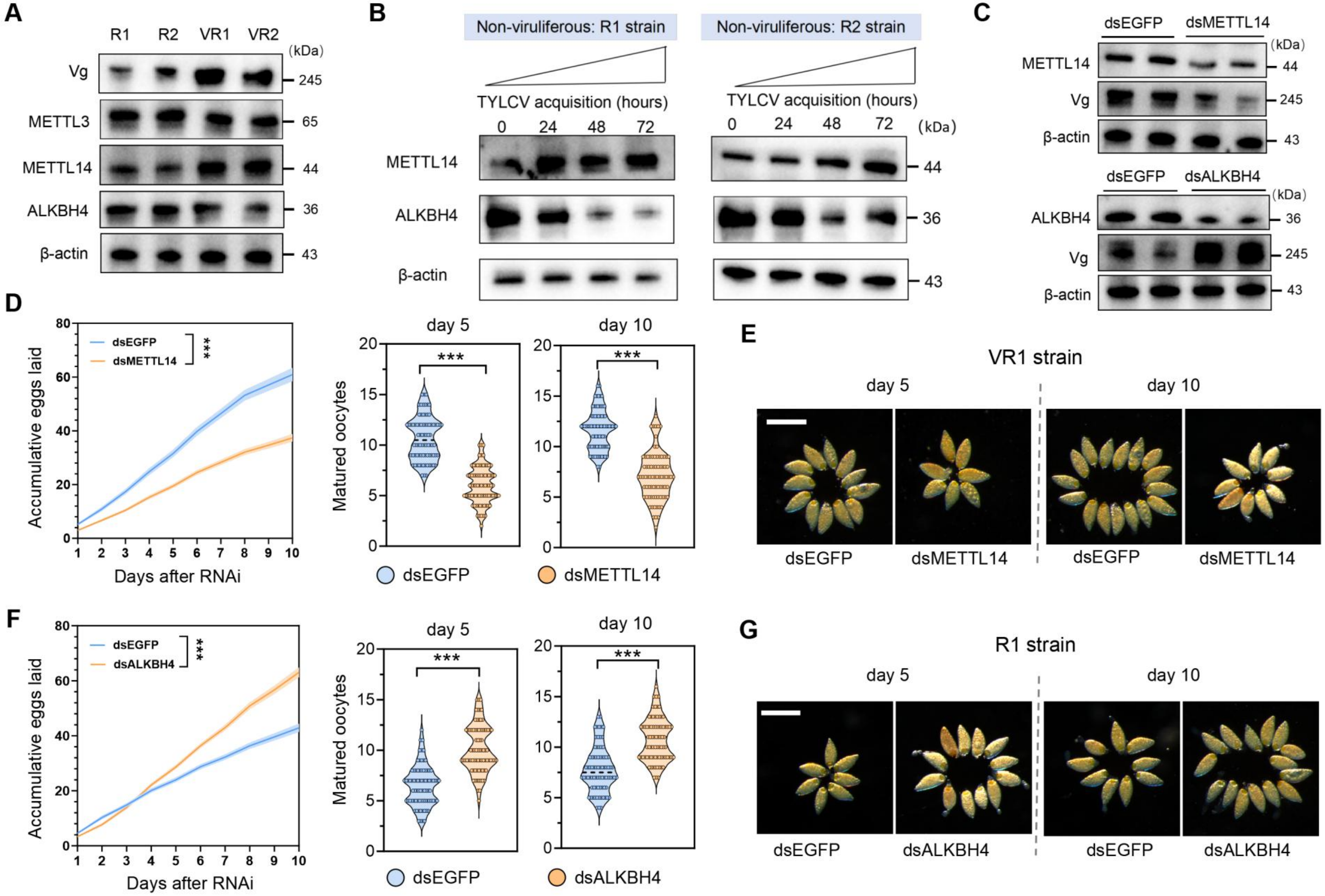

A
R1 R2 VR1 VR2
(kDa)
Vg 245
METTL3 65
METTL14 44
ALKBH4 36
β-actin 43
B
Non-viruliferous: R1 strain
Non-viruliferous: R2 strain
TYLCV acquisition (hours)
0 24 48 72
(kDa)
METTL14 44
ALKBH4 36
β-actin 43
C
dsEGFP dsMETTL14
METTL14 44
Vg 245
β-actin 43
dsEGFP dsALKBH4
ALKBH4 36
Vg 245
β-actin 43
D
Accumulative eggs laid
Days after RNAi
dsEGFP
dsMETTL14
***
day 5
day 10
Matured oocytes
dsEGFP
dsMETTL14
E
VR1 strain
day 5
day 10
dsEGFP
dsMETTL14
F
Accumulative eggs laid
Days after RNAi
dsEGFP
dsALKBH4
***
Matured oocytes
dsEGFP
dsALKBH4
G
R1 strain
day 5
day 10
dsEGFP
dsALKBH4

**Fig. S13 The writer METTL14 and eraser ALKBH4 coordinately enhance Vg expression leading to reproductive compensation in the viruliferous whitefly**. (**A**) Western blot assay of protein level of Vg, METTL3, METTL14 and ALKBH4 in female adults of the experimental strains. (**B**) Protein level of METTL14 and ALKBH4 following TYLCV acquisition at various time points in the vector. (**C**) Qualification of Vg mRNA level after RNAi knockdown of METTL14 or ALKBH4 in female adults of the viruliferous strain VR1 (n = 3, Student's *t* test: ***$P < 0.001$). (**D**-**G**) Phenotypic observation of female fecundity and mature oocytes following silencing of METTL14 (**D** and **E**) and ALKBH4 (**F** and **G**) in the viruliferous strain VR1 (n ≥ 60, Student’s *t* test: ***$P < 0.001$). Scar bar, 200 μm.

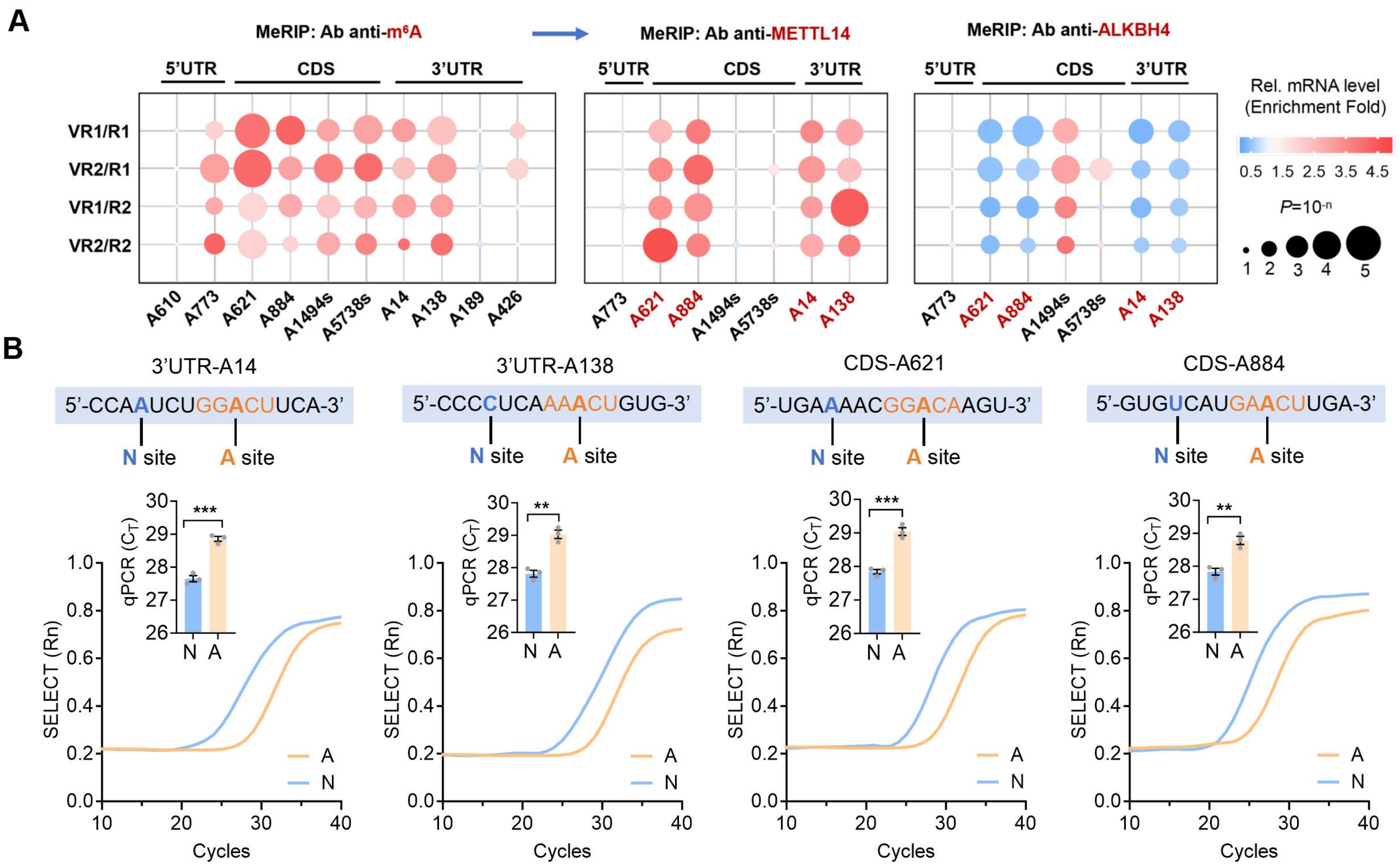
A
MeRIP: Ab anti-m6A
MeRIP: Ab anti-METTL14
MeRIP: Ab anti-ALKBH4
5'UTR
CDS
3'UTR
VR1/R1
VR2/R1
VR1/R2
VR2/R2
A610
A773
A621
A884
A1494s
A5738s
A14
A138
A189
A426
Rel. mRNA level
(Enrichment Fold)
0.5 1.5 2.5 3.5 4.5
P=10-n
1 2 3 4 5
B
3'UTR-A14
5'-CCAAUCUGGACUUCA-3'
3'UTR-A138
5'-CCCCUCAAAACUGUG-3'
CDS-A621
5'-UGAAAACGGACAAGU-3'
CDS-A884
5'-GUGUCAUGAACUUGA-3'
N site
A site
qPCR (CT)
***
**
N
A
SELECT (Rn)
Cycles

**Fig. S14 Identification of functional m$^6$A sites of *Vg* involved in reproductive compensation in the viruliferous whitefly**. (**A**) MeRIP-qPCR assay of binding enrichment of the m$^6$A , METTL3 and ALKBH4 protein to the putative m$^6$A sites of *Vg* mRNA in female adults of the experimental strains (n = 3, Student's *t* test). (**B**) The SELECT results for validation of m$^6$A modification on the functional m$^6$A sites of *Vg* at single-base resolution in female adults of the whitefly strain VR1 (n = 3, Student's *t* test: $^{**}P < 0.01$, $^{***}P < 0.001$ ).

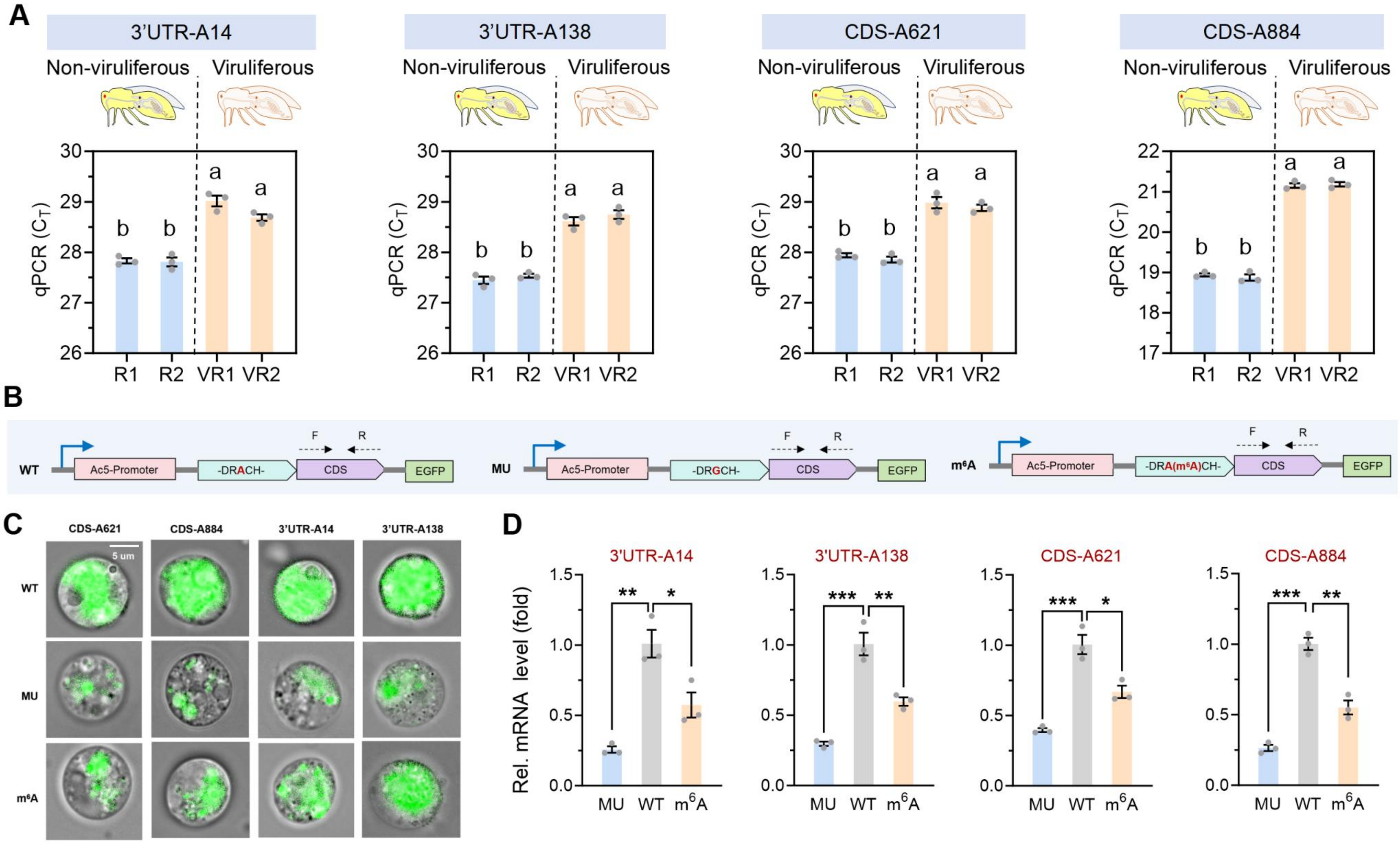
A
3'UTR-A14
3'UTR-A138
CDS-A621
CDS-A884
Non-viruliferous
Viruliferous
qPCR (C_T)
R1 R2 VR1 VR2
B
WT
MU
m6A
Ac5-Promoter
-DRACH-
-DRGCH-
-DRA(m6A)CH-
CDS
EGFP
F
R
C
CDS-A621
CDS-A884
3'UTR-A14
3'UTR-A138
5 um
WT
MU
m6A
D
3'UTR-A14
3'UTR-A138
CDS-A621
CDS-A884
Rel. mRNA level (fold)
MU WT m6A

**Fig. S15 Quantification of the four functional m$^6$A sites regulating *Vg* expression. (A)** The SELECT results for comparison of m$^6$A modification on the functional m$^6$A sites of *Vg* at single-base resolution in the vector (Tukey's HSD multiple comparisons test: $P < 0.05$). (**B**) Schematic diagram of there types of constructs containing the m$^6$A motifs cloned into the pAC5.1b-G3-X2-EGFP vector to quantify *Vg* expression following transfection into Drosophila S2 cells. (**C**) Representative images displaying fluorescence intensity of Vg in S2 cells following transfection of the constructs into S2 cells for 72 h. Scar bar, 5 μm. (**D**) qPCR analysis of Vg mRNA level in S2 cells after transfection of the constructs into S2 cells for 72 h (n = 3, Student's *t* test: *$P < 0.05$, **$P < 0.01$, ***$P < 0.001$).

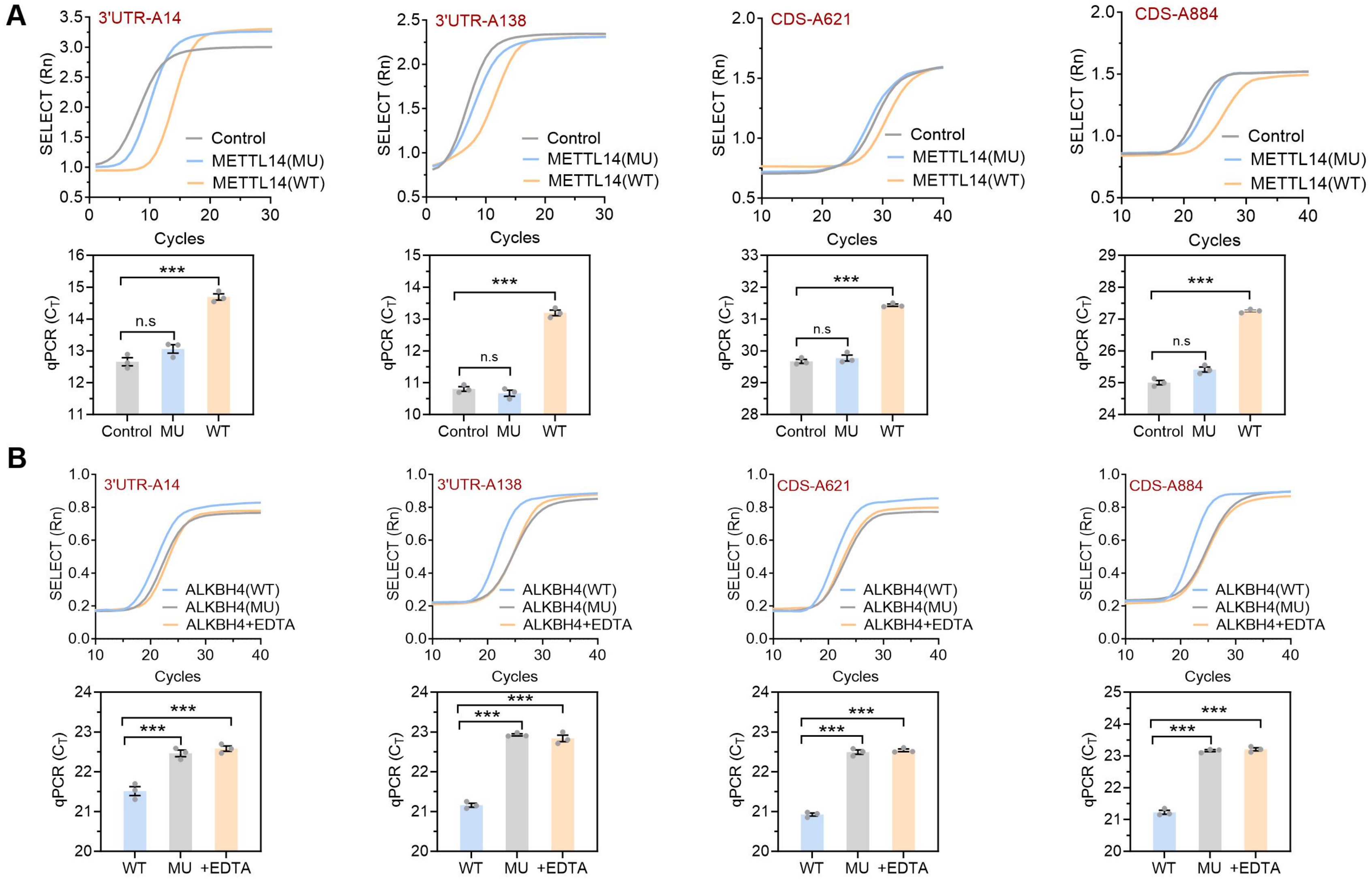
A
3'UTR-A14
3'UTR-A138
CDS-A621
CDS-A884
SELECT (Rn)
Cycles
Control
METTL14(MU)
METTL14(WT)
qPCR ($C_T$)
n.s
***
Control
MU
WT
B
ALKBH4(WT)
ALKBH4(MU)
ALKBH4+EDTA
WT
MU
+EDTA

**Fig. S16 Functional role of METTL14 and ALKBH4 in controlling m$^6$A modification of *Vg* at single-base resolution.** (**A**) The SELECT assays for validation of the functional m$^6$A sites in female adults of the whitefly strain VR1 using the recombinant METTL14 protein for methylation (n = 3, Student 's *t* test: $^{***}P < 0.001$). (**B**) The SELECT assays for validation of the functional m$^6$A sites in female adults of the whitefly strain VR1 using the recombinant ALKBH4 protein for demethylation (n = 3, Student's *t* test: $^{***}P < 0.001$).

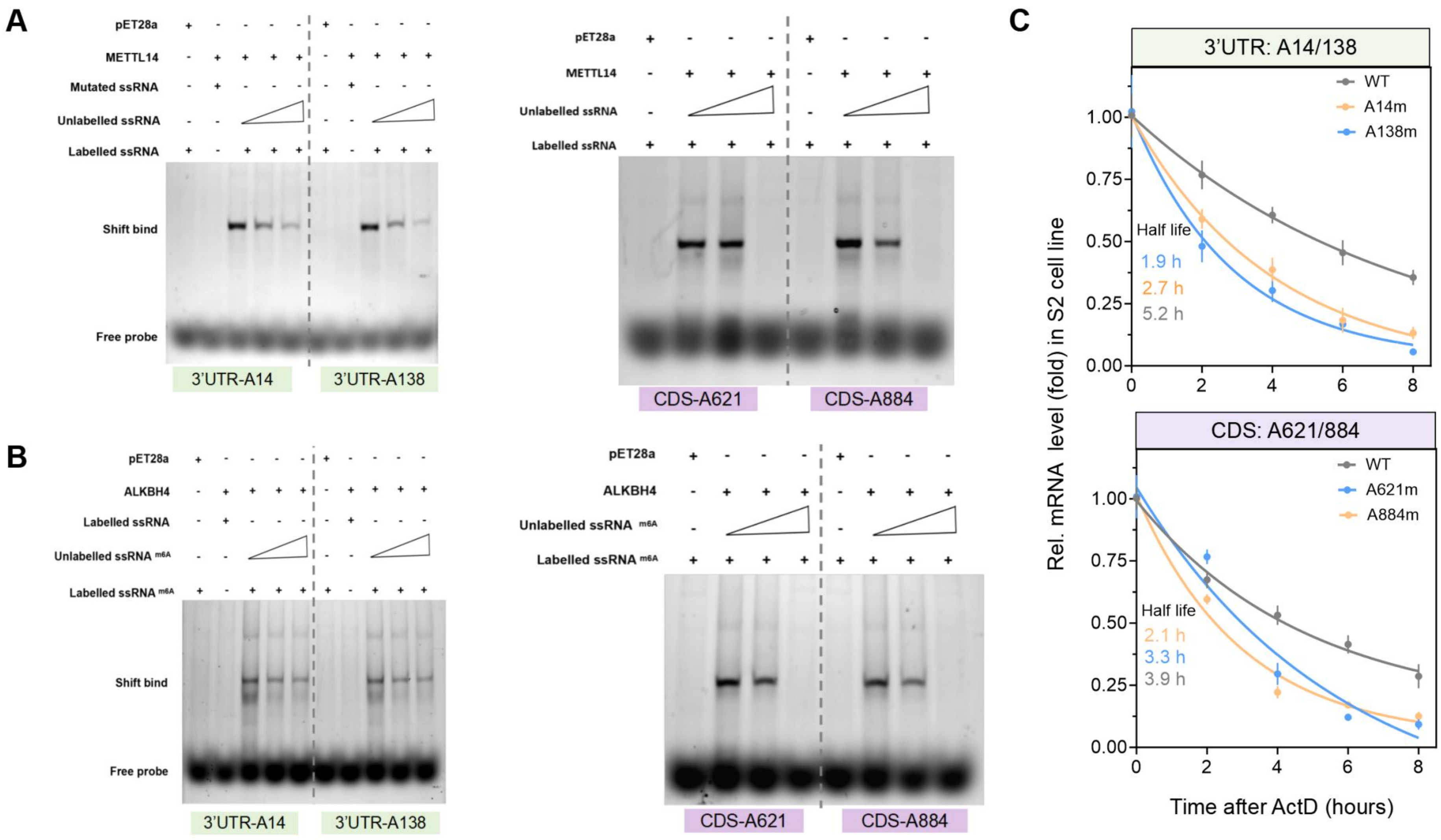
A
pET28a
METTL14
Mutated ssRNA
Unlabelled ssRNA
Labelled ssRNA
Shift bind
Free probe
3'UTR-A14
3'UTR-A138
CDS-A621
CDS-A884
B
ALKBH4
Labelled ssRNA
Unlabelled ssRNA m6A
Labelled ssRNA m6A
C
3'UTR: A14/138
WT
A14m
A138m
Half life
1.9 h
2.7 h
5.2 h
CDS: A621/884
A621m
A884m
2.1 h
3.3 h
3.9 h
Rel. mRNA level (fold) in S2 cell line
Time after ActD (hours)

**Fig. S17 Binding of METTL14 and ALKBH4 to *Vg* m$^6$A sites, resulting in stabilizing *Vg* transcripts.** (**A**) EMSA assay of the binding of recombinant METTL14 to the functional m$^6$A sites of *Vg*. A 25bp-length single strain RNA (ssRNA) containing the m$^6$A sites of *Vg* was used as a probe. (**B**) EMSA assay of the binding of recombinant ALKBH4 to the functional m$^6$A sites of *Vg*. The ssRNA with m$^6$A modification on A site containing the m$^6$A sites of *Vg* was used as a probe. (**C**) mRNA stability of *Vg* following transfection of wide-type (WT) and mutant (m) constructs in to Drosophila S2 cells.

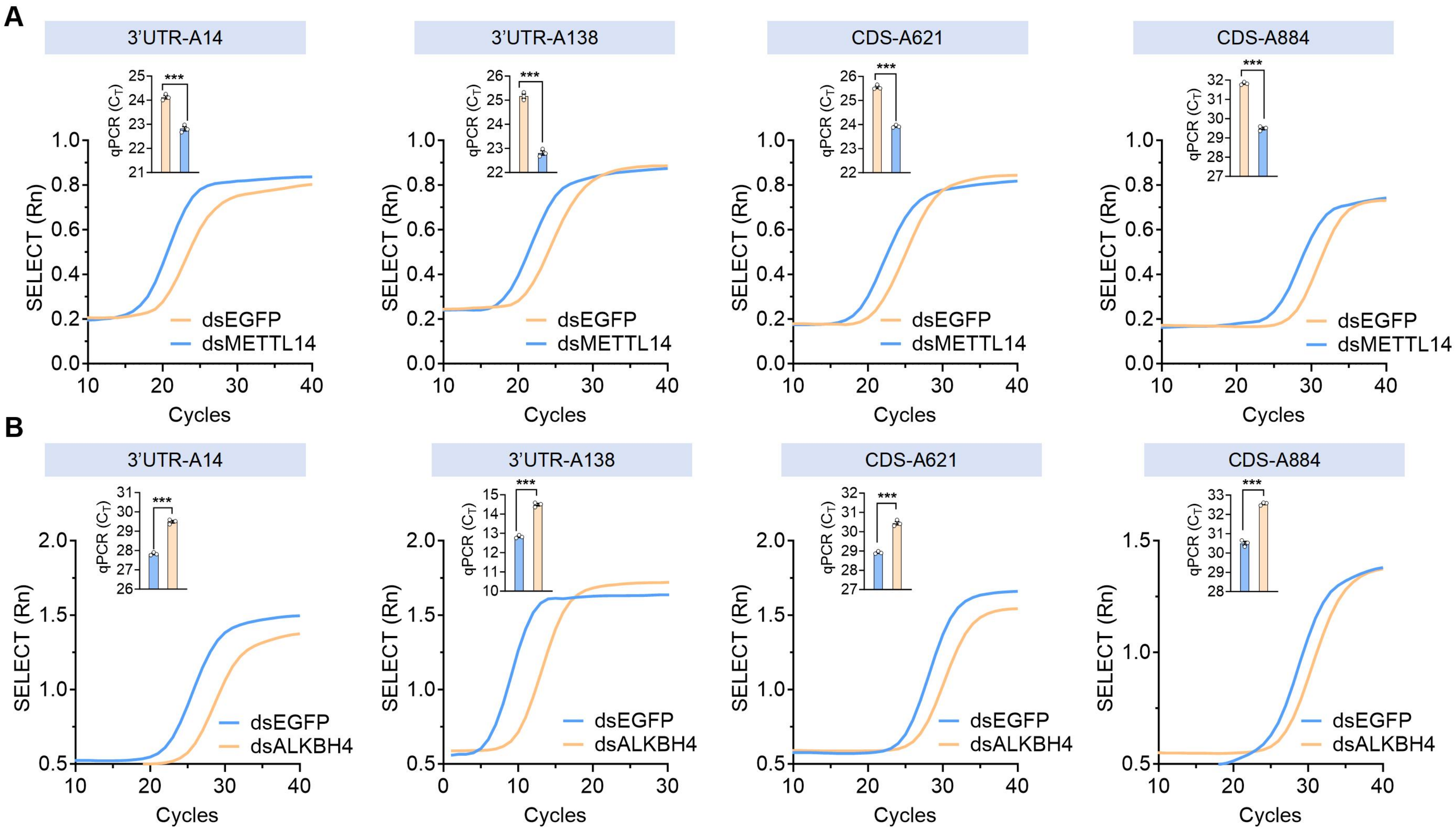

A
3'UTR-A14
3'UTR-A138
CDS-A621
CDS-A884
qPCR ($C_T$)
***
SELECT (Rn)
Cycles
dsEGFP
dsMETTL14
B
3'UTR-A14
3'UTR-A138
CDS-A621
CDS-A884
qPCR ($C_T$)
***
SELECT (Rn)
Cycles
dsEGFP
dsALKBH4

**Fig. S18 RNAi uncovers the importance of METTL14 and ALKBH4 in regulating m$^6$A modification on *Vg* at single-base resolution.** (**A**, **B**) The SELECT results for detection of m$^6$A modification on the functional m$^6$A sites of *Vg* post RNAi knockdown of METTL14 (**A**) or ALKBH4 (**B**) (n = 3, Student's *t* test: ****P* < 0.001).

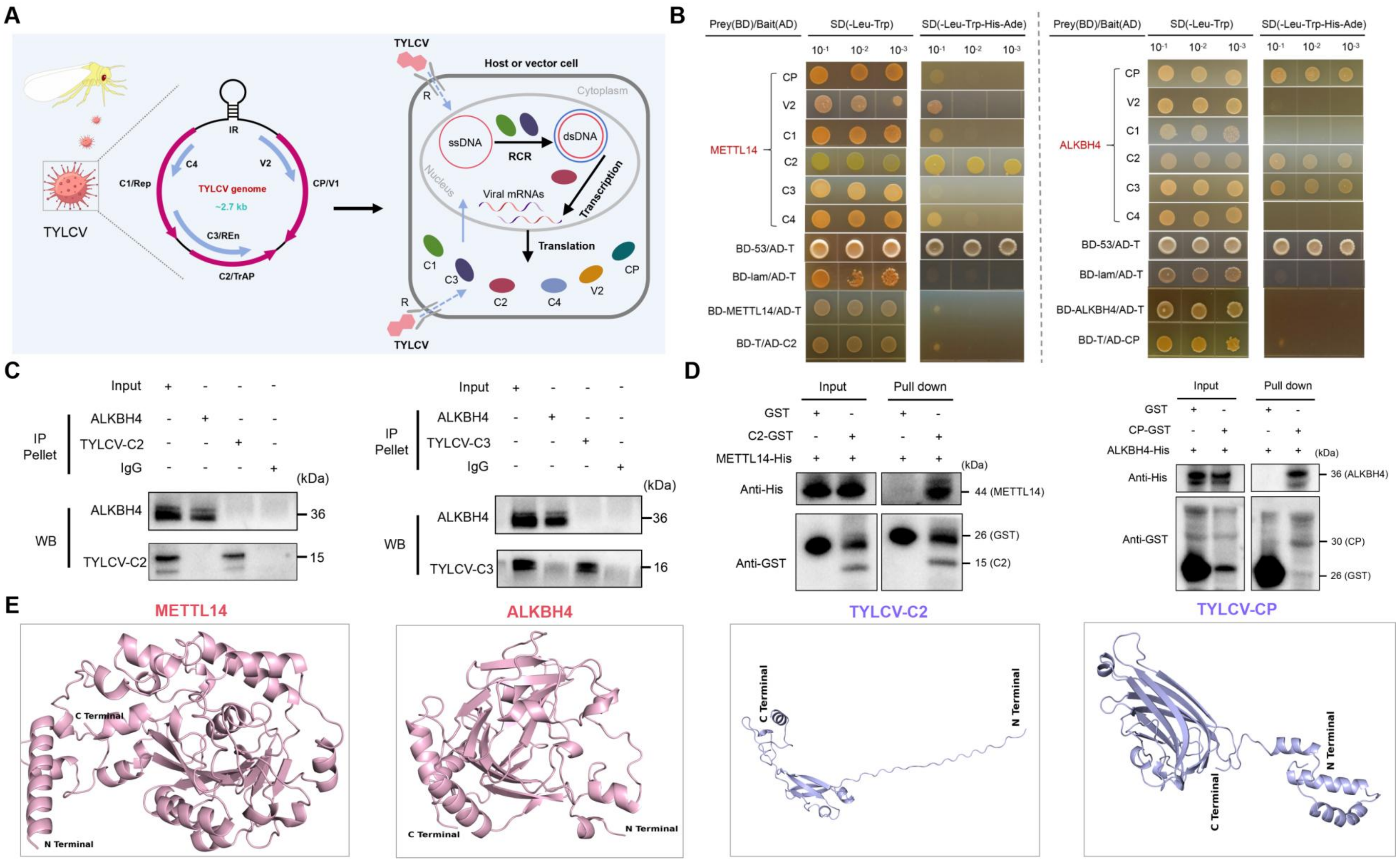
A
TYLCV
TYLCV genome
~2.7 kb
IR
C4
V2
C1/Rep
CP/V1
C3/REn
C2/TrAP
Host or vector cell
Cytoplasm
Nucleus
ssDNA
dsDNA
RCR
Transcription
Viral mRNAs
Translation
R
C1
C2
C3
C4
V2
CP
B
Prey(BD)/Bait(AD)
SD(-Leu-Trp)
SD(-Leu-Trp-His-Ade)
10-1 10-2 10-3
METTL14
ALKBH4
CP
V2
C1
C2
C3
C4
BD-53/AD-T
BD-lam/AD-T
BD-METTL14/AD-T
BD-T/AD-C2
BD-ALKBH4/AD-T
BD-T/AD-CP
C
Input
IP Pellet
ALKBH4
TYLCV-C2
TYLCV-C3
IgG
(kDa)
WB
36
15
16
D
Input
Pull down
GST
C2-GST
CP-GST
METTL14-His
ALKBH4-His
Anti-His
Anti-GST
44 (METTL14)
26 (GST)
15 (C2)
36 (ALKBH4)
30 (CP)
E
METTL14
ALKBH4
TYLCV-C2
TYLCV-CP
C Terminal
N Terminal

**Fig. S19 The direct binding of TYLCV protein C2 and CP to vector's METTL14 and ALKBH4, respectively.** (**A**) Schematic diagram showing six structural genes of TYLCV genome and their replication and translation in host or vector cell. (**B**) Y2H assay of binding interactions of METTL14 and ALKBH4 to TYLCV structural proteins. (**C**) IP assays demonstrating no interaction between ALKBH4 and C2 or C3 in the vector strain VR1. (**D**) GST pull-down assays of the direct interaction between METTL14 and viral C2 protein, and between ALKBH4 and viral CP protein. (**E**) Protein structure of METTL14, ALKBH4 and viral C2 and CP predicted by AlphaFold3.

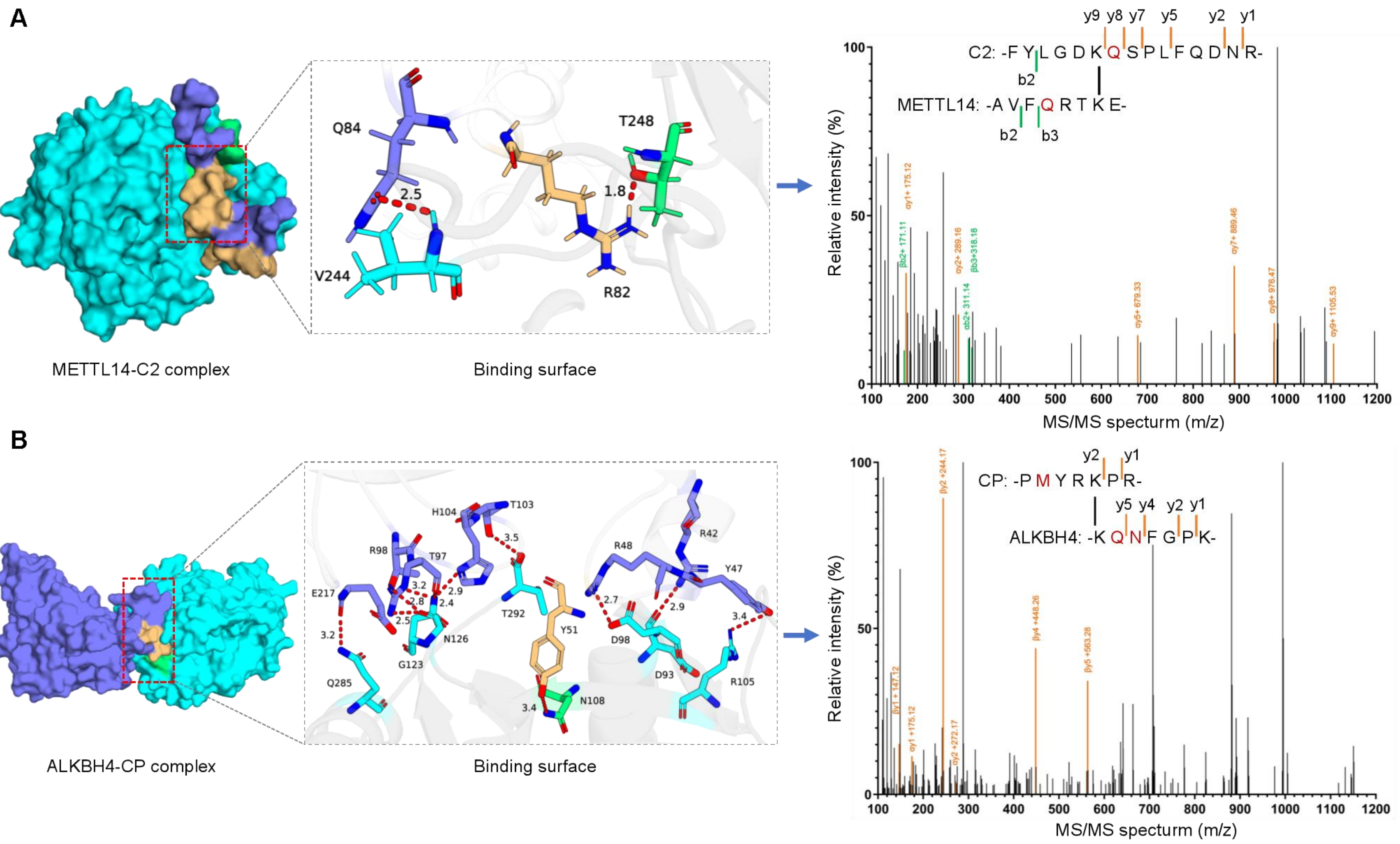
A
Q84
T248
2.5
1.8
V244
R82
METTL14-C2 complex
Binding surface
y9 y8 y7 y5 y2 y1
C2: -F Y L G D K Q S P L F Q D N R-
b2
METTL14: -A V F Q R T K E-
b2 b3
Relative intensity (%)
MS/MS specturm (m/z)
B
H104
T103
R98
T97
E217
3.2
2.8
2.9
2.4
2.5
3.5
T292
Y51
N126
G123
Q285
N108
3.4
R48
R42
Y47
2.7
2.9
3.4
D98
D93
R105
ALKBH4-CP complex
Binding surface
y2 y1
CP: -P M Y R K P R-
y5 y4 y2 y1
ALKBH4: -K Q N F G P K-
Relative intensity (%)
MS/MS specturm (m/z)

**Fig. S20 The cross-linking mass spectrometry (XL-MS) analysis of binding interfaces of METTL14-C2 or ALKBH4-CP complex.** (**A**) Binding interfaces of METTL14-C2 interaction and the inter-linked peptide pairs involved in binding interaction. (**B**) Binding interfaces of ALKBH4-CP interaction and the inter-linked peptide pairs involved in binding interaction.

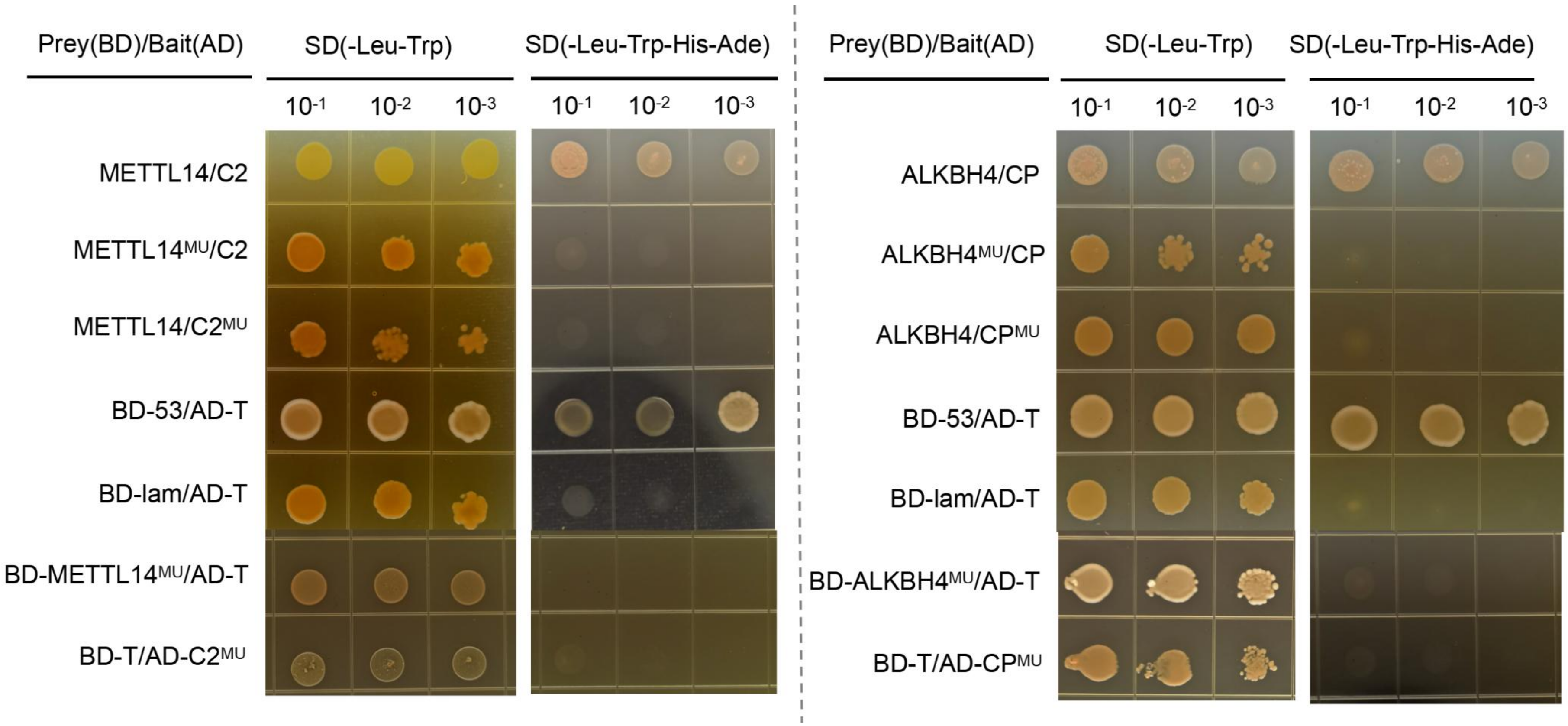


**Fig. S21 Y2H assays of the binding of METTL14- C2 and ALKBH4 -CP interaction by mutating their binding residues based on molecular docking and XL-MS data.**

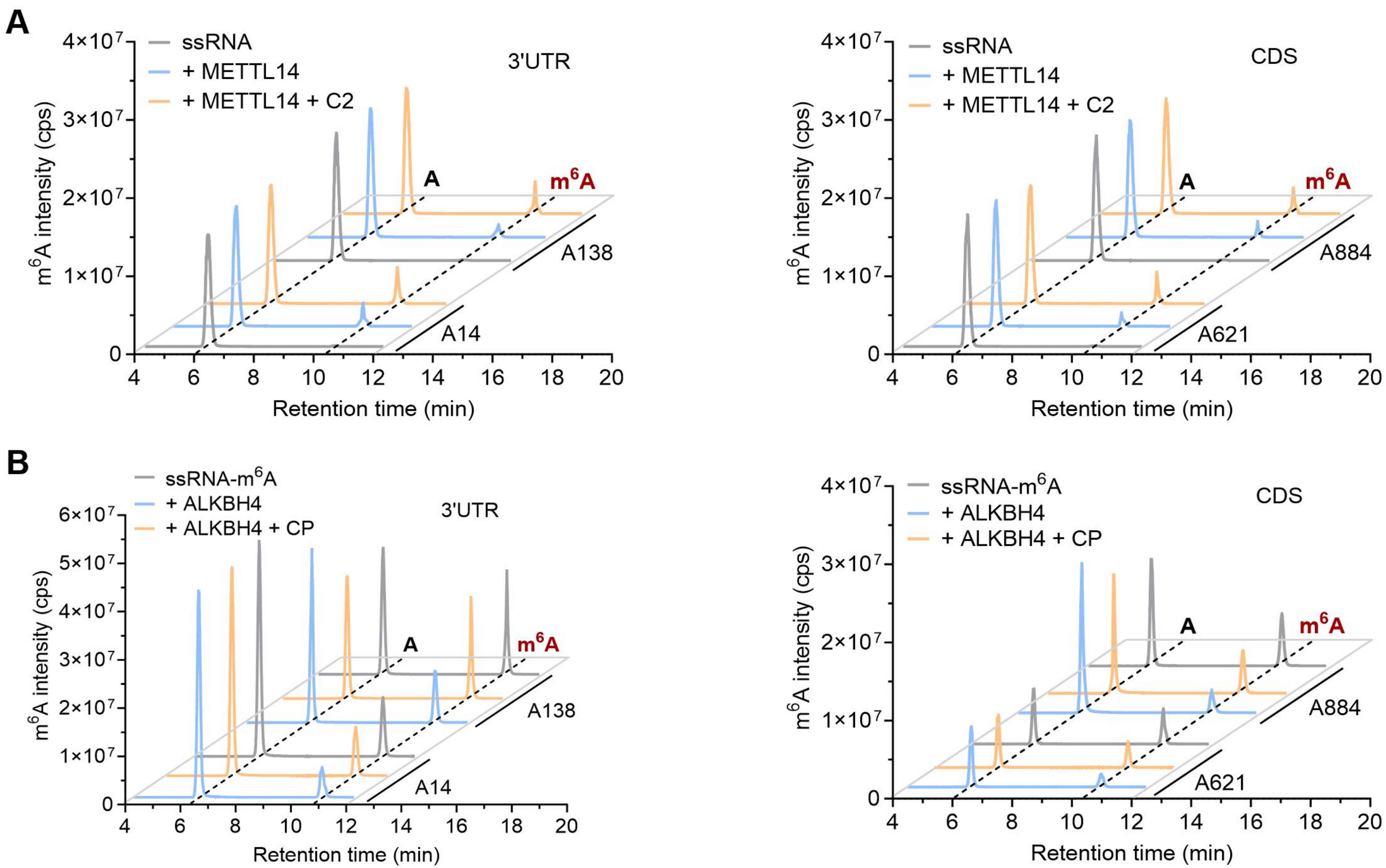
A
3'UTR
CDS
ssRNA
+ METTL14
+ METTL14 + C2
m6A intensity (cps)
Retention time (min)
A
m6A
A138
A14
A884
A621
B
ssRNA-m6A
+ ALKBH4
+ ALKBH4 + CP

**Fig. S22** UPLC-MS/MS analysis of the effect of viral protein on m$^6$A modification of *Vg* . (**A**) The presence of recombinant C2 increased m$^6$A level of each *Vg* m$^6$A sites by coincubation with METTL14. (**B**) The presence of recombinant CP increased m$^6$A level of each *Vg* m$^6$A sites by coincubation with ALKBH4.

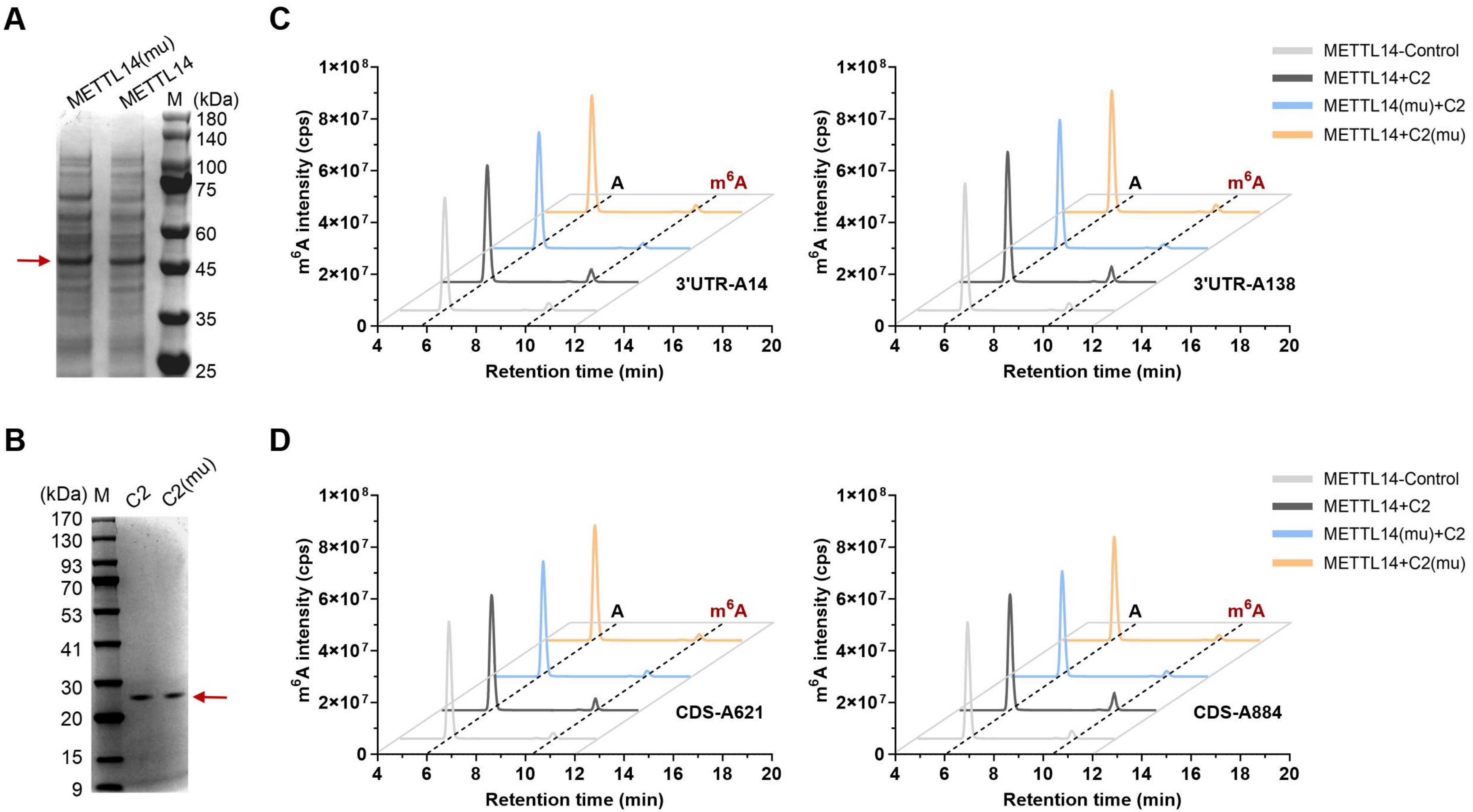

A
METTL14(mu)
METTL14
M
(kDa)
180
140
100
75
60
45
35
25
B
(kDa)
M
C2
C2(mu)
170
130
93
70
53
41
30
20
15
9
C
D
METTL14-Control
METTL14+C2
METTL14(mu)+C2
METTL14+C2(mu)
m6A intensity (cps)
Retention time (min)
A
m6A
3'UTR-A14
3'UTR-A138
CDS-A621
CDS-A884

**Fig. S23 Disruption of METTL14-C2 interaction reduces m$^6$A level of *Vg* in vitro.** (**A**, **B**) Heterologous expression of recombinant METTL14 (**A**) and viral C2 (**B**) protein in *E. coli* as detected by SDS-PAGE gel. Protein marker (M). (**C**, **D**) UPLC-MS/MS analysis of m$^6$A modification of *Vg* m$^6$A sites (**C**: 3UTR-A14, A138; **D**: CDS-A621, A884) following breaking METTL14-C2 interaction. Note that the mutated proteins (mu) of METTL14 or C2 were created by mutating their binding residues based on the results of molecular docking. A 15bp-length ssRNA covering the functional m$^6$A site of *Vg* was used as a substrate of METTL14.

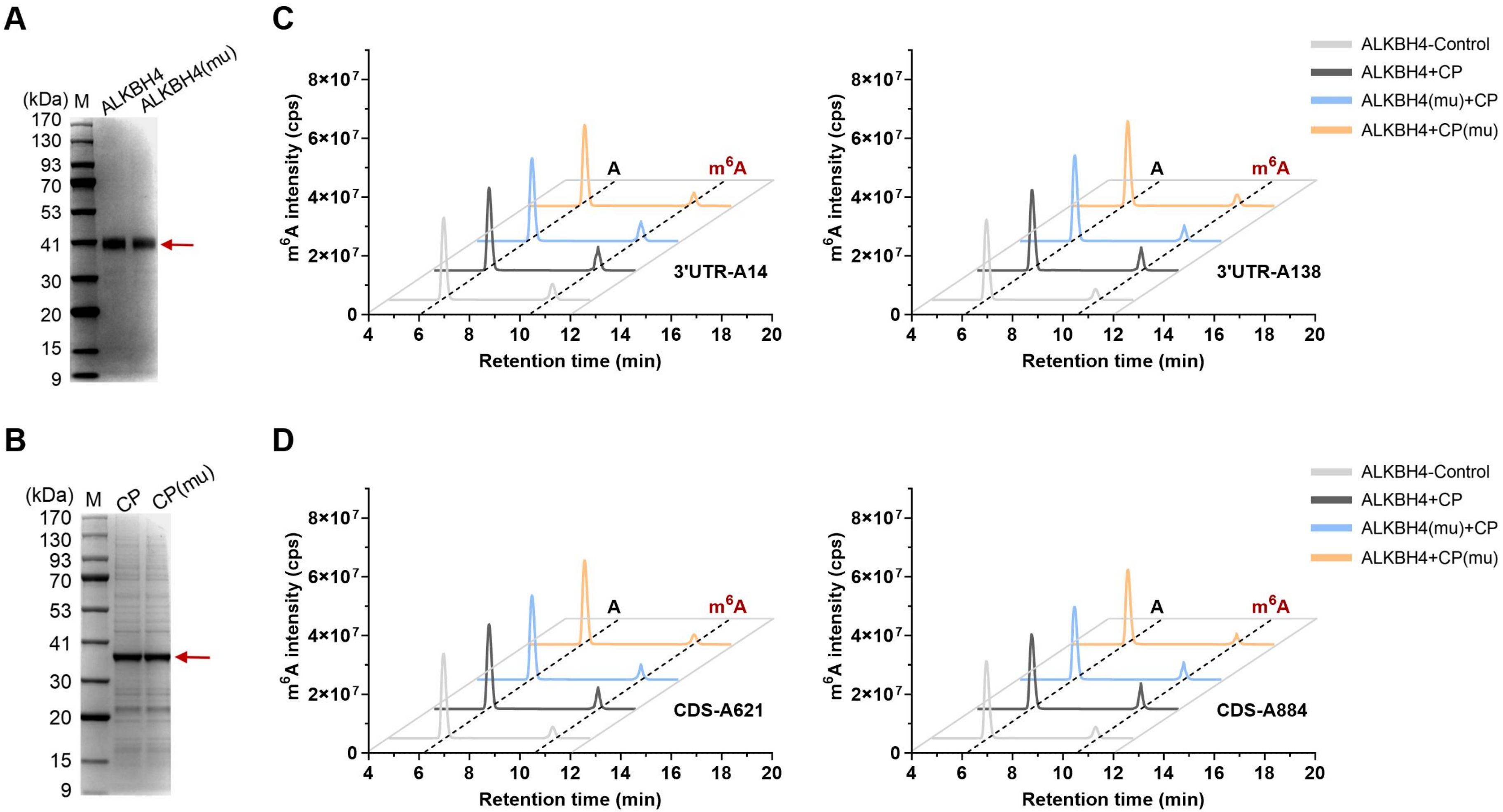

A
(kDa)
M
ALKBH4
ALKBH4(mu)
170
130
93
70
53
41
30
20
15
9
B
CP
CP(mu)
C
D
ALKBH4-Control
ALKBH4+CP
ALKBH4(mu)+CP
ALKBH4+CP(mu)
m6A intensity (cps)
Retention time (min)
A
m6A
3'UTR-A14
3'UTR-A138
CDS-A621
CDS-A884

**Fig. S24 Disruption of ALKBH4-CP interaction reduces m$^6$A level of *Vg* in vitro.** (**A**, **B**) Heterologous expression of recombinant ALKBH4 (**A**) and viral CP (**B**) protein in *E. coli* as detected by SDS-PAGE gel. Red arrow, the target protein. Protein marker (M). (**C**, **D**) UPLC-MS/MS analysis of m$^6$A modification of *Vg* m$^6$A sites (**C**: 3UTR-A14, A138; **D**: CDS-A621, A884) following breaking ALKBH4-CP interaction. Note that the mutated proteins (mu) of ALKBH4 or CP were created by mutating their binding residues based on the results of molecular docking. A 15bp-length ssRNA covering the functional m$^6$A site of *Vg* was used as a substrate of ALKBH4.

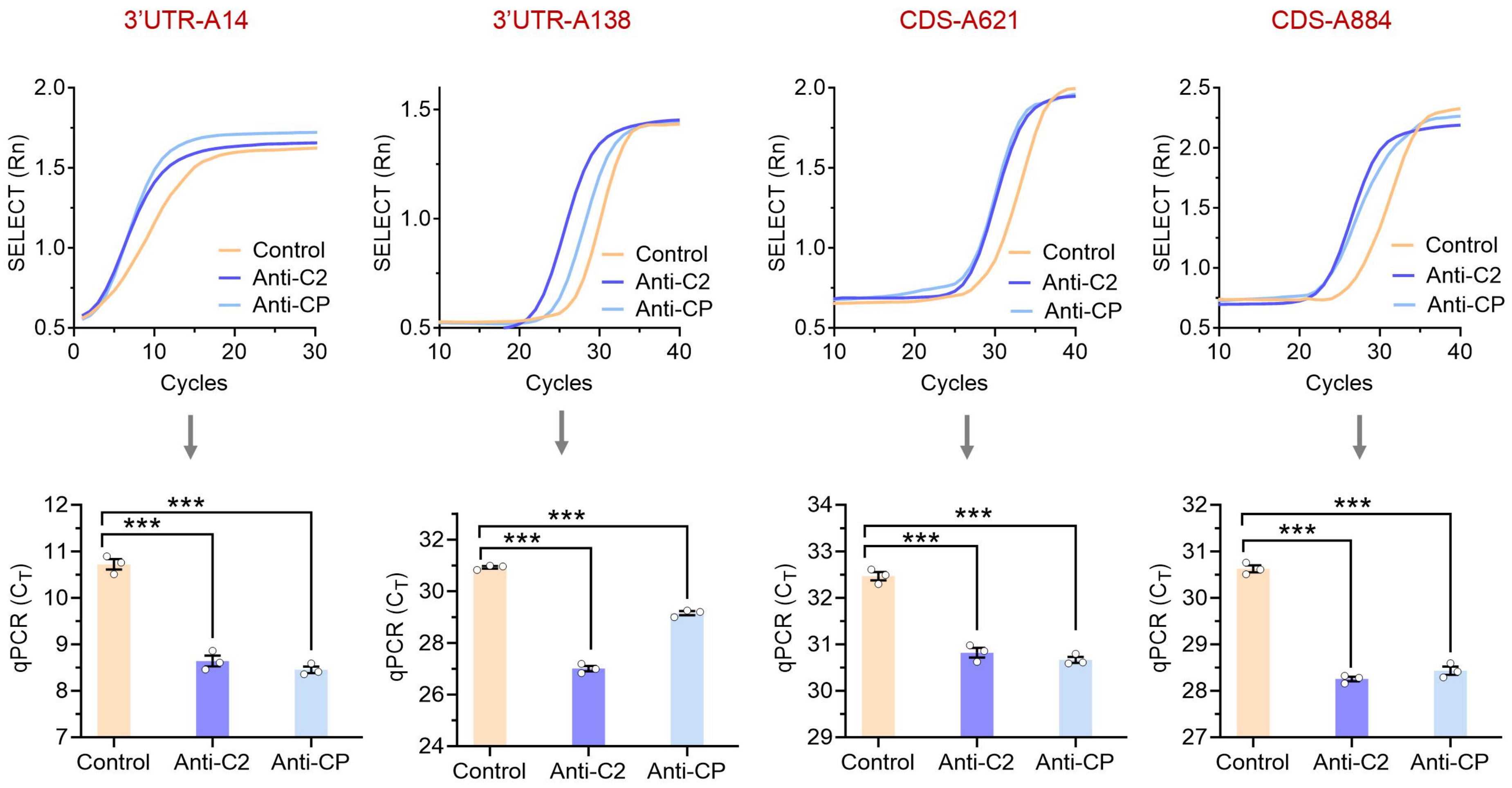

3'UTR-A14
3'UTR-A138
CDS-A621
CDS-A884
SELECT (Rn)
Cycles
Control
Anti-C2
Anti-CP
qPCR ($C_T$)
***

**Fig. S25 Inhibition of viral C2 and CP declines m$^6$A level of *Vg* in the whitefly.** The SELECT results displaying a reduced m$^6$A modification of each Vg-m$^6$A site in female adult of strain VR1 following feeding with anti-C2 or anti-CP antibody (n = 3, Student's *t* test: ***$P < 0.001$).

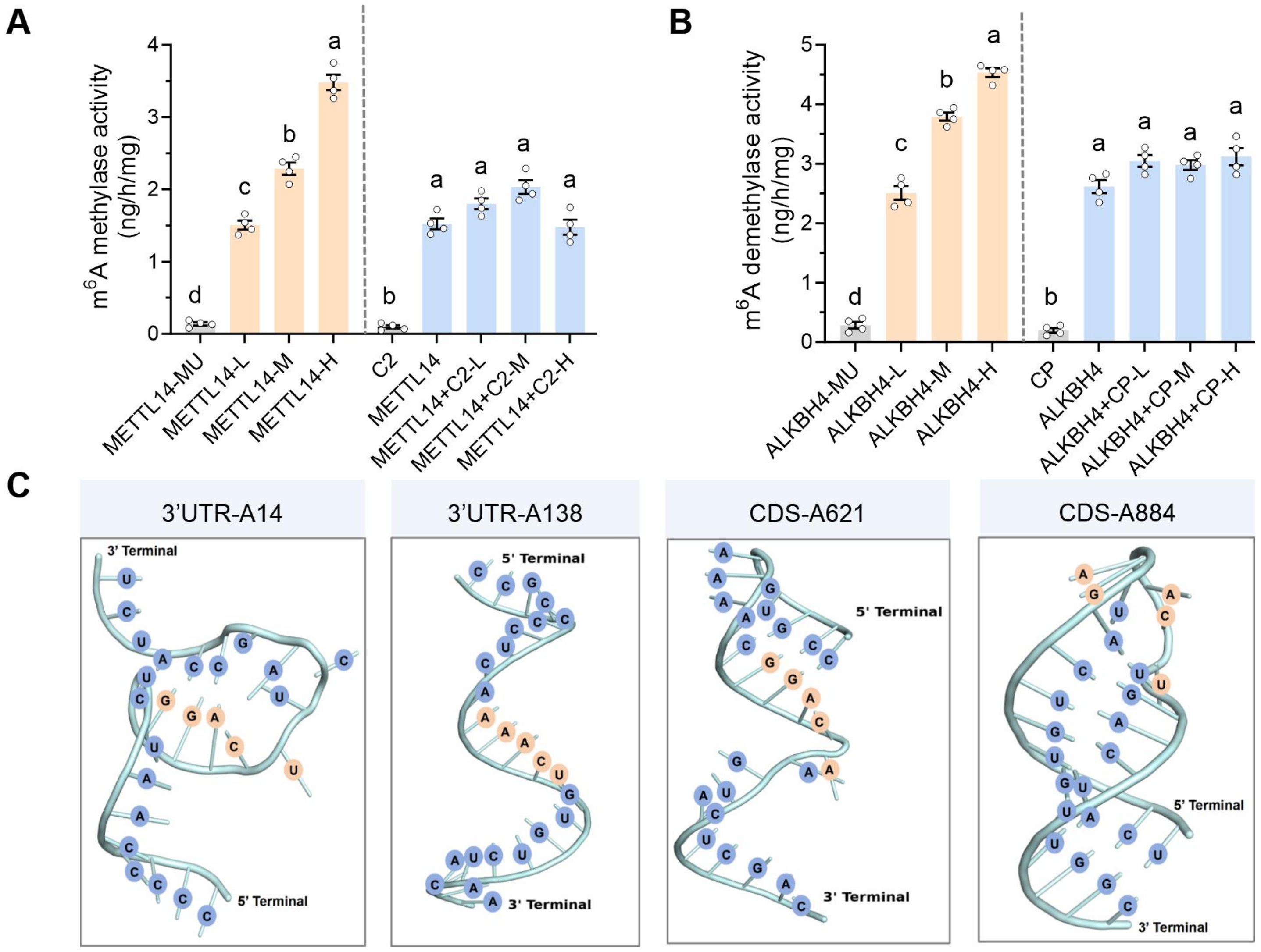
A
B
C
m6A methylase activity (ng/h/mg)
0
1
2
3
4
d
c
b
a
b
a
a
a
a
METTL14-MU
METTL14-L
METTL14-M
METTL14-H
C2
METTL14
METTL14+C2-L
METTL14+C2-M
METTL14+C2-H
m6A demethylase activity (ng/h/mg)
0
1
2
3
4
5
d
c
b
a
b
a
a
a
a
ALKBH4-MU
ALKBH4-L
ALKBH4-M
ALKBH4-H
CP
ALKBH4
ALKBH4+CP-L
ALKBH4+CP-M
ALKBH4+CP-H
3'UTR-A14
3' Terminal
5' Terminal
3'UTR-A138
5' Terminal
3' Terminal
CDS-A621
5' Terminal
3' Terminal
CDS-A884
5' Terminal
3' Terminal

**Fig. S26 The viral C2 or CP has no role on METTL14 or ALKBH4 activity.** (**A**) The effect of recombinant C2 on $m^6A$ methylase activity of METTL14 (Tukey's HSD multiple comparisons test: $P < 0.05$). (**B**) The effect of recombinant CP on $m^6A$ demethylase activity of ALKBH4 (Tukey's HSD multiple comparisons test: $P < 0.05$). (**C**) AlphaFold3 prediction of RNA structure of the 25bp RNA strand containing the functional $m^6A$ sites of *Vg*. The orange nucleobases represent the $m^6A$ motif.

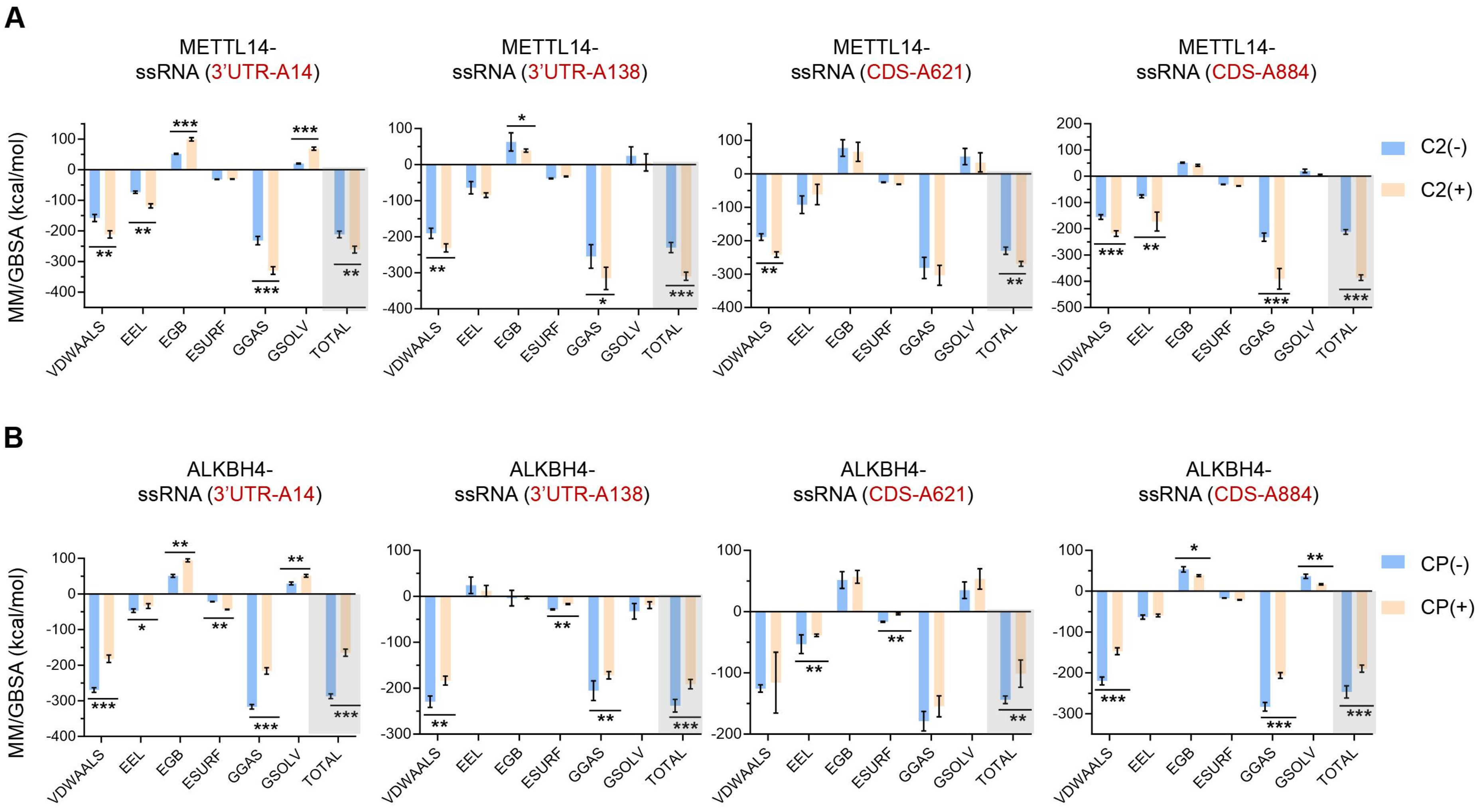

A
METTL14-ssRNA (3'UTR-A14)
METTL14-ssRNA (3'UTR-A138)
METTL14-ssRNA (CDS-A621)
METTL14-ssRNA (CDS-A884)
C2(-)
C2(+)
B
ALKBH4-ssRNA (3'UTR-A14)
ALKBH4-ssRNA (3'UTR-A138)
ALKBH4-ssRNA (CDS-A621)
ALKBH4-ssRNA (CDS-A884)
CP(-)
CP(+)
MM/GBSA (kcal/mol)
VDWAALS
EEL
EGB
ESURF
GGAS
GSOLV
TOTAL

**Fig. S27 Molecular dynamics simulations of binding free energies analyzed by MM/GBSA method.** (**A**) Change in binding free energies of METTL14 to substrate RNAs (*Vg* $m^6A$ site) with C2 (+) or without (-). (**B**) Change in binding free energies of ALKBH4 to substrate RNAs (*Vg* $m^6A$ site) with CP (+) or without (-). Van der Waals energy (VDWAALS), Electrostatic energy (EEL), Polar solvation energy (EGB), Nonpolar solvation energy (ESURF), Gas-phase free energy (GGAS), Solvation free energy (GSOLV), Total binding free energy (TOTAL, $\triangle E_{Bind}$).

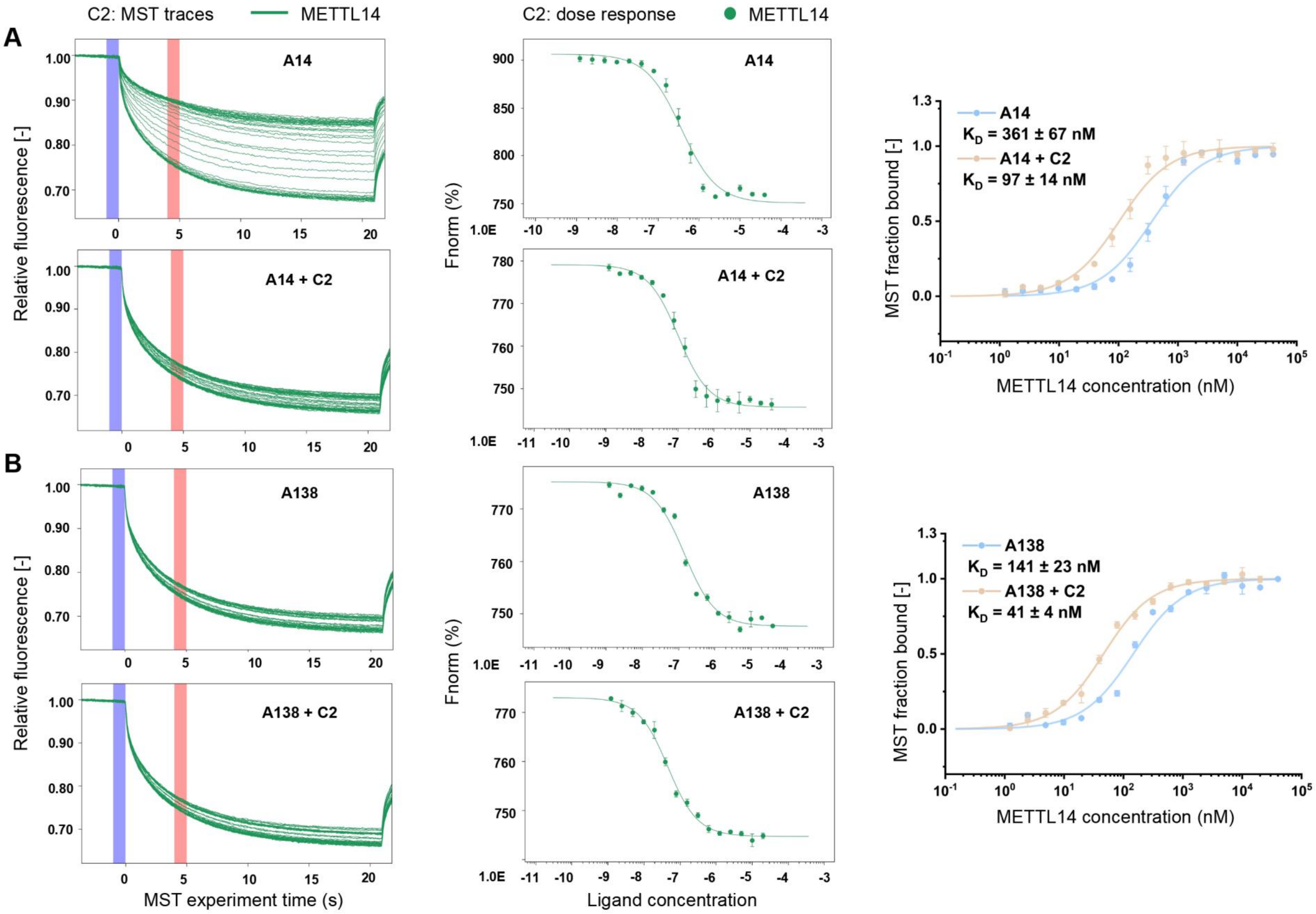
A
B
C2: MST traces
METTL14
C2: dose response
METTL14
A14
A14 + C2
A138
A138 + C2
Relative fluorescence [-]
MST experiment time (s)
Fnorm (%)
Ligand concentration
1.0E
MST fraction bound [-]
METTL14 concentration (nM)
A14
K$_D$ = 361 ± 67 nM
A14 + C2
K$_D$ = 97 ± 14 nM
A138
K$_D$ = 141 ± 23 nM
A138 + C2
K$_D$ = 41 ± 4 nM

**Fig. S28 Changes in MST binding affinities of METTL14 to *Vg* m$^6$A sites in the presence of C2.** (**A**) *Vg* m$^6$A sites: 3UTR-A14. (**B**) *Vg* m$^6$A sites: 3UTR-A138. The vertical bars show the cold fluorescence detected at −1-0 s (blue); hot fluorescence detected at 4-5 s (red).

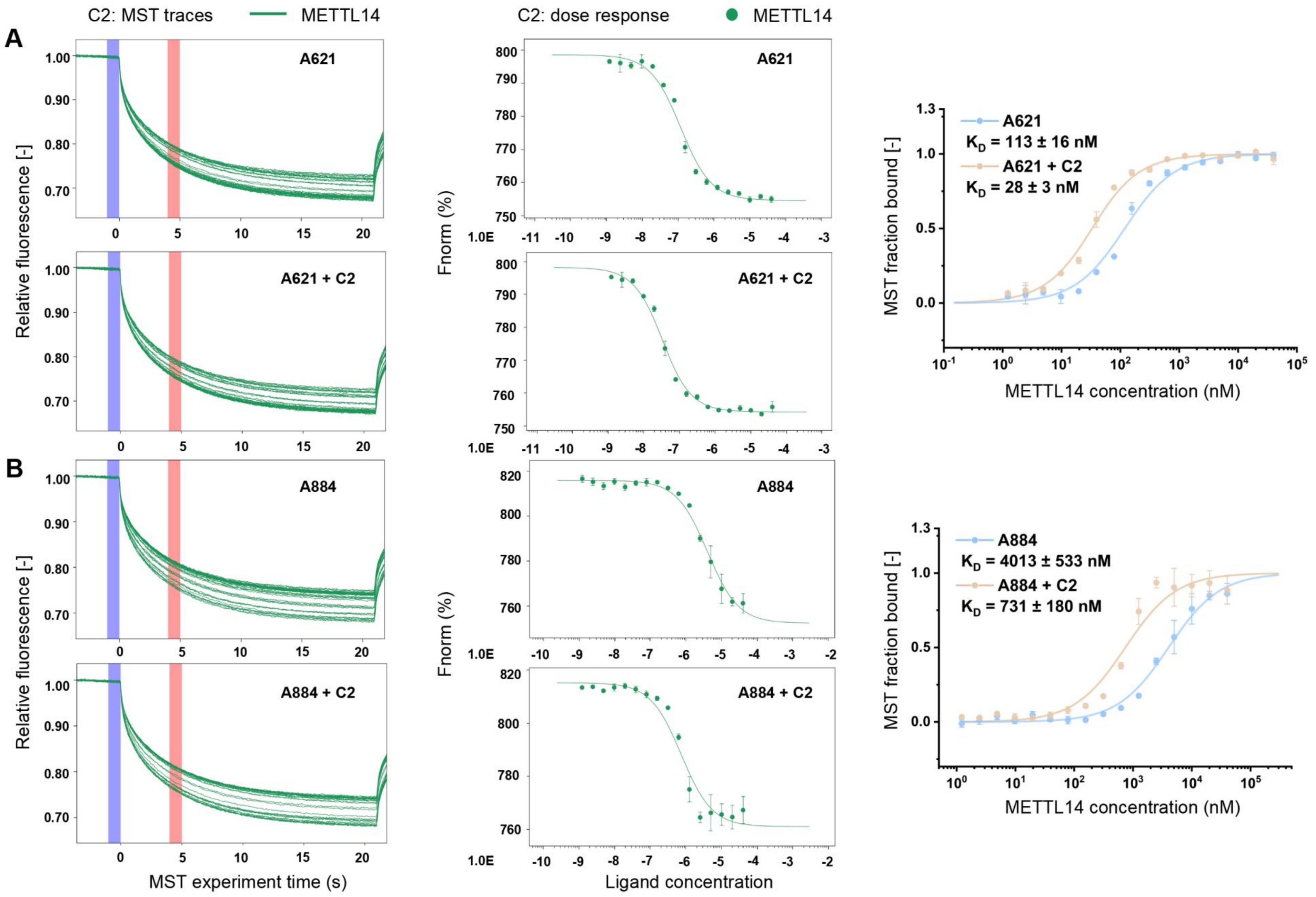
C2: MST traces
METTL14
C2: dose response
METTL14
A
B
A621
A621 + C2
A884
A884 + C2
Relative fluorescence [-]
MST experiment time (s)
Fnorm (%)
Ligand concentration
1.0E
MST fraction bound [-]
METTL14 concentration (nM)
A621
$K_D$ = 113 ± 16 nM
A621 + C2
$K_D$ = 28 ± 3 nM
A884
$K_D$ = 4013 ± 533 nM
A884 + C2
$K_D$ = 731 ± 180 nM

**Fig. S29 Changes in MST binding affinities of METTL14 to *Vg* m$^6$A sites in the presence of C2.** (**A**) *Vg* m$^6$A sites: CDS-621. (**B**) *Vg* m$^6$A sites: CDS-A884. The vertical bars show the cold fluorescence detected at −1-0 s (blue); hot fluorescence detected at 4-5 s (red).

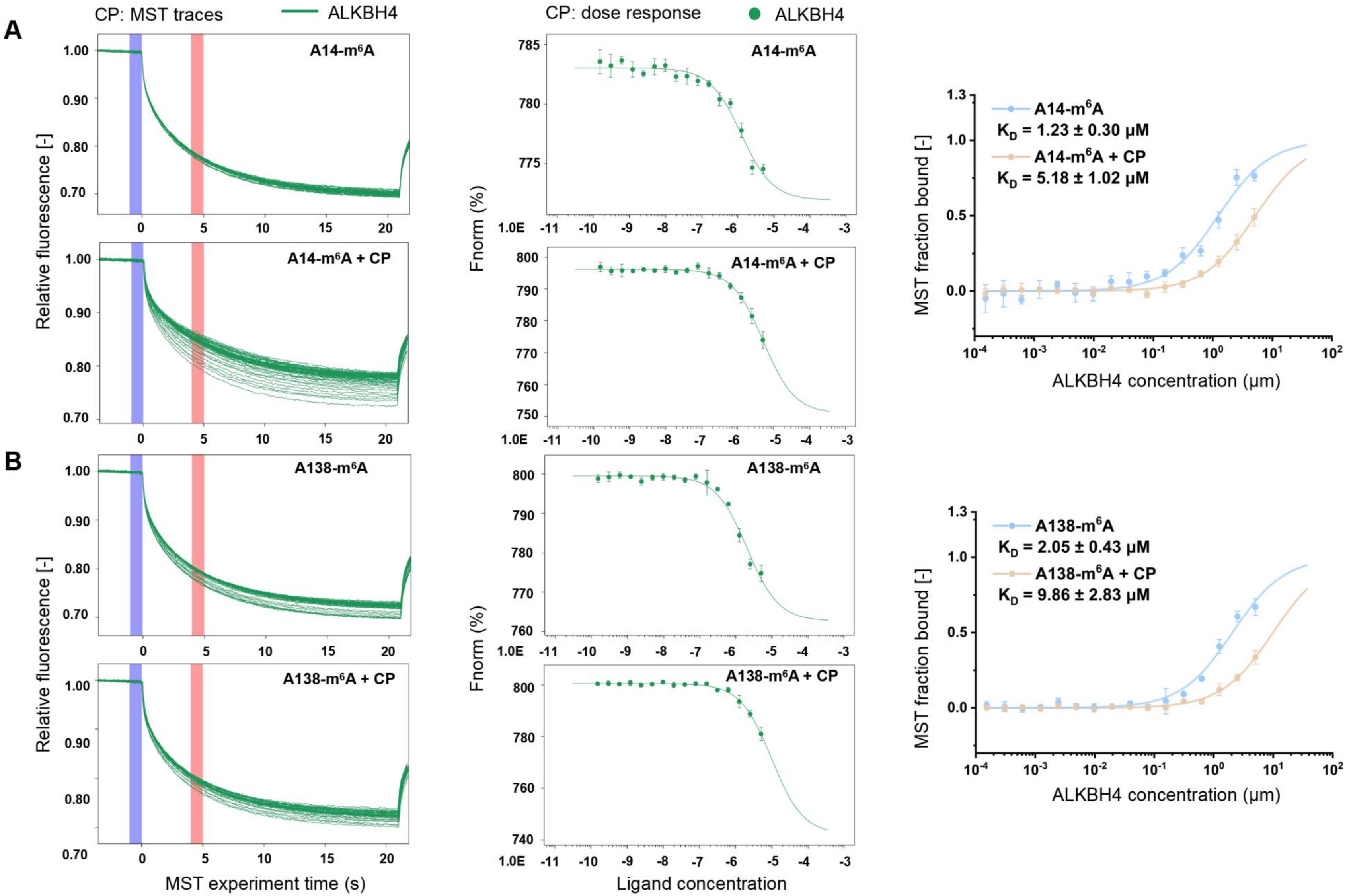

A
B
CP: MST traces
ALKBH4
CP: dose response
ALKBH4
A14-m6A
A14-m6A + CP
A138-m6A
A138-m6A + CP
Relative fluorescence [-]
MST experiment time (s)
Fnorm (%)
Ligand concentration
MST fraction bound [-]
ALKBH4 concentration (μm)
A14-m6A
KD = 1.23 ± 0.30 μM
A14-m6A + CP
KD = 5.18 ± 1.02 μM
A138-m6A
KD = 2.05 ± 0.43 μM
A138-m6A + CP
KD = 9.86 ± 2.83 μM

**Fig. S30 Changes in MST binding affinities of ALKBH4 to *Vg* m$^6$A sites in the presence of CP.** (**A**) *Vg* m$^6$A sites: 3'UTR-A14. (**B**) *Vg* m$^6$A sites: 3'UTR-A138. The vertical bars show the cold fluorescence detected at −1-0 s (blue); hot fluorescence detected at 4-5 s (red).

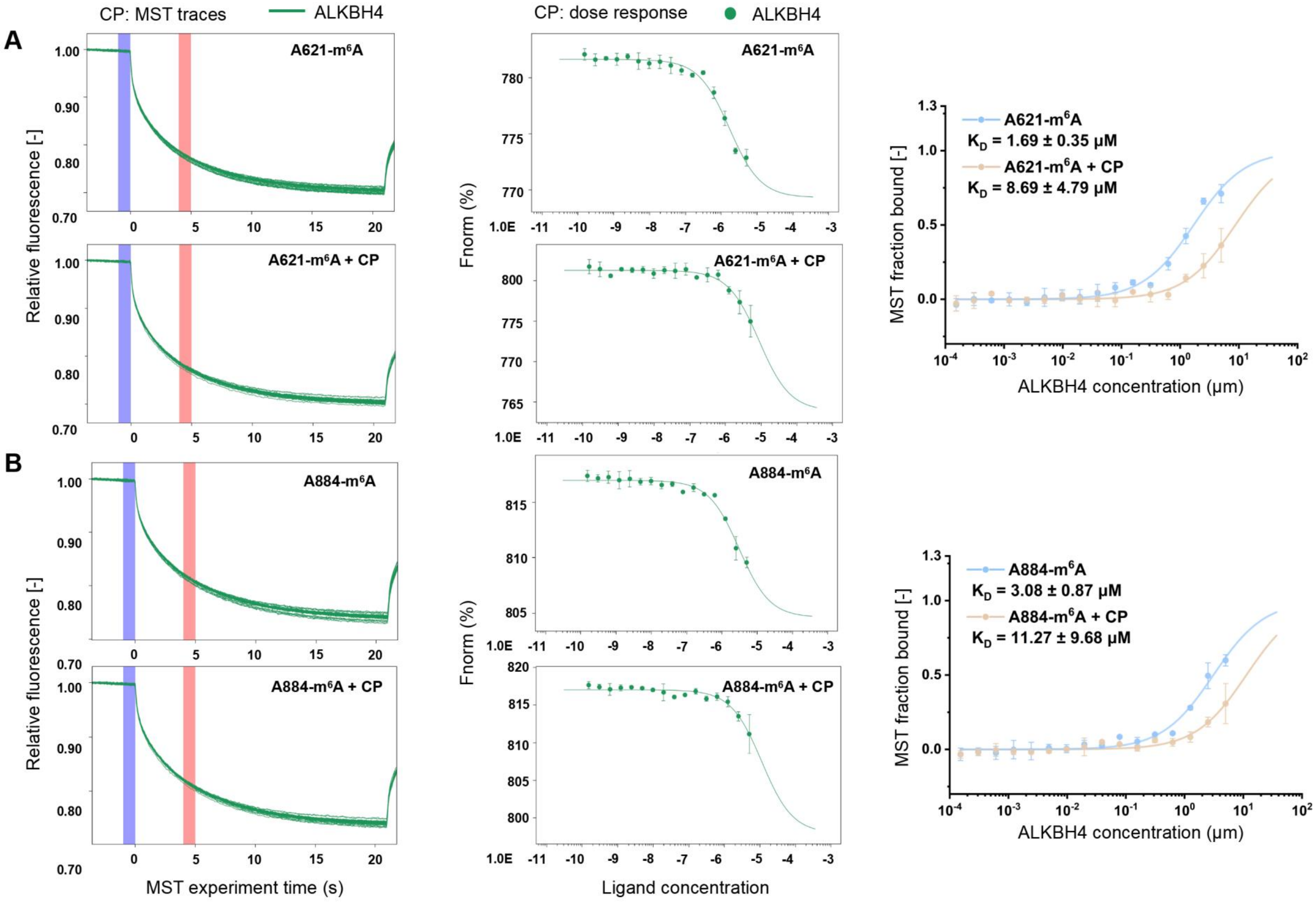

A
B
CP: MST traces
ALKBH4
CP: dose response
ALKBH4
A621-m6A
A621-m6A + CP
A884-m6A
A884-m6A + CP
Relative fluorescence [-]
MST experiment time (s)
Fnorm (%)
Ligand concentration
A621-m6A
KD = 1.69 ± 0.35 μM
A621-m6A + CP
KD = 8.69 ± 4.79 μM
A884-m6A
KD = 3.08 ± 0.87 μM
A884-m6A + CP
KD = 11.27 ± 9.68 μM
MST fraction bound [-]
ALKBH4 concentration (μm)

**Fig. S31 Changes in MST binding affinities of ALKBH4 to *Vg* m$^6$A sites in the presence of CP.** (**A**) *Vg* m$^6$A sites: CDS-A621. (**B**) *Vg* m$^6$A sites: CDS-A884. The vertical bars show the cold fluorescence detected at −1-0 s (blue); hot fluorescence detected at 4-5 s (red).

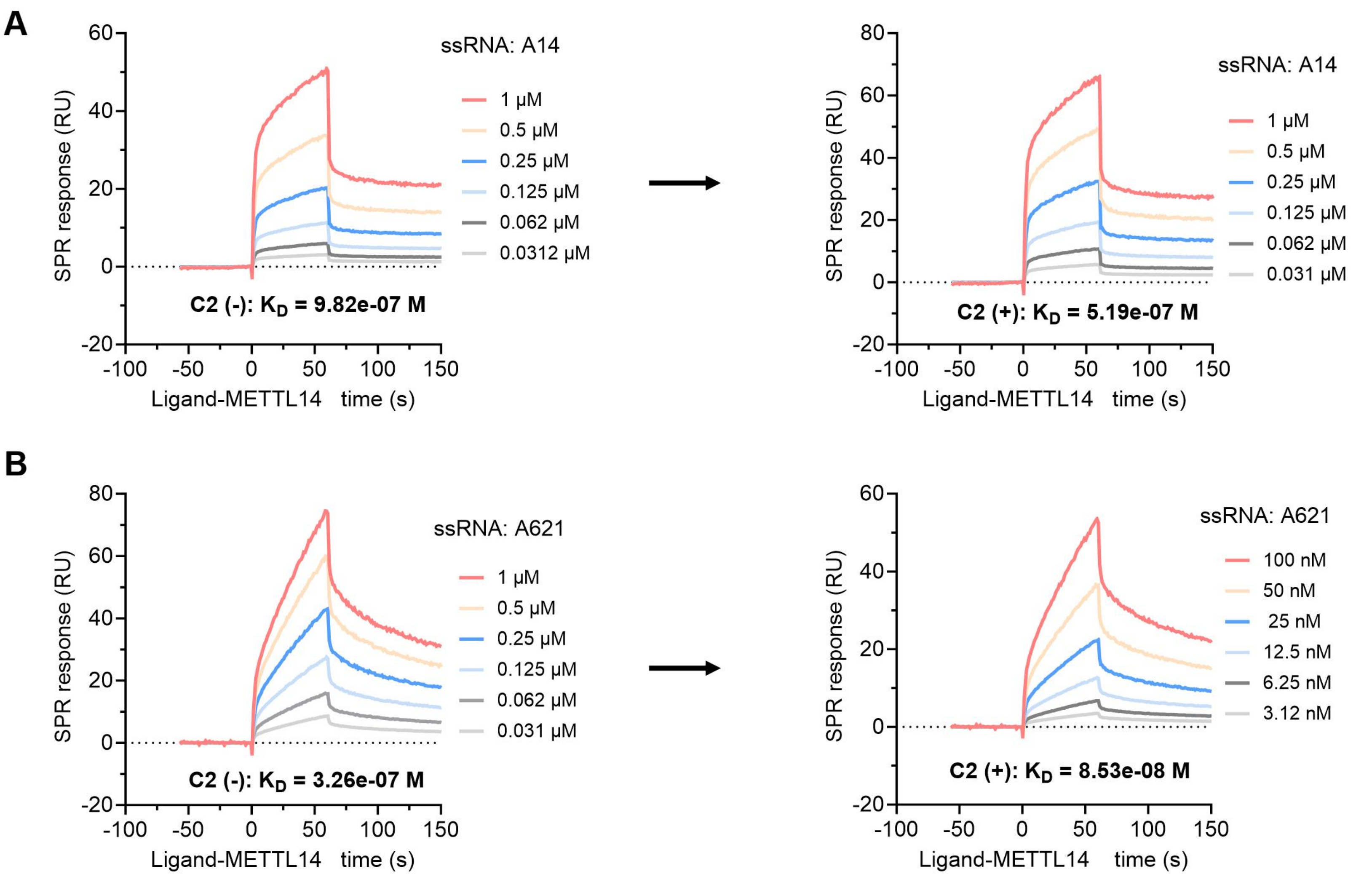
A
ssRNA: A14
1 μM
0.5 μM
0.25 μM
0.125 μM
0.062 μM
0.0312 μM
C2 (-): K_D = 9.82e-07 M
SPR response (RU)
Ligand-METTL14 time (s)
ssRNA: A14
1 μM
0.5 μM
0.25 μM
0.125 μM
0.062 μM
0.031 μM
C2 (+): K_D = 5.19e-07 M
B
ssRNA: A621
1 μM
0.5 μM
0.25 μM
0.125 μM
0.062 μM
0.031 μM
C2 (-): K_D = 3.26e-07 M
ssRNA: A621
100 nM
50 nM
25 nM
12.5 nM
6.25 nM
3.12 nM
C2 (+): K_D = 8.53e-08 M

**Fig. S32 The SPR assay of binding affinities of METTL14 to *Vg* m$^6$A sites in the presence of viral C2 protein.** (**A**) Changes in binding affinities of METTL14 to substrate RNA 3'UTR-A14 with C2 (+) or without (-). (**B**) Changes in binding affinities of METTL14 to substrate RNA CDS-A621 with C2 (+) or without (-).

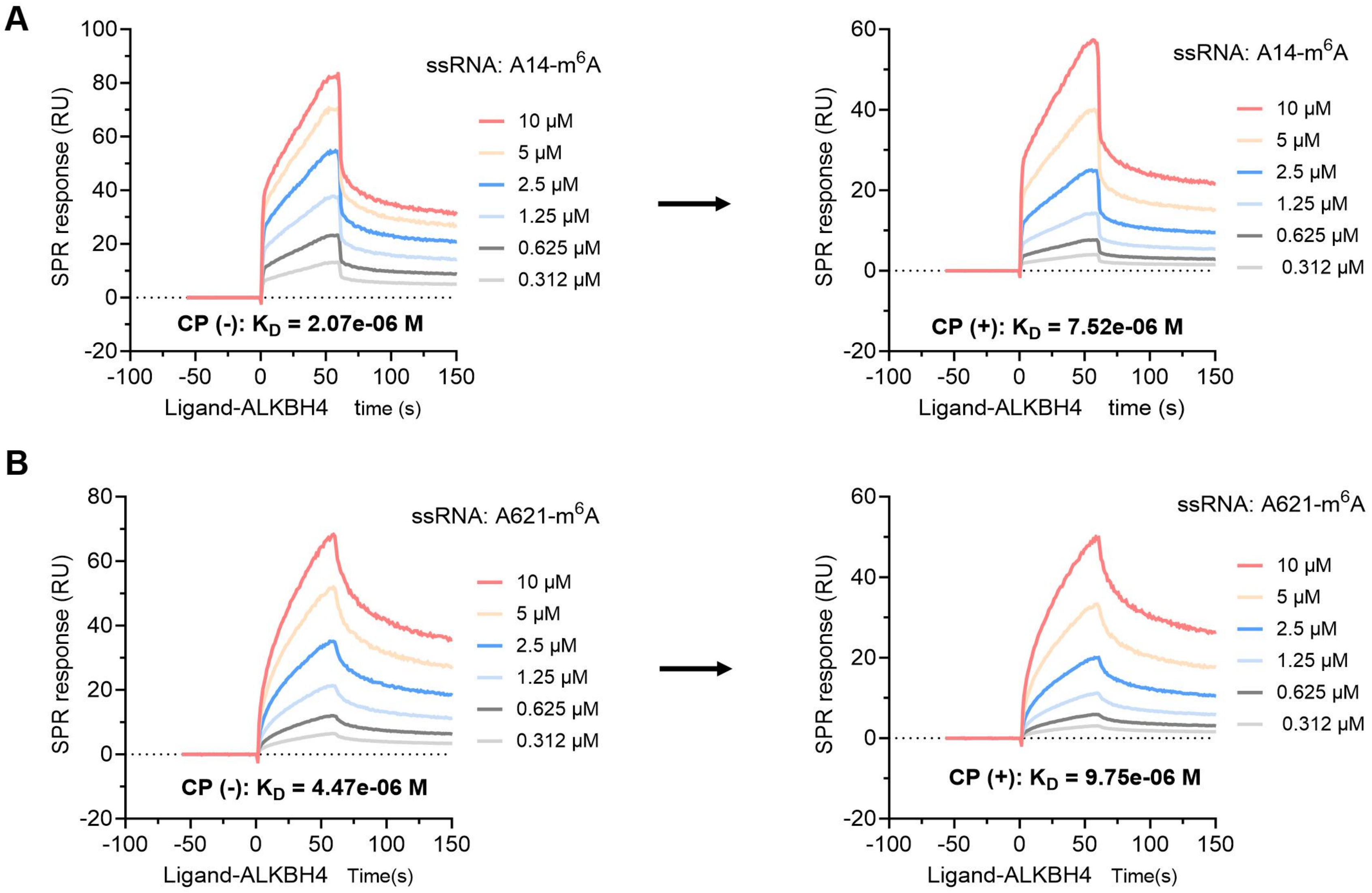
A
ssRNA: A14-m6A
10 μM
5 μM
2.5 μM
1.25 μM
0.625 μM
0.312 μM
SPR response (RU)
CP (-): K_D = 2.07e-06 M
Ligand-ALKBH4 time (s)
ssRNA: A14-m6A
10 μM
5 μM
2.5 μM
1.25 μM
0.625 μM
0.312 μM
SPR response (RU)
CP (+): K_D = 7.52e-06 M
Ligand-ALKBH4 time (s)
B
ssRNA: A621-m6A
10 μM
5 μM
2.5 μM
1.25 μM
0.625 μM
0.312 μM
SPR response (RU)
CP (-): K_D = 4.47e-06 M
Ligand-ALKBH4 Time(s)
ssRNA: A621-m6A
10 μM
5 μM
2.5 μM
1.25 μM
0.625 μM
0.312 μM
SPR response (RU)
CP (+): K_D = 9.75e-06 M
Ligand-ALKBH4 Time(s)

**Fig. S33 The SPR assay of binding affinities of ALKBH4 to *Vg* m$^6$A sites in the presence of viral CP protein.** (**A**) Changes in binding affinities of ALKBH4 to substrate RNA 3'UTR-A14 with CP (+) or without (-). (**B**) Changes in binding affinities of ALKBH4 to substrate RNA CDS-A621 with CP (+) or without (-).

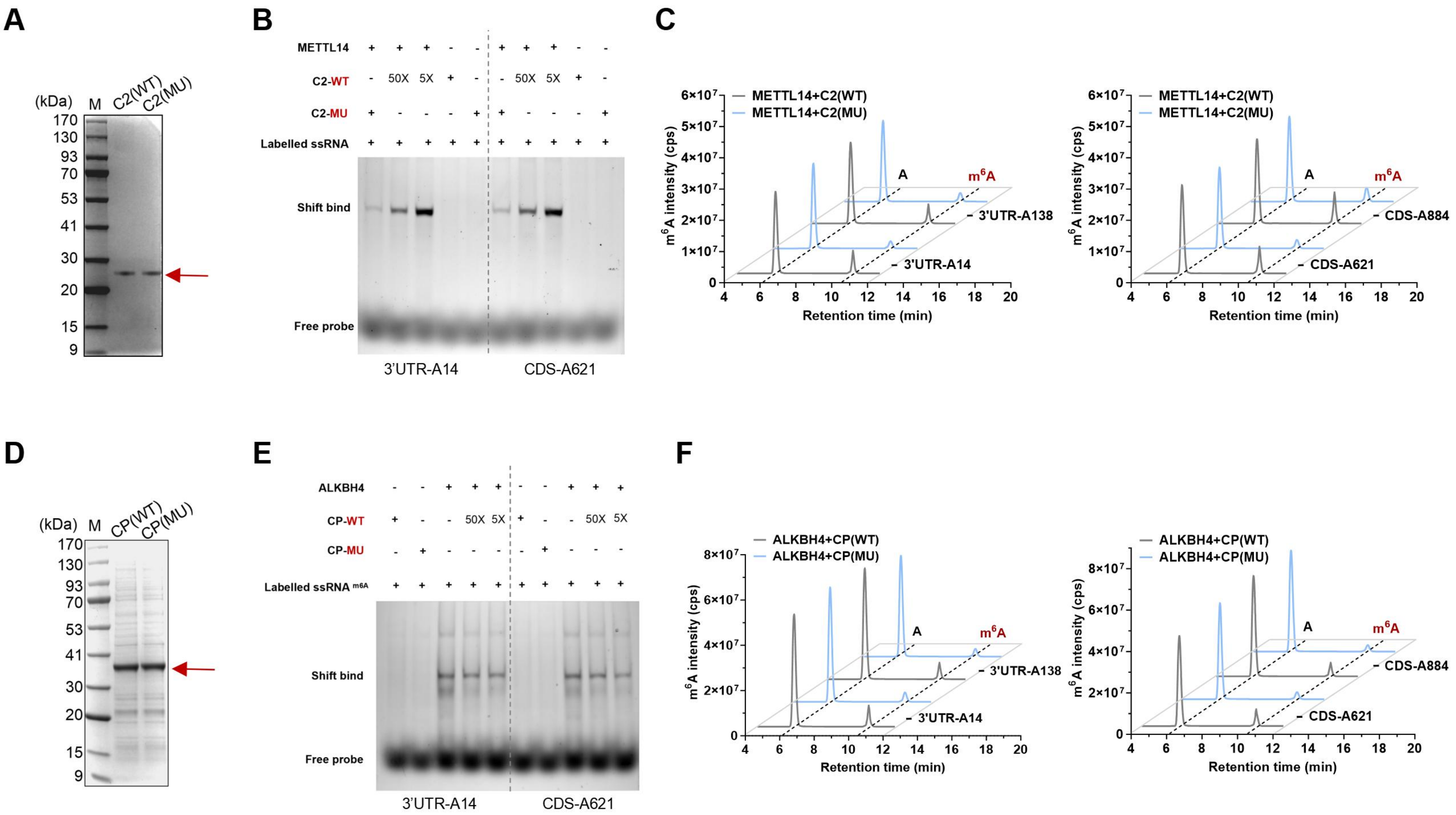
A
(kDa)
M
C2(WT)
C2(MU)
170
130
93
70
53
41
30
20
15
9
B
METTL14
C2-WT
C2-MU
Labelled ssRNA
50X
5X
Shift bind
Free probe
3'UTR-A14
CDS-A621
C
METTL14+C2(WT)
METTL14+C2(MU)
m6A intensity (cps)
Retention time (min)
A
m6A
3'UTR-A138
3'UTR-A14
CDS-A884
CDS-A621
D
(kDa)
M
CP(WT)
CP(MU)
E
ALKBH4
CP-WT
CP-MU
Labelled ssRNA m6A
Shift bind
Free probe
3'UTR-A14
CDS-A621
F
ALKBH4+CP(WT)
ALKBH4+CP(MU)
m6A intensity (cps)
Retention time (min)
3'UTR-A138
3'UTR-A14
CDS-A884
CDS-A621

**Fig. S34 The presence of C2 or CP alters binding specificity of METTL14 or ALKBH4 to the substrate *Vg* m$^6$A sites, leading to an increased m$^6$A modification on *Vg*.** (**A**) Heterologous expression of recombinant C2 protein in *E. coli* as detected by SDS-PAGE gel. The mutated C2(MU) protein was generated by mutating its binding bounds in the METTL14-ssRNA-C2 interaction. Protein marker (M). (**B**) EMSA assay of METTL14 binding to *Vg* m$^6$A sites in the presence of wild-type C2 (WT) or its mutated protein (MU). A 25bp-length ssRNA covering the functional m$^6$A site was used as a probe. (**C**) UPLC-MS/MS analysis of the effect of C2 and mutated C2 protein on m$^6$A modification of *Vg* by METTL14. (**D**) Heterologous expression of recombinant CP protein in *E. coli* as detected by SDS-PAGE gel. The mutated CP(MU) protein was generated by mutating its binding bounds in the METTL14-ssRNA-C2 interaction. Protein marker (M). (**E**) EMSA assay of ALKBH4 binding to *Vg* m$^6$A sites in the presence of wild-type CP (WT) or its mutated protein (MU). A 25bp-length ssRNA covering the functional m$^6$A site labelled with m$^6$A on A site was used as a probe. (**F**) UPLC-MS/MS analysis of the effect of CP and mutated CP protein on m$^6$A modification of *Vg* by ALKBH4. The mutated CP-MU protein was generated by mutating its binding bounds in the ALKBH4-ssRNA-CP interaction.

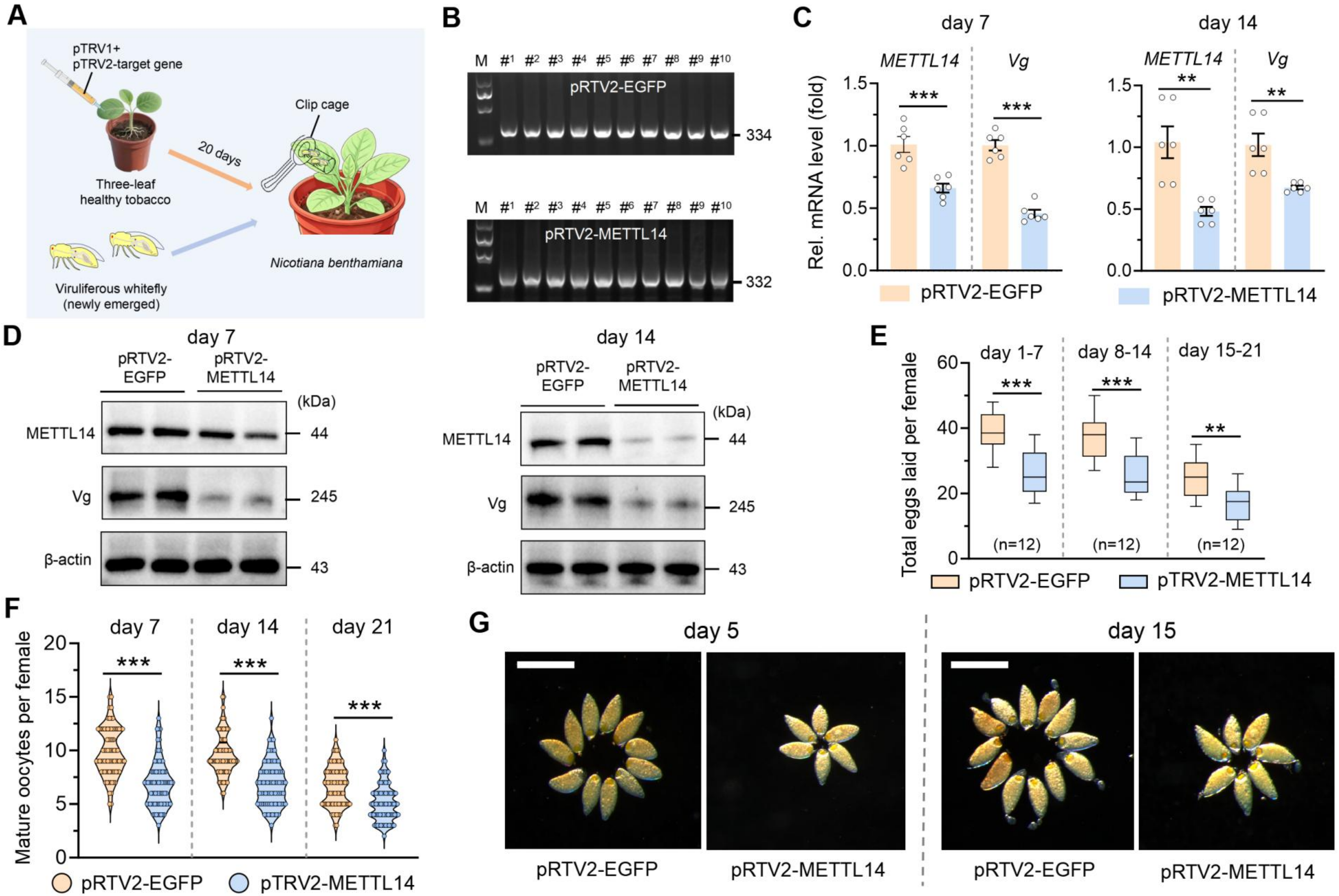
A
pTRV1+
pTRV2-target gene
20 days
Clip cage
Three-leaf
healthy tobacco
Viruliferous whitefly
(newly emerged)
Nicotiana benthamiana
B
M #1 #2 #3 #4 #5 #6 #7 #8 #9 #10
pRTV2-EGFP
334
pRTV2-METTL14
332
C
day 7
day 14
METTL14
Vg
Rel. mRNA level (fold)
pRTV2-EGFP
pRTV2-METTL14
D
day 7
day 14
pRTV2-EGFP
pRTV2-METTL14
(kDa)
METTL14
44
Vg
245
β-actin
43
E
day 1-7
day 8-14
day 15-21
Total eggs laid per female
(n=12)
pRTV2-EGFP
pTRV2-METTL14
F
day 7
day 14
day 21
Mature oocytes per female
pRTV2-EGFP
pTRV2-METTL14
G
day 5
day 15
pRTV2-EGFP
pRTV2-METTL14

**Fig. S35 VIGS-mediated inhibition of METTL14-C2 binding interaction returns to reproductive cost of neonicotinoids resistance in the vector.** (**A**) Flow chart for constructing the VIGS-plants *Nicotiana benthamiana* and its use in inhibiting METTL14-C2 interaction through persistently silencing of METTL14 in the viruliferous vector. (**B**) PCR products amplified using cDNA from pTRV2-EGFP (334-bp dsEGFP fragments) and pTRV2-METTL14 (332-bp dsMETTL14 fragments) tobacco leaves treated for 15 days. M, marker. lanes #1-10, PCR products from 10 tobacco seedlings. (**C**) qPCR analysis of mRNA level of *METTL14* and *Vg* in female adults feeding on pTRV2-METTL14 tobacco leaves for 7 and 14 days (n = 3, Student's *t* test: $^{**}P < 0.01$, $^{***}P < 0.001$). (**D**) Western blot analysis of protein level of METTL14 and Vg in female adults feeding on pTRV2-METTL14 tobacco leaves for 7 and 14 days. (**E-G**) Phenotypic observation of female fecundity and mature oocytes of vector adults feeding on pTRV2-METTL14 tobacco leaves (n ≥ 60, Student's *t* test: $^{***}P < 0.001$). Scar bar, 200 μm.

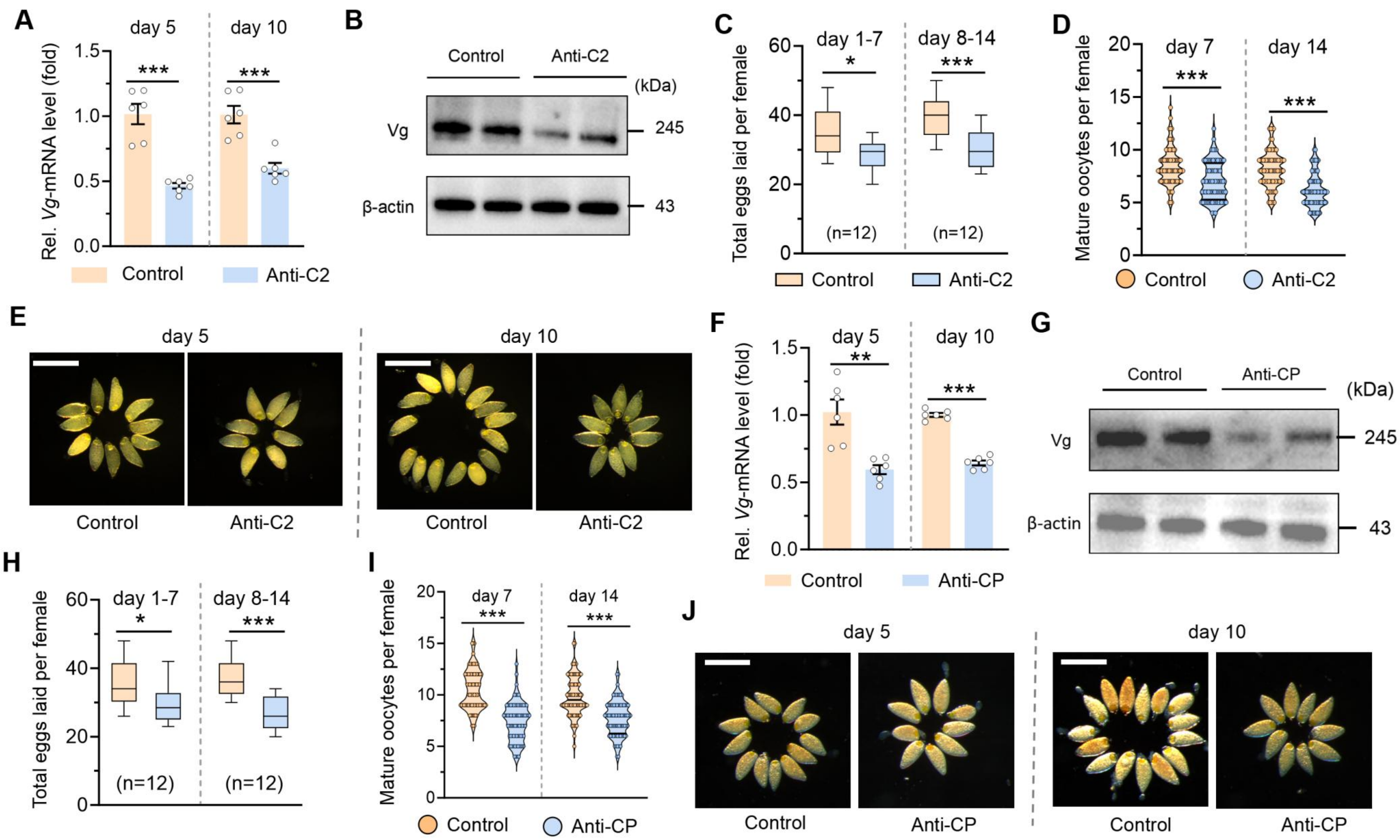

A
day 5
day 10
***
***
Rel. Vg-mRNA level (fold)
1.5
1.0
0.5
0.0
Control
Anti-C2
B
Control
Anti-C2
(kDa)
Vg
245
β-actin
43
C
day 1-7
day 8-14
*
***
Total eggs laid per female
60
40
20
0
(n=12)
(n=12)
Control
Anti-C2
D
day 7
day 14
***
***
Mature oocytes per female
20
15
10
5
0
Control
Anti-C2
E
day 5
day 10
Control
Anti-C2
Control
Anti-C2
F
day 5
day 10
**
***
Rel. Vg-mRNA level (fold)
1.5
1.0
0.5
0.0
Control
Anti-CP
G
Control
Anti-CP
(kDa)
Vg
245
β-actin
43
H
day 1-7
day 8-14
*
***
Total eggs laid per female
60
40
20
0
(n=12)
(n=12)
I
day 7
day 14
***
***
Mature oocytes per female
20
15
10
5
0
Control
Anti-CP
J
day 5
day 10
Control
Anti-CP
Control
Anti-CP

**Fig. S36 Inhibition of C2 or CP leads to reproductive cost of neonicotinoids resistance in the vector.** (**A**-**B**) Transcript (**A**) and protein (**B**) level of C2 and Vg in female adults feeding on anti-C2 antibody. (**C**-**E**) Average female fecundity (**C**) and mature oocytes (**D, E**) of female adults feeding on anti-C2 antibody. All data were presented as the mean ± SEM of at least three biological replicates. Scar bar, 200 μm. Data were analyzed by Student's *t* test: *$P$ < 0.05, **$P$ < 0.01, ***$P$ < 0.001. (**F**-**G**) Transcript (**F**) and protein (**G**) level of C2 and Vg in female adults feeding on anti-CP antibody. (**H**-**J**) Average female fecundity (**H**) and mature oocytes (**I**, **J**) of female adults feeding on anti-CP antibody. All data were presented as the mean ± SEM of at least three biological replicates. Scar bar, 200 μm. Data were analyzed by Student's *t* test: *$P$ < 0.05, **$P$ < 0.01, ***$P$ < 0.001.

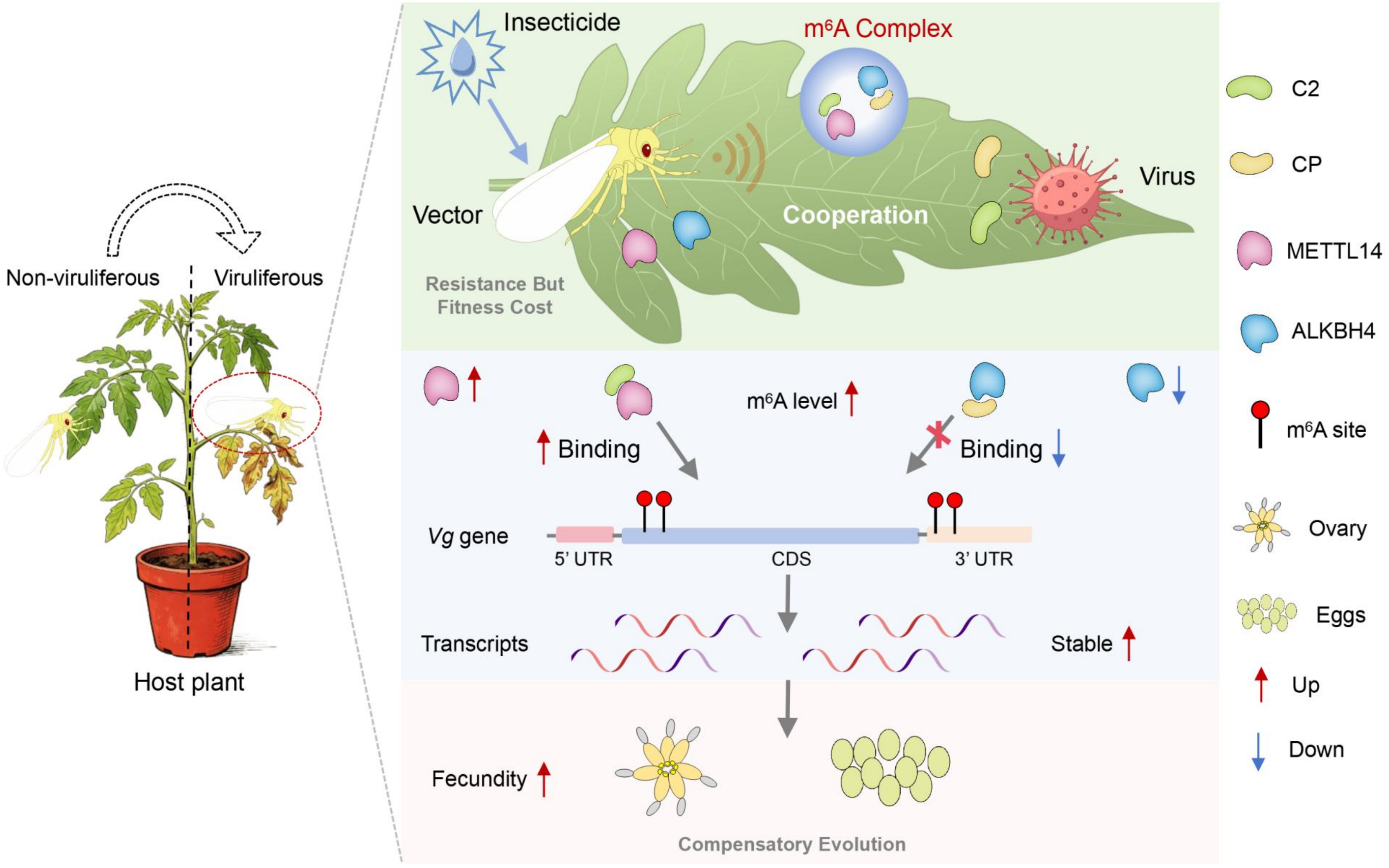

Non-viruliferous
Viruliferous
Host plant
Insecticide
m6A Complex
Vector
Cooperation
Virus
Resistance But
Fitness Cost
m6A level
Binding
Binding
Vg gene
5' UTR
CDS
3' UTR
Transcripts
Stable
Fecundity
Compensatory Evolution
C2
CP
METTL14
ALKBH4
m6A site
Ovary
Eggs
Up
Down

**Fig. S37 A proposed working model for epigenetic regulation of insect vector-virus beneficial relationship.** The plant virus TYLCV acts as a cooperative partner to rescue the fitness costs of neonicotinoid resistance by employing $m^6A$ pathway in the vector *B. tabaci*. Specifically, the viral protein C2 and CP separately binds to the writer METTL14 and eraser ALKBH4, remodeling their structural conformation and RNA binding affinity to Vg $m^6A$ sites for $m^6A$ modification —a process largely responsible for this compensatory evolution. Thus, virus can co-opt vectors' epigenetic machinery to establish beneficial virus-vector relationships, offering a new paradigm for their coevolution.

**Table S1** Source of the field vectors *B. tabaci* MED

| Population name | Sampling location | Longitude and latitude | Collection date | Host plant | Sample type |
|---|---|---|---|---|---|
| SY1 | Sanya, Hainan | 108°56′E, 18°9′N | Mar. 2023 | Eggplant | field |
| SY2 | Sanya, Hainan | 108°56′E, 18°9′N | Apr. 2023 | Hami melon | field |
| SY3 | Sanya, Hainan | 108°56′E, 18°9′N | Apr. 2023 | Eggplant | field |
| HK | Haikou, Hainan | 110°19′E, 19°14′N | Apr. 2023 | Tomato | field |
| YY | Yueyang, Hunan | 112°93′E, 29°46′N | Apr. 2023 | Chili | field |
| JN | Jinan, Shandong | 117°40′E, 36°64′N | Jun. 2023 | Tomato | field |
| YM | Yuanmou, Yunnan | 101°52′E, 25°47′N | Jul. 2023 | Sweet potato | field |
| BJ1 | Beijing | 116°33′E, 39°96′N | Aug. 2023 | Eggplant | field |
| BJ2 | Beijing | 116°33′E, 39°96′N | Sep. 2023 | Eggplant | field |
| YC | Yuncheng, Shangxi | 110°97′E, 35°03′N | Aug. 2023 | Eggplant | field |
| FY | Fuyang, Anhui | 115°56′E, 32°56′N | Sep. 2023 | Soybean | field |
| TH | Taihe, Anhui | 115°38′E, 33°14′N | Sep. 2023 | White gourd | field |
| CS | Changsha, Hunan | 113°05′E, 28°12′N | Oct. 2023 | Chili | field |
| HZ | Hangzhou, Zhejiang | 118°21′E 29°11′N | Oct. 2023 | Eggplant | field |
| YT | Yantai, Shandong | 121°27′E, 37°48′N | Nov. 2023 | Tomato | field |
| KS | Kashi, Xinjiang | 75°39′E, 39°28′N | Oct. 2024 | Cotton | field |
| YC | Yuncheng, Shangxi | 110°97′E, 35°03′N | Aug. 2024 | Eggplant | field |
| JN | Jinan, Shandong | 117°40′E, 36°64′N | Jun. 2024 | Cucumber | field |
| BD | Baoding, Hebei | 115°27′E, 38°52′N | Apr. 2024 | Tomato | field |
| TH | Taihe, Anhui | 115°38′E, 33°14′N | Mar. 2024 | Tomato | field |
| HF | Hefei, Anhui | 117°13′E, 31°49′N | Nov. 2024 | Chili | field |
| DQ | Deqing, Zhejiang | 119°58′E, 30°23′N | Sep. 2024 | Eggplant | field |
| HZ | Hangzhou, Zhejiang | 118°21′E 29°11′N | Sep. 2024 | Eggplant | field |
| WH | Wuhan, Hubei | 113°53′E, 29°58′N | Oct. 2024 | Eggplant | field |
| HK | Haikou, Hainan | 110°19′E, 19°14′N | Mar. 2024 | Tomato | field |
| SY | Sanya, Hainan | 108°56′E, 18°9′N | Mar. 2024 | Eggplant | field |
| SH | Shanghai | 121°28′E, 31°13′N | Jun. 2024 | Eggplant | field |
| CS | Changsha, Hunan | 113°05′E, 28°12′N | Oct. 2024 | Chili | field |
| KS | Kashi, Xinjiang | 75°39′E, 39°28′N | Oct. 2025 | Cotton | field |
| YC | Yuncheng, Shangxi | 110°97′E, 35°03′N | Aug. 2025 | Eggplant | field |
| JN | Jinan, Shandong | 117°40′E, 36°64′N | Jun. 2025 | Cucumber | field |
| TJ | Tianjin | 117°04′E, 39°38′N | Sep. 2025 | Eggplant | field |
| XX | Xinxiang, Henan | 113°55′E, 35°18′N | Sep. 2025 | Pumpkin | field |
| DQ | Deqing, Zhejiang | 119°58′E, 30°23′N | Aug. 2025 | Pumpkin | field |
| HZ | Hangzhou, Zhejiang | 118°21′E 29°11′N | Sep. 2025 | Eggplant | field |
| WH | Wuhan, Hubei | 113°53′E, 29°58′N | Oct. 2025 | Melon | field |
| HK | Haikou, Hainan | 110°19′E, 19°14′N | Mar. 2025 | Eggplant | field |
| CS | Changsha, Hunan | 113°05′E, 28°12′N | Oct. 2025 | Eggplant | field |
| XSBN | Xishuangbanna, Yunnan | 100°47′E, 22°0′N | Mar. 2025 | Eggplant | field |

**Table S2** Imidacloprid resistance and TYLCV infection rate in field vector *B. tabaci* MED

| Population name | Year | Slope [a] | $LC_{50}$ (mg/L, 95%FL [b]) | $χ^2$ [c] | $RR_{50}$ [d] (fold) | Resistance level [e] | TYLCV [f] (%) |
|---|---|---|---|---|---|---|---|
| Reference [g] | - | - | 0.99 | - | 1.00 | Sensitive | - |
| SY1 | | 1.19 | 1915.49 (1038-6556) | 2.90 | 1934.84 | Extremely high | 100.00 |
| SY2 | | 1.08 | 1884.49 (1000-6819) | 1.41 | 1903.53 | Extremely high | 100.00 |
| SY3 | | 0.80 | 657.33 (382-1860) | 3.20 | 663.97 | Very high | 0.00 |
| HK | | 0.86 | 551.05 (332-1346) | 2.89 | 556.62 | Very high | 100.00 |
| YY | | 1.25 | 2041.98 (1503-3008) | 3.87 | 2062.61 | Extremely high | 100.00 |
| JN | | 1.57 | 55.18 (19-92) | 1.16 | 55.74 | High | 0.00 |
| YM | | 0.57 | 471.46 (191-998) | 3.89 | 476.22 | Very high | 30.00 |
| BJ1 | 2023 | 1.79 | 385.66 (300-489) | 3.80 | 389.56 | Very high | 0.00 |
| BJ2 | | 1.43 | 165.19 (77-259) | 4.55 | 166.86 | High | 0.00 |
| YC | | 1.00 | 3353.69 (1892-11376) | 0.25 | 3387.57 | Extremely high | 100.00 |
| FY | | 1.01 | 124.86 (58-196) | 0.80 | 126.12 | High | 0.00 |
| TH | | 0.84 | 159.41 (73-253) | 1.46 | 161.02 | High | 0.00 |
| CS | | 0.87 | 1970.38 (1214-4570) | 0.50 | 1990.28 | Extremely high | 0.00 |
| HZ | | 0.69 | 759.00 (468-1408) | 0.39 | 766.67 | Very high | 0.00 |
| YT | | 0.79 | 2733.72 (1792-4832) | 0.46 | 2761.33 | Extremely high | 70.00 |
| KS | | 0.87 | 9917.91 (4381-88744) | 4.23 | 10018.09 | Extremely high | 100.00 |
| YC | | 1.16 | 1258.56 (919-1866) | 0.86 | 1271.27 | Extremely high | 100.00 |
| JN | | 0.79 | 2733.73 (1792-4832) | 0.61 | 2761.34 | Extremely high | 100.00 |
| BD | | 0.89 | 525.33 (356-776) | 0.51 | 530.64 | Very high | 70.00 |
| TH | | 0.84 | 159.41 (73-253) | 1.46 | 161.02 | High | 50.00 |
| HF | | 1.03 | 132.30 (63-207) | 0.82 | 133.64 | High | 30.00 |
| DQ | 2024 | 0.71 | 826.53 (510-1570) | 0.39 | 834.88 | Very high | 100.00 |
| HZ | | 1.04 | 501.21 (361-743) | 1.32 | 506.27 | Very high | 100.00 |
| WH | | 0.94 | 8565.58 (5158-22768) | 0.28 | 8652.10 | Extremely high | 100.00 |
| HK | | 1.01 | 435.04 (310-683) | 0.87 | 439.43 | Very high | 100.00 |
| SY | | 1.23 | 478.09 (361-652) | 0.61 | 482.92 | Very high | 55.00 |
| SH | | 0.76 | 951.05 (576-2313) | 0.39 | 960.66 | Very high | 100.00 |
| CS | | 1.14 | 3353.90 (2314-5931) | 1.54 | 3387.78 | Extremely high | 100.00 |
| KS | | -0.466 | > 40000 (-) | 15.9 | 40404.04 | Extremely high | 100.00 |
| YC | | 1.65 | 514.82 (357-713) | 1.25 | 520.02 | Very high | 100.00 |
| JN | | 1.33 | 53.37 (7-120) | 17.46 | 53.91 | High | 30.00 |
| TJ | | 1.56 | 257.09 (198-324) | 0.52 | 259.69 | Very high | 70.00 |
| XX | | 0.57 | 2061.40 (929-100027) | 1.73 | 2082.22 | Extremely high | 100.00 |
| DQ | 2025 | 1.46 | 309.13 (216-419) | 1.32 | 31.25 | High | 60.00 |
| HZ | | 1.68 | 432.54 (180-796) | 9.77 | 436.91 | Very high | 100.00 |
| WH | | 1.54 | 6496.21 (3994-7215) | 1.25 | 6561.83 | Extremely high | 100.00 |
| HK | | 2.32 | 106.52 (20-193) | 6.79 | 107.60 | High | 80.00 |
| CS | | 1.53 | 3365.86 (1647-7520) | 13.22 | 3399.86 | Extremely high | 100.00 |
| XSBN | | 1.47 | 486.64 (374-617) | 1.73 | 491.56 | Very high | 70.00 |

[a] Slope, calculated by Probit and Logit Analysis.

[b] FL, concentration of insecticide killing 50% of adults and its 95% fiducial limits.

[c] $\chi^2$, Chi-square testing linearity of dose-mortality responses.

[d] $RR_{50}$, resistance ratio = the $LC_{50}$ value of the field population divided by the $LC_{50}$ value of the sensitive reference strain.

[e] extremely high resistance ($RR_{50} \geq 1000$), very high resistance ($1000 < RR_{50} \leq 300$), high resistance ($300 < RR_{50} \leq 30$ )

[f] a total of 20 whitefly individuals for each population were used for TYLCV detection.

[g] the sensitive reference strain from the study by Wang et al., (82).

**Table S3** Thiamethoxam resistance and TYLCV infection rate in field *B. tabaci* MED

| Population name | Year | Slope [a] | $LC_{50}$ (mg/L, 95%FL [b]) | $\chi^2$ [c] | $RR_{50}$ [d] (fold) | Resistance level [e] | TYLCV [f] (%) |
|---|---|---|---|---|---|---|---|
| Reference [g] | - | - | 1.19 | - | 1.00 | Sensitive | - |
| SY1 | 2023 | 1.42 | 1653.18 (1184-2673) | 3.81 | 1389.23 | Extremely high | 100.00 |
| SY2 | | 1.24 | 6498.12 (3448-24716) | 3.32 | 5460.61 | Extremely high | 100.00 |
| SY3 | | 1.07 | 435.96 (292-627) | 0.83 | 366.35 | Very high | 0.00 |
| HK | | 1.03 | 470.94 (287-730) | 0.45 | 395.75 | Very high | 100.00 |
| YY | | 2.08 | 3752.09 (2000-7513) | 12.0 | 3153.02 | Extremely high | 100.00 |
| JN | | 2.37 | 319.73 (185-524) | 12.20 | 268.68 | High | 0.00 |
| YM | | 2.42 | 225.41 (85-410) | 14.90 | 189.42 | High | 30.00 |
| BJ1 | | 2.01 | 801.48 (567-1177) | 4.83 | 673.51 | Very high | 0.00 |
| BJ2 | | 1.65 | 167.33 (120-216) | 3.99 | 140.61 | High | 0.00 |
| YC | | 1.18 | 2897.34 (1730-8285) | 1.02 | 2434.74 | Extremely high | 100.00 |
| FY | | 1.03 | 66.40 (24-113) | 0.43 | 55.80 | High | 0.00 |
| TH | | 1.02 | 99.53 (53-148) | 1.30 | 83.64 | High | 0.00 |
| CS | | 0.87 | 681.98 (463-1079) | 0.44 | 573.09 | Very high | 0.00 |
| HZ | | 0.61 | 751.33 (439-1639) | 0.43 | 631.37 | Very high | 0.00 |
| YT | | 0.87 | 2459.01 (1656-4256) | 0.52 | 2066.39 | Extremely high | 70.00 |
| KS | 2024 | 1.46 | 2102.85 (1450-3733) | 10.40 | 1767.10 | Extremely high | 100.00 |
| YC | | 0.79 | 453.02 (282-691) | 0.02 | 380.69 | Very high | 100.00 |
| JN | | 0.87 | 2459.03 (1656-4256) | 0.52 | 2066.41 | Extremely high | 100.00 |
| BD | | 1.13 | 795.34 (584-1124) | 1.16 | 668.35 | Very high | 70.00 |
| SZ | | 1.05 | 70.10 (26.56-119) | 0.42 | 58.91 | High | 80.00 |
| TH | | 1.03 | 99.53 (53-149) | 1.30 | 83.64 | High | 50.00 |
| PG | | 1.68 | 174.69 (126-226) | 3.85 | 146.80 | High | 90.00 |
| DQ | | 0.63 | 825.21 (483-1870) | 0.44 | 693.45 | Very high | 100.00 |
| HZ | | 0.91 | 459.85 (317-731) | 1.13 | 386.43 | Very high | 100.00 |
| WH | | 1.03 | 3989.04 (2687-7438) | 1.58 | 3352.13 | Extremely high | 100.00 |
| HK | | 0.90 | 556.90 (374-1011) | 0.53 | 467.98 | Very high | 100.00 |
| CS | | 1.45 | 3685.24 (2678-5950) | 3.36 | 3096.84 | Extremely high | 100.00 |
| SH | | 1.14 | 220.22 (158-298) | 0.65 | 185.06 | High | 100.00 |
| SY | | 1.40 | 1096.58 (851-1436) | 1.37 | 921.50 | Very high | 100.00 |
| KS | 2025 | 1.13 | 1853.61 (623-4822) | 15.77 | 1557.66 | Extremely high | 100.00 |
| YC | | 3.46 | 112.36 (52-190) | 12.32 | 94.42 | High | 100.00 |
| SZ | | 2.38 | 478.97 (229-917) | 8.36 | 402.50 | Very high | 85.00 |
| TJ | | 1.48 | 160.32 (72.98-265) | 1.61 | 134.72 | High | 70.00 |
| XX | | 1.54 | 4155.54 (3010-5727) | 0.80 | 3492.05 | Extremely high | 95.00 |
| DQ | | 1.81 | 236.05 (171-309) | 0.82 | 198.36 | High | 60.00 |
| HZ | | 2.23 | 1117.00 (823-1731) | 2.50 | 938.66 | Very high | 100.00 |
| WH | | 1.52 | 5700.90 (2988-14156) | 10.13 | 4790.67 | Extremely high | 100.00 |
| HK | | 1.63 | 79.13 (13-167) | 6.92 | 66.50 | High | 75.00 |
| CS | | 1.29 | 1199.48 (858-1565) | 3.31 | 1007.97 | Extremely high | 100.00 |

[a] Slope, calculated by Probit and Logit Analysis.

[b] FL, concentration of insecticide killing 50% of adults and its 95% fiducial limits.

[c] $\chi^2$, Chi-square testing linearity of dose-mortality responses.

[d] $RR_{50}$, resistance ratio = the $LC_{50}$ value of the field population divided by the $LC_{50}$ value of the sensitive reference strain.

[e] extremely high resistance ($RR_{50} \geq 1000$), very high resistance ($1000 < RR_{50} \leq 300$), high resistance ($300 < RR_{50} \leq 30$ )

[f] a total of 20 whitefly individuals for each population were used for TYLCV detection.

[g] the sensitive reference strain from the study by Wang et al., (83).

**Table S4** The laboratory strains of the vector *B. tabaci* MED used in the study

| Strain name | Origin strains [a] | Rearing year | Rearing plant | Resistance or sensitive | TYLCV-infected | Strain type |
|---|---|---|---|---|---|---|
| S1 | S#1 | From 2023 | Tomato | Sensitive | - | Non-viruliferous |
| S2 | S#2 | From 2023 | Tomato | Sensitive | - | Non-viruliferous |
| R1 | R#1 | From 2023 | Tomato | Resistance | - | Non-viruliferous |
| R2 | R#2 | From 2023 | Tomato | Resistance | - | Non-viruliferous |
| VS1 | S#1 | From 2023 | Tomato | Sensitive | + | Viruliferous |
| VS2 | S#2 | From 2023 | Tomato | Sensitive | + | Viruliferous |
| VR1 | R#1 | From 2023 | Tomato | Resistance | + | Viruliferous |
| VR2 | R#2 | From 2023 | Tomato | Resistance | + | Viruliferous |

[a] the origin of these strains from our previous work (33).

**Table S5** Details of technical grade insecticides used in the study

| Active agent | Abbr. | Formulation [a] | Group [b] | Manufacture |
|---|---|---|---|---|
| Imidacloprid | IMI | 95% TC | Neonicotinoids | Shanghai yuanye Bio-Tech. Co., Ltd, China |
| Thiamethoxam | THX | 95% TC | Neonicotinoids | Shanghai yuanye Bio-Tech. Co., Ltd, China |
| Clothianidin | CLO | 98% TC | Neonicotinoids | Shanghai yuanye Bio-Tech. Co., Ltd, China |
| Acetamiprid | ACE | 95.8% TC | Neonicotinoids | Shanghai yuanye Bio-Tech. Co., Ltd, China |

[a] TC indicates technical compounds.

[b] Group descriptors for all of the active ingredients listed in the table are the insecticide resistance action committee (IRAC) Mode of Action group names.

**Table S6** Developmental duration of the whitefly experimental strains

| Strains name | E (days) | N1 (days) | N2 (days) | N3 (days) | N4 (days) | N1-N4 (days) | Preadult (days) |
|---|---|---|---|---|---|---|---|
| S1 | 7.01±0.01 | 3.11±0.03 | 5.21±0.18 | 5.76±0.15 | 3.91±0.09 | 17.83±0.27 | 24.48±0.27 |
| (n) | (151) c | (152) a | (122) b | (100) b | (99) d | (100) d | (99) e |
| VS1 | 8.48±0.05 | 1.64±0.04 | 6.66±0.14 | 6.38±0.14 | 4.08±0.05 | 18.63±0.17 | 27.09±0.17 |
| (n) | (151) a | (152) d | (140) a | (125) a | (123) d | (124) c | (123) bc |
| S2 | 8.04±0.06 | 2.53±0.04 | 5.41±0.06 | 5.91±0.07 | 5.88±0.10 | 19.64±0.16 | 27.53±0.17 |
| (n) | (200) b | (188) c | (162) b | (136) b | (119) b | (120) b | (119) b |
| VS2 | 8.14±0.05 | 2.45±0.04 | 5.13±0.06 | 5.75±0.06 | 4.68±0.10 | 17.97±0.17 | 25.94±0.17 |
| (n) | (210) b | (201) c | (183) b | (153) b | (130) c | (131) cd | (130) d |
| R1 | 8.48±0.07 | 2.73± 0.05 | 4.62±0.05 | 5.61±0.06 | 6.39±0.11 | 19.49±0.15 | 27.31±0.14 |
| (n) | (196 ) a | (187) b | (163) c | (133) b | (105) ab | (106) b | (105) bc |
| VR1 | 8.00±0.05 | 2.56±0.04 | 4.40±0.06 | 6.25±0.07 | 6.04±0.09 | 19.06±0.13 | 26.94±0.14 |
| (n) | (220) b | (209) c | (188) c | (172) a | (134) b | (135) bc | (134) c |
| R2 | 8.34±0.05 | 2.90±0.05 | 4.74±0.05 | 5.94±0.07 | 6.65±0.10 | 19.80±0.14 | 27.94±0.10 |
| (n) | (205) ab | (188) b | (165) c | (126) b | (95) a | (96) b | (95) b |
| VR2 | 8.41±0.06 | 3.17±0.05 | 4.68±0.06 | 6.29±0.09 | 6.92±0.08 | 20.92±0.13 | 29.04±0.14 |
| (n) | (190) a | (173) a | (154) c | (133) a | (111) a | (112) a | (111) a |
| *F*-value | 60.79 | 119.10 | 66.06 | 10.39 | 166.20 | 37.10 | 52.98 |
| (*DF*, *N*) | (7, 1515) | (7,1439) | (7,1270) | (7, 1072) | (7, 908) | (7, 908) | (7, 908) |
| *P*-value | <0.0001 | <0.0001 | <0.0001 | <0.0001 | <0.0001 | <0.0001 | <0.0001 |

All biological raw data were analyzed using the TWO-SEX-MS program. n represents the number of individuals used in life-table study. Data are presented as mean ± SEM. Different letters in the same column indicate significant difference at $P < 0.05$ level determined by ANOVA through Tukey's HSD multiple comparisons test. The developmental stages examined` include egg (E), 1st-insatr nymph (N1), 2nd-instar nymph (N2), 3rd-instar nymph (N3) and 4th-instar nymph (N4). Preadult stage contains from E to N4, also refer to the total immature stage.

**Table S7** Reproductive parameters of the whitefly experimental strains

| Strain name | APOP (days) | TPOP (days) | OP (days) | Male adult longevity (days) | Female adult longevity (days) | Fecundity (No. of eggs laid/female) | Sex ratio (female%) |
|---|---|---|---|---|---|---|---|
| S1 | 0.32± 0.01 c | 24.46± 0.03 e | 22.98± 0.19 b | 20.69± 1.55 a | 27.60± 1.37 b | 110.74± 6.69 b | 57.57 |
| VS1 | 0.36± 0.06 bc | 27.11± 0.23 c | 27.74± 0.15 a | 21.33± 1.28 a | 32.90± 1.20 a | 156.10± 0.17 a | 58.54 |
| S2 | 0.89± 0.09 a | 28.05± 0.18 b | 22.26± 0.58 c | 18.67± 0.73 ab | 24.46± 0.67 bc | 130.02± 2.37 b | 62.18 |
| VS2 | 0.52± 0.09 b | 26.10± 0.24 d | 25.98± 0.63 a | 20.56± 0.54 a | 27.73 ± 0.71 b | 155.34± 2.74 a | 54.62 |
| R1 | 1.02± 0.09 a | 27.95± 0.16 b | 20.07± 0.56 d | 15.40± 0.50 b | 21.98± 0.63 c | 83.40± 2.05 c | 55.24 |
| VR1 | 1.09± 0.08 a | 27.75± 0.20 bc | 22.91± 0.58 c | 20.17± 0.37 a | 25.11± 0.64 bc | 151.59± 3.73 a | 55.97 |
| R2 | 1.18± 0.12 a | 28.98± 0.16 ab | 20.09± 0.65 d | 16.96± 0.50 b | 21.82± 0.73 c | 88.79± 2.40 c | 46.31 |
| VR2 | 0.96± 0.09 a | 29.50± 0.20 a | 22.31± 0.56 c | 17.69± 0.67 ab | 24.06± 0.64 bc | 146.40± 2.96 a | 64.86 |
| *F*-value | 14.75 | 50.28 | 16.20 | 6.57 | 17.57 | 50.17 | / |
| (*df*, *N*) | (7, 515) | (7, 515) | (7, 515) | (7, 385) | (7, 515) | (7, 515) | |
| *P*-value | <0.0001 | <0.0001 | <0.0001 | <0.0001 | <0.0001 | <0.0001 | / |

All biological raw data were analyzed using the TWO-SEX-MS program. Data are presented as mean ± SEM. Different letters in the same column indicate significant difference at $P < 0.05$ level determined by ANOVA through Tukey's multiple comparisons test. APOP, adult preoviposition period; TPOP, total preoviposition period; OP, oviposition period.

**Table S8** Life-table parameters and the relative fitness (*Rf*) associated with neonicotinoid resistance in the vector *B. tabaci* MED strains

| Strains | *r* | *λ* | $R_0$ | *T* | *GRR* | $Rf_r$ | $Rf_{R0}$ |
|---|---|---|---|---|---|---|---|
| S1 | 0.1069± 0.0038 | 1.1128± 0.0042 | 41.8013± 5.0332 | 34.9170± 0.4810 | 103.4500± 11.1020 | 1.0000 | 1.0000 |
| VS1 | 0.1149± 0.0028 | 1.1217± 0.0031 | 74.4283± 6.9189 | 37.5220± 0.3310 | 137.1900± 10.2570 | 1.0748 | 1.7804 |
| S2 | 0.1055± 0.0027 | 1.1112± 0.0030 | 48.1085± 4.5134 | 36.7190± 0.2120 | 97.7600± 6.4410 | 0.9869 | 1.1509 |
| VS2 | 0.1129± 0.0031 | 1.1195± 0.0034 | 52.5190± 5.1600 | 35.0840± 0.2500 | 101.6600± 7.1950 | 1.0561 | 1.2564 |
| R1 | 0.0902± 0.0033 | 1.0944± 0.0036 | 24.6786± 2.7715 | 35.5460± 0.2060 | 58.4900± 4.6840 | 0.8438 | 0.5904 |
| VR1 | 0.1070± 0.0028 | 1.1129± 0.0031 | 51.6773± 4.9930 | 36.8780± 0.2200 | 105.4400± 7.6720 | 1.0009 | 1.2363 |
| R2 | 0.0799± 0.0038 | 1.0831± 0.0042 | 19.0584± 2.6023 | 36.8990± 0.2270 | 58.3300± 6.5890 | 0.7474 | 0.4559 |
| VR2 | 0.1043± 0.0026 | 1.1099± 0.0029 | 55.4789± 5.2816 | 38.4980± 0.2530 | 126.4200± 9.5640 | 0.9757 | 1.3272 |

All biological raw data were analyzed using the TWO-SEX-MS program. Data are presented as mean ± SEM. *r,* the intrinsic rate of increase; $R_0$, the net reproductive rate; *λ*, the finite rate; *T*, the mean generation time; *GRR*, gross reproduction rate . $Rf_r$, the relative fitness (*Rf*) calculated with *r*. $Rf_{R0}$, the relative fitness (*Rf*) calculated with $R_0$.

**Table S9** Primers used in qPCR analysis

| Primer names | Primer sequence (5’-3’) |
|---|---|
| METTL3-F | TCTGCGGACACTTTAGGCATTAT |
| METTL3-R | AAATAACGATGTGACTGGCAATG |
| METTL14-F | TTAACGTCCTCTGCCTCTACCTC |
| METTL14-R | GGTTGTGTTTGTTCGTATCCAGC |
| WTAP-F | AGTTTACGCAAAGTTGAAAGCCCT |
| WTAP-R | TTCTTCAGTCACTTTTTCACCATCA |
| KIAA1429-F | AAGGAGGAGGACCATCAGGACA |
| KIAA1429-R | TCAGAGCGGTGGGAAAGAACA |
| RBM15-F | CCCGAGACGCCAAGCATAAT |
| RBM15-R | GGAGGTAAAGGTCGGTGGTGAT |
| ALKBH1-F | GATTTGAGGAGGTTCTGGATTTTTC |
| ALKBH1-R | TGTGAAAGGGTTGCGGATAAATA |
| ALKBH2-F | CCTGAACCTCCAGCCAAAAAGA |
| ALKBH2-R | CCATTTGAGAGGGGAGAGACTTTTA |
| ALKBH4-F | GTGATTCGGTTCTTACGCTTCG |
| ALKBH4-R | TCCCATTCATACCTGGCCTCA |
| ALKBH6-F | AAGAAGATATGTACCATCGCCACCT |
| ALKBH6-R | CTTTATTTTGGTACTCTTTGGTGCG |
| ALKBH7-F | ATGAGGAATGCGATGCGTTAT |
| ALKBH7-R | TTCTTCTTGATTTGTGGACTTGTTC |
| ALKBH8-F | AAGTCGTCCTGAACTGTCCTTATGG |
| ALKBH8-R | GAAGCATTCAACGAAAGAGACAAGA |
| Ex-F | GTGACAAGAAAGCGGGCATAA |
| Ex-R | TCAGCGAATCCCTTTACCAGAG |
| Va-F | ATGGACGACTGGGACTTTAGCG |
| Va-r | GCCGTCTTCTTCCTCGTATTGA |
| Bg-F | TCCGAGGGTACGTCAAAATGTC |
| Bg-R | GCGGTAACTCTTTGAGGTCCAG |
| Vg-F | TGGACTCCCGTATTATTGTGCC |
| Vg-R | TCAGAGTGCGTCCGTTGACTT |

F: forward primer; R: reverse primer.

**Table S10** Information of antibodies used in this study

| Antibody | Immune peptides / Product No. | Manufacture | Source |
|---|---|---|---|
| Ex | TTPGKPENGHAVKGVPPGDYRL (Thr5 to Leu26) | Jiaxuan Biotech., China | Rabbit |
| Va | RNRDDDDDNDGGDSYNRDRDDGGRG (Arg81 to Arg106) | Jiaxuan Biotech., China | Rabbit |
| Bg | DFRTAASTSRNTNYTKPTNYHGD (Asp1188 to Asp1210) | Jiaxuan Biotech., China | Rabbit |
| Vg | SKCEERSAYHFGITGLTNWKPAS (Ser218 to Ser240) | Jiaxuan Biotech., China | Rabbit |
| METTL3 | #ab240595 | Abcam, UK | Rabbit |
| METTL14 | #ab252562 | Abcam, UK | Rabbit |
| ALKBH4 | HGDNSKYNLFYSDHE (His197 to Glu211) | Jiaxuan Biotech., China | Rabbit |
| TYLCV-CP | KRRSWTYRPMYRKPR (Lys41 to Arg55) | AtaGenix, China | Mouse |
| TYLCV-C2 | GDKQSPLFQDNRTQP (Gly71 to Pro85) | AtaGenix, China | Mouse |
| TYLCV-C3 | NTINVTETHDIKYKFY (Cys119 to Tyr134) | AtaGenix, China | Mouse |
| *β*-actin | #ab8227 | Abcam, UK | Rabbit |
| HRP-IgG | #CW0103S | CWBIO, China | Rabbit |
| HRP-IgG | #ab205719 | Abcam, UK | Mouse |
| His | #ab9108 | Abcam, UK | Rabbit |
| GST | #ab9085 | Abcam, UK | Rabbit |
| $m^6A$ | #202003 | Synaptic Systems, Germany | Rabbit |

**Table S11** Primers used in RNAi experiments

| Primer names | Primer sequence (5'-3') |
|---|---|
| METTL3-F | TAATACGACTCACTATAGG GAACAGGACGGACTGGTCAT |
| METTL3-R | TAATACGACTCACTATAGG CACCATGCAATTTCCATCAG |
| METTL14-F | TAATACGACTCACTATAGG AACGGACAAAGGAACATTGC |
| METTL14-R | TAATACGACTCACTATAGG TTCGATACGTTCAGTGCAGC |
| WTAP-F | TAATACGACTCACTATAGG AAACCTCAGGAAGTGGGCTT |
| WTAP-R | TAATACGACTCACTATAGG CTGAGCTCACGTTCTGCTTG |
| KIAA1429-F | TAATACGACTCACTATAGG CATACCCATGCGTCAAAGTG |
| KIAA1429-R | TAATACGACTCACTATAGG GTAAGGCAAGGCACATTGGT |
| RBM15-F | TAATACGACTCACTATAGG ACCCGTACGGATCAAGAGTG |
| RBM15-R | TAATACGACTCACTATAGG ACCCGTACGGATCAAGAGTG |
| ALKBH1-F | TAATACGACTCACTATAGG GACTCAGCCGAGTGGTCATT |
| ALKBH1-R | TAATACGACTCACTATAGG GAAGTTCGATTTTCGTCCCA |
| ALKBH2-F | TAATACGACTCACTATAGG TTTGTGTTTGGAAAATGGCA |
| ALKBH2-R | TAATACGACTCACTATAGG GTGCTTCCATGGGTGAAGAT |
| ALKBH4-F | TAATACGACTCACTATAGG TAAAGGCGTCAGGACTTGCT |
| ALKBH4-R | TAATACGACTCACTATAGG GCCGGGTCATATTCAAGAGA |
| ALKBH6-F | TAATACGACTCACTATAGG ACCGAGGTTCCTCCAAAAGT |
| ALKBH6-R | TAATACGACTCACTATAGG AGATCCGCAGGTTATGGTCG |
| ALKBH7-F | TAATACGACTCACTATAGG ATGGTGAAACGCCTCACTTC |
| ALKBH7-R | TAATACGACTCACTATAGG TATTGCCGCAAAATCGTACA |
| ALKBH8-F | TAATACGACTCACTATAGG TATCCATGTCACCAGGCAAA |
| ALKBH8-R | TAATACGACTCACTATAGG GCGGGATACTTTCCATCAGA |
| Ex-F | TAATACGACTCACTATAGG GCAAGAACGCTAGTCAATTACGC |
| Ex-R | TAATACGACTCACTATAGG CCATTTGTTTCGGCTACAGGCT |
| Va-F | TAATACGACTCACTATAGG GGCGAAGTGGACCCCAATAA |
| Va-r | TAATACGACTCACTATAGG TGGATGACTGGCAGAAGAAAGG |
| Bg-F | TAATACGACTCACTATAGG CAAACCTACTCCGCCGAACA |
| Bg-R | TAATACGACTCACTATAGG GTGGCTCCAAGGCATAAGATAAGA |
| Vg-F | TAATACGACTCACTATAGG GCGAATGCGAGACTCTTTACG |
| Vg-R | TAATACGACTCACTATAGG CTGATGACGATTTTGTTGGTGG |

F: forward primer; R: reverse primer.

**Table S12** Primers used in CDS cloning and RACE cloning

| Primer names | Primer sequence (5'-3') |
|---|---|
| METTL3-F | ATGTCAGATGCATGGGAGGATATC |
| METTL3-R | TCATGATTTGGGAGGCACCAT |
| METTL14-F | ATGAGTATGTTGCGAGAGTTGAAAG |
| METTL14-R | TTAACGTCCTCTGCCTCTACCTC |
| WTAP-F | GCGTACGTTTTCGTTTTTTAAGGTT |
| WTAP-R | GATCAATCCCTGGTTGCGGTT |
| KIAA1429-F | GTGTTTACAGTGATTTTCTTTGGCC |
| KIAA1429-R | ACCTGGGGAATGGACGTAAGAG |
| RBM15-F | ATGAATGAAGTGTCGGGCAAGT |
| RBM15-R | TGAGATGATTCAAGCCGTGCC |
| ALKBH1-F | ATGACTTTTCGGGATAAATTCAAAT |
| ALKBH1-R | TCAAGCCTCCTTTAAGAGAGTTTCC |
| ALKBH2-F | TGTCATTTCGCTCCTGTCTTTTC |
| ALKBH2-R | AGTCCTGTGACATAAAAGAGTTTCCAT |
| ALKBH4-F | ATGGACCGATCCAACTTTTGTG |
| ALKBH4-R | TCAAGTGAGGAAAAAGTTCTTTGCT |
| ALKBH6-F | GCTCATCCTAGGATTGTCACATCTC |
| ALKBH6-R | ATGCCAATTGGTCATTTTATCACAT |
| ALKBH7-F | GCCATATGATGCTTTTTAGTCACTG |
| ALKBH7-R | GACTGAAGACTCAGAGGTTAGACGG |
| ALKBH8-F | ATGGAAGATCTACCTCTTTTGTATCTTG |
| ALKBH8-R | GCTCTTCACTAAGATGACACACCAA |
| Vg-CDS1-F | ATCATCATGTGGACTCCAGCG |
| Vg-CDS1-R | GTAAAGCCGATGTCAACGCC |
| Vg-CDS2-F | TCGCCTGGAAATCCTTCTGT |
| Vg-CDS2-R | TGCTTACGCTCGTAGCCCA |
| Vg-CDS3-F | CGTAACGATTTCTTCTCCGCT |
| Vg-CDS3-F | TACTGTGTTGCCTAAGCAAACATAC |
| Vg-5'UTR-RACE-R1 | AATGACACGGTTGATGTTGGAGCGG |
| Vg-5'UTR-RACE-R2 | TGGCGGGTTTCCAGTTGGTGAGAC |
| Vg-5'UTR-RACE-R3 | TTTCCTTGGGGAGTTGGTTGTGGCT |
| Vg-5'UTR-RACE-R4 | CCTTGATGATGTTGAGTTCCCAGTCGG |
| Vg-5'UTR-RACE-R5 | GCGGCTGCCACGAGCAAACACAA |
| Vg-3'UTR-RACE-F1 | TGACTTTTTCGCCCCCCACCCCG |
| Vg-3'UTR-RACE-F2 | CCTGATGGTGCTATCCGCTTCTTCGC |
| Vg-3'UTR-RACE-F3 | ACGCTGACGACTTCACCGCCCCT |
| Vg-3'UTR-RACE-F4 | TCTTCCCCCTCTCAAGTGAAATCCCAG |

F: forward primer; R: reverse primer.

**Table S13** UPLC-MS/MS parameters used in this study

| Compound | Transitions (m/z) | CV (V)[a] | CE (V)[b] | DT (ms)[c] | Quantitative ion |
|---|---|---|---|---|---|
| A | 268.1+>119.1 | 40 | 40 | 0.082 | Positive |
| | 268.1+>136.0 | 40 | 16 | | |
| $m^6A$ | 282.1+>123.1 | 38 | 38 | 0.082 | Positive |
| | 282.1+>150.1 | 38 | 14 | | |
| dA | 252.1+>119.1 | 14 | 36 | 0.081 | Positive |
| | 252.1+>136.0 | 14 | 10 | | |

[a] Cone voltage (V)

[b] Collione energy (V)

[c] Dwell time (ms)

**Table S14** Predicted m6A sites of *Vg* gene by SRAMP

| Position | Name | RNA sequence (5'-3', m6A motif) | Score (combined) | Confidence | Exon |
|---|---|---|---|---|---|
| 5'-UTR | A516 | GUGAA**UGACU**GCAGC | 0.596 | Moderate | - |
| 5'-UTR | A610 | AGCUC**AGACU**UCAAG | 0.565 | Moderate | - |
| 5'-UTR | A773 | AGCUC**GGACU**GGGGA | 0.611 | High | - |
| CDS | A7 | CAUGU**GGACU**CCAGC | 0.618 | High | E1 |
| CDS | A188 | UCCAG**UGACA**GAUUA | 0.554 | Low | E3 |
| CDS | A240 | ACACU**GAACU**CCCUG | 0.602 | High | E4 |
| CDS | A284 | CAGUU**GAACU**ACAAG | 0.644 | High | E4 |
| CDS | A293 | UACAA**GAACA**UGCCC | 0.589 | Moderate | E4 |
| CDS | A384 | ACUGG**GAACU**CAACA | 0.611 | High | E4 |
| CDS | A440 | GGUCA**AAACU**UGAAG | 0.57 | Moderate | E4 |
| CDS | A512 | AUGGA**AGACU**CCGUC | 0.555 | Low | E4 |
| CDS | A621 | AAAAC**GGACA**AGUCA | 0.636 | High | E4 |
| CDS | A735 | AGAUG**GGACA**GUUCC | 0.652 | High | E4 |
| CDS | A884 | GUCAU**GAACU**UGACU | 0.676 | Very high | E5 |
| CDS | A889 | GAACU**UGACU**UUGGC | 0.586 | Moderate | E5 |
| CDS | A1467 | GCAAC**GGACA**CAACG | 0.626 | High | E5 |
| CDS | A1476 | ACAAC**GGACA**CAACG | 0.621 | High | E5 |
| CDS | A1494 | ACAAC**GGACA**CAACG | 0.639 | High | E5 |
| CDS | A1512 | ACAAC**GGACA**CAACGG | 0.641 | High | E5 |
| CDS | A1521 | ACAAC**GGACA**CAAUGG | 0.637 | High | E5 |
| CDS | A1530 | ACAAU**GGACA**CAACG | 0.659 | High | E5 |
| CDS | A1548 | ACAAU**GGACA**AAACA | 0.658 | High | E5 |
| CDS | A1850 | GAAGA**GGACU**AUGAA | 0.669 | High | E5 |
| CDS | A1916 | AUCGG**AAACU**ACGGU | 0.549 | Low | E5 |
| CDS | A2064 | UUGAG**GAACU**CAGAA | 0.625 | High | E5 |
| CDS | A2094 | AAAUG**GAACA**AGCUU | 0.550 | Low | E5 |
| CDS | A2214 | GUACC**GGACC**UGCCU | 0.573 | Moderate | E6 |
| CDS | A2333 | UACAU**GAACU**CUUUC | 0.676 | Very high | E6 |
| CDS | A2358 | ACUAU**GAACA**AAACC | 0.548 | Low | E7 |
| CDS | A2609 | AUUGG**AAACU**UUGCU | 0.551 | Low | E7 |
| CDS | A2712 | UUAAC**GAACU**UGCAC | 0.629 | High | E7 |
| CDS | A2952 | GUCAA**GAACU**CGCCC | 0.646 | High | E7 |
| CDS | A2998 | CCCUA**AGACU**UACGG | 0.601 | High | E8 |
| CDS | A3098 | AGUGA**GGACU**CCUUA | 0.631 | High | E8 |
| CDS | A3200 | GCCAG**UGACC**UCGUU | 0.539 | Low | E9 |
| CDS | A3222 | UCCAA**GAACA**GUUCA | 0.627 | High | E9 |
| CDS | A3266 | GAAAU**GAACC**AGCAA | 0.547 | Low | E9 |
| CDS | A3297 | UCUUC**AGACA**GUUCA | 0.545 | Low | E9 |
| CDS | A3317 | AAGCC**UGACU**ACCCA | 0.543 | Low | E9 |
| CDS | A3450 | AAAAU**GGACA**AUCUU | 0.632 | High | E10 |
| CDS | A3702 | AAUCU**GAACU**UGGAU | 0.623 | High | E11 |

| | | | | | |
|---|---|---|---|---|---|
| CDS | A3758 | GCCAA**GAACA**UCUUU | 0.575 | Moderate | E11 |
| CDS | A3800 | AAUGU**GGACA**UCGCC | 0.674 | Very high | E11 |
| CDS | A3842 | CCUAU**GAACA**ACCAG | 0.553 | Low | E11 |
| CDS | A3791 | AACAA**GAACA**AGUUC | 0.553 | Low | E11 |
| CDS | A4061 | UUCGC**UGACU**UCUAC | 0.533 | Low | E12 |
| CDS | A4202 | GCCUU**GAACU**AUGCC | 0.657 | High | E12 |
| CDS | A4247 | GAAGG**UAACU**CUAAC | 0.540 | Low | E13 |
| CDS | A4311 | ACAAC**GAACA**GAGAA | 0.542 | Low | E13 |
| CDS | A4401 | UCUUC**GGACA**AAAGA | 0.567 | Moderate | E14 |
| CDS | A4691 | UUCCG**UGACU**ACUUC | 0.56 | Moderate | E16 |
| CDS | A4850 | GUCAA**GAACU**ACACU | 0.594 | Moderate | E17 |
| CDS | A4910 | UCUGA**AAACU**UCGGA | 0.533 | Low | E17 |
| CDS | A5054 | GUCGC**UGACU**UUUUC | 0.531 | Low | E17 |
| CDS | A5306 | UACAG**AGACU**ACAAC | 0.625 | High | E18 |
| CDS | A5462 | GGAAA**GAACA**ACGCU | 0.569 | Moderate | E19 |
| CDS | A5499 | AAACA**AGAAC**AACAAU | 0.553 | Low | E19 |
| CDS | A5528 | UUCAC**UGACU**UCCAC | 0.608 | High | E19 |
| CDS | A5538 | UCCAC**GGACA**CAAUG | 0.629 | High | E19 |
| CDS | A5613 | AAUCU**GGACU**UCAAG | 0.682 | Very high | E19 |
| CDS | A5738 | CCUCA**AAACU**GUGUC | 0.629 | High | E20 |
| CDS | A5762 | CCAGA**AGACU**UCGCU | 0.612 | High | E20 |
| CDS | A5789 | GUCAU**UGACU**CUUCU | 0.648 | High | E20 |
| CDS | A6002 | UACAA**GAACA**ACAAG | 0.544 | Low | E20 |
| CDS | A6020 | GCCAA**GGACC**AGGAC | 0.581 | Moderate | E20 |
| CDS | A6026 | GACCA**GGACU**CCUCC | 0.648 | High | E20 |
| CDS | A6086 | UCCAG**UGACU**CCAGC | 0.559 | Moderate | E20 |
| CDS | A6360 | AAAAC**GGACC**CUCUA | 0.575 | Moderate | E20 |
| CDS | A6375 | UCCGC**AAACU**GUACC | 0.588 | Moderate | E20 |
| CDS | A6482 | AUGGU**UGACU**UCUAC | 0.578 | Moderate | E21 |
| CDS | A6599 | GAAGU**GAACA**UUCCC | 0.531 | Low | E21 |
| 3'UTR | A14 | AAUCU**GGACU**UCAGC | 0.668 | High | - |
| 3'UTR | A138 | CCUCA**AAACU**GUGUC | 0.631 | High | - |
| 3'UTR | A162 | CCAGA**AGACU**UCGCU | 0.618 | High | - |
| 3'UTR | A189 | GUCAU**UGACU**CUUCU | 0.657 | High | - |
| 3'UTR | A240 | CAGCA**AAACU**UCUGU | 0.533 | Low | - |
| 3'UTR | A402 | UACAA**GAACA**ACAAG | 0.54 | Low | - |
| 3'UTR | A420 | GCCAA**GGACC**AGGAC | 0.575 | Moderate | - |
| 3'UTR | A426 | GACCA**GGACU**CCUCC | 0.644 | High | - |
| 3'UTR | A486 | UCCAG**UGACU**CCAGC | 0.569 | Moderate | - |
| 3'UTR | A709 | AAAAC**GGACC**CUCUA | 0.572 | Moderate | - |
| 3'UTR | A724 | UCCGC**AAACU**GUACC | 0.585 | Moderate | - |
| 3'UTR | A831 | AUGGU**UGACU**UCUAC | 0.588 | Moderate | - |

**Table S15** Primers used in MeRIP-qPCR analysis for identification of *Vg* m$^6$A sites

| Position | Putative m$^6$A sites | F/R | Primer sequence (5'-3') |
|---|---|---|---|
| 5'UTR | A516 | F | ATCTGAGAGCAACGCGAAAA |
| 5'UTR | A516 | R | ATTTCGCTGCAGTCATTCAC |
| 5'UTR | A610 | F | ACCTCCCTCAAAAGCTCAGA |
| 5'UTR | A610 | R | CGATTCATTTTCCGCGCGAA |
| 5'UTR | A773 | F | CTTTTGGTGGAATGCTCGGA |
| 5'UTR | A773 | R | CTCCCCAGTCCGAGCTTTTA |
| CDS | A621 | F | CCCTCCAAACTAGACCCTGG |
| CDS | A621 | R | CAAAGTGGTAGGCAGAACGT |
| CDS | A735 | F | ACGTTCTGCCTACCACTTTG |
| CDS | A735 | R | GACACGGTTGATGTTGGAGC |
| CDS | A884 | F | TTGACTTTGGCCTCCTTCCA |
| CDS | A884 | R | GTTGTAGTGTTGAGCGTCGG |
| CDS-A1467s | A1467、A1476 | F | GAGAACGTCTCTCGCAGTCG |
| CDS-A1467s | A1467、A1476 | R | TGTTTCCGTTGTGTCCGTTG |
| CDS-A1494s | A1494、A1512、A1530 | F | CAACGGAAACAACGGACACA |
| CDS-A1494s | A1494、A1512、A1530 | R | GCTGGCGAATTTACCGTTGT |
| CDS | A2998 | F | CAAATCCGCCCTTGAAAACG |
| CDS | A2998 | R | AGACACTCTGTAGGCAAGGT |
| CDS | A3098 | F | AAGAAATGGCTCCGCGACTA |
| CDS | A3098 | R | TGGAGGTCATGGCTCTGAAG |
| CDS | A3222 | F | TTCAGAGCCATGACCTCCAG |
| CDS | A3222 | R | GGGCTTCAACTTCTTGTGGG |
| CDS | A3800 | F | TTCAACCACGAACGTTACGT |
| CDS | A3800 | R | AATTCGACCTTGGTGTTGGC |
| CDS | A5538 | F | AACTCCGGAAAGAACAACGC |
| CDS | A5538 | R | TGTCCGTGGAAGTCAGTGAA |
| CDS | A5613 | F | TTCCTGATGGTGCTATCCGC |
| CDS | A5613 | R | GTTGGAGTAGGTTCCGCACA |
| CDS-A5738s | A5738、A5762、A5789 | F | CGCATCAAGATTCAAGCTGC |
| CDS-A5738s | A5738、A5762、A5789 | R | AGCGAAGTCTTCTGGGTTCT |
| CDS | A6026 | F | CCCTCCAACTCCCAGTACAA |
| CDS | A6026 | R | GGAGGAGGAAGAAGAGGAGC |
| CDS-6360s | A6360、A6375 | F | CAGCCAAGAAGCCAGTTACG |
| CDS-6360s | A6360、A6375 | R | TGAAGCACATGTCGTCTCCT |
| 3'UTR | A14 | F | CCCAATCTGGACTTCAGCCA |
| 3'UTR | A14 | R | GTTGGAGTAGGTTCCGCACA |
| 3'UTR | A138 | F | AGATTCAAGCTGCCAACCAA |
| 3'UTR | A138 | R | AGCGAAGTCTTCTGGGTTCT |

| | | | |
|---|---|---|---|
| 3'UTR | A189 | F | GTGTCTACAAGAACCCAGAAGA |
| 3'UTR | A189 | R | CTTGCGGGCACAGAAGTTTT |
| 3'UTR | A402 | F | CCCTCCAACTCCCAGTACAA |
| 3'UTR | A402 | R | GGGAGGAAGAAGAGGAGCTG |
| 3'UTR | A426 | F | ACAAGTACGCCAAGGACCAG |
| 3'UTR | A426 | R | TGGGGGAGGAAGAAGAGGAG |
| 3'UTR | A486 | F | GCAGCTCATCCAGTGACTCC |
| 3'UTR | A486 | R | GGACTGGGAAGAGCTGGAAG |
| 3'UTR | A724 | F | CAGCCAAGAAGCCAGTTACG |
| 3'UTR | A724 | R | TGAAGCACATGTCGTCTCCT |

F: forward primer; R: reverse primer.

**Table S16** Molecular docking and molecular dynamics simulations of METTL14-C2 interaction and binding free energies

| METTL14 binding residues | C2 binding residues | Bonds type | Distance ( Å) |
|---|---|---|---|
| R14 | D111 | Salt bridge | 3.4 |
| R14 | H105 | π–π | 3.8 |
| R385 | I101 | Hydrogen | 2.6 |
| G379 | S127 | Hydrogen | 2.8 |
| G379 | D128 | Salt bridge | 3.4 |
| R218 | D122 | Salt bridge | 3.6 |
| D123 | D122 | Hydrogen | 2.7 |
| R209 | D122 | Salt bridge | 3.9 |
| R209 | H95 | Salt bridge | 4.4 |
| K249 | D99 | Salt bridge | 3.4 |
| R214 | S40 | Hydrogen | 2.8 |
| D184 | R30 | Salt bridge | 3.4 |

| Binding free energy (MM/GBSA analysis) | kcal/mol |
|---|---|
| $\triangle E_{vdw}$ | -132.84 ± 9.69 |
| $\triangle E_{ele}$ | -37.09 ± 5.37 |
| $\triangle E_{GB}$ | 42.53 ± 4.39 |
| $\triangle E_{surf}$ | -21.75 ± 0.65 |
| $\triangle E_{Gas}$ | -169.93 ± 15.80 |
| $\triangle E_{solv}$ | 20.78 ± 14.40 |
| $\triangle E_{Bind}$ | -149.15 ± 8.28 |

Note: Van der Waals energy ($\triangle E_{vdw}$), Electrostatic energy ($\triangle E_{ele}$), Polar solvation energy ($\triangle E_{GB}$), Nonpolar solvation energy ($\triangle E_{surf}$), Gas-phase free energy ($\triangle E_{Gas}$), Solvation free energy ($\triangle E_{solv}$), Total binding free energy (TOTAL, $\triangle E_{Bind}$).

**Table S17** Molecular docking and molecular dynamics simulations of ALKBH4-CP interaction and binding free energies

| ALKBH4 binding residues | CP binding residues | Bonds type | Distance (Å) |
|---|---|---|---|
| E271 | R4 | Salt bridge | 3.8 |
| E274 | R67 | Salt bridge | 3.4 |
| H117 | Y57 | Hydrogen | 2.6 |
| D147 | R144 | Salt bridge | 3.6 |
| E178 | R142 | Salt bridge | 3.4 |
| R179 | D176 | Hydrogen | 2.7 |
| E243 | R182 | Salt bridge | 3.6 |
| E71 | T169 | Hydrogen | 2.8 |
| N67 | S166 | Hydrogen | 2.8 |
| H60 | F161 | π–π | 5.8 |

| **Binding free energy (MM/GBSA analysis)** | **kcal/mol** |
|---|---|
| $\triangle E_{vdw}$ | -130.26 ± 10.70 |
| $\triangle E_{ele}$ | -26.37 ± 7.31 |
| $\triangle E_{GB}$ | 34.44 ± 6.03 |
| $\triangle E_{surf}$ | -21.23 ± 1.09 |
| $\triangle E_{Gas}$ | -156.63 ± 17.72 |
| $\triangle E_{solv}$ | 13.21 ± 15.90 |
| $\triangle E_{Bind}$ | -143.42 ± 9.88 |

Note: Van der Waals energy ($\triangle E_{vdw}$), Electrostatic energy ($\triangle E_{ele}$), Polar solvation energy ($\triangle E_{GB}$), Nonpolar solvation energy ($\triangle E_{surf}$), Gas-phase free energy ($\triangle E_{Gas}$), Solvation free energy ($\triangle E_{solv}$), Total binding free energy (TOTAL, $\triangle E_{Bind}$).

**Table S18** Molecular docking of METTL14-ssRNA interaction with C2 protein or without

| METTL14 binding residues | 3'UTR-A14 binding sites | C2 binding residues | Bond type | Distance (Å) |
|---|---|---|---|---|
| R209 | G19 | without C2 | Salt bridge | 4.0 |
| R209 | G19 | without C2 | Hydrogen | 2.7 |
| R391 | C20 | without C2 | π–π | 5.4 |
| K368 | C24 | without C2 | Salt bridge | 3.5 |
| K366 | U25 | without C2 | Hydrogen | 2.7 |
| K366 | G12 | without C2 | Hydrogen | 2.8 |
| K366 | M3 | without C2 | Salt bridge | 3.7 |
| K366 | G11 | without C2 | Hydrogen | 2.8 |
| R5 | C3 | without C2 | Salt bridge | 3.9 |
| R376 | C17 | without C2 | π–π | 5.9 |
| R385 | U8 | without C2 | Hydrogen | 2.8 |
| R380 | A7 | without C2 | Hydrogen | 2.7 |
| Q12 | C5 | without C2 | Salt bridge | 3.4 |
| K16 | U8 | without C2 | Salt bridge | 3.9 |
| - | U15 | S9 (with) | Hydrogen | 2.6 |
| - | A6 | K24 (with) | Salt bridge | 3.8 |
| R213 | A18 | - (with) | Hydrogen | 2.8 |
| R376 | C17 | - (with) | Hydrogen | 2.7 |
| R218 | G19 | - (with) | π–π | 3.8 |
| K366 | G11 | - (with) | Salt bridge | 3.6 |
| R385 | C9 | - (with) | Hydrogen | 2.8 |
| N382 | U8 | - (with) | Hydrogen | 2.7 |
| R380 | A13 | - (with) | π–π | 3.7 |
| R5 | C5 | - (with) | Hydrogen | 2.7 |
| K8 | C2 | - (with) | Salt bridge | 3.8 |
| Q20 | C3 | - (with) | Hydrogen | 2.8 |
| **METTL14 binding residues** | **3'UTR-A138 binding sites** | **C2 binding residues** | **Bond type** | **Distance (Å)** |
| R219 | U19 | without C2 | Hydrogen | 2.6 |
| R391 | A22 | without C2 | Hydrogen | 2.7 |
| R389 | C23 | without C2 | Hydrogen | 2.7 |
| R94 | A13 | without C2 | Hydrogen | 2.7 |
| K13 | A25 | without C2 | Salt bridge | 4.0 |
| T60 | A12 | without C2 | Hydrogen | 2.7 |
| R94 | A12 | without C2 | Salt bridge | 3.5 |
| R55 | C7 | without C2 | Hydrogen | 2.6 |
| K56 | C4 | without C2 | Salt bridge | 4.0 |
| K8 | U17 | without C2 | Salt bridge | 3.9 |
| F245 | U15 | without C2 | π–π | 4.0 |
| F245 | G16 | without C2 | π–π | 5.4 |
| - | A11 | S130 | Hydrogen | 2.6 |

| | | | | |
|---|---|---|---|---|
| K257 | A22 | - (with) | Salt bridge | 3.6 |
| R218 | U19 | - (with) | Hydrogen | 2.6 |
| R261 | A24 | - (with) | π–π | 4.0 |
| K368 | C14 | - (with) | Salt bridge | 3.5 |
| K10 | C4 | - (with) | Salt bridge | 3.6 |
| K13 | C4 | - (with) | Salt bridge | 3.9 |
| K13 | C5 | - (with) | Salt bridge | 3.3 |
| **METTL14 binding residues** | **CDS-A621 binding sites** | **C2 binding residues** | **Bond type** | **Distance (Å)** |
| Agr391 | C2 | without C2 | Salt bridge | 3.9 |
| K10 | A8 | without C2 | Salt bridge | 3.9 |
| K10 | G3 | without C2 | Hydrogen | 2.7 |
| R380 | A13 | without C2 | Hydrogen | 2.7 |
| R380 | A15 | without C2 | π–π | 4.5 |
| R219 | A16 | without C2 | π–π | 4.0 |
| S373 | A16 | without C2 | Hydrogen | 2.7 |
| T259 | C22 | without C2 | Hydrogen | 2.7 |
| - | C25 | K17 | Salt bridge | 3.8 |
| - | U21 | K21 | Salt bridge | 3.7 |
| - | A8 | Y42 | π–π | 4.8 |
| K150 | G12 | - (with) | Salt bridge | 3.5 |
| R306 | G17 | - (with) | π–π | 3.4 |
| **METTL14 binding residues** | **CDS-A884 binding sites** | **C2 binding residues** | **Bond type** | **Distance (Å)** |
| R5 | U21 | without C2 | Salt bridge | 3.9 |
| R14 | U7 | without C2 | Hydrogen | 2.8 |
| K15 | G6 | without C2 | Salt bridge | 3.6 |
| R376 | G6 | without C2 | Hydrogen | 2.7 |
| R376 | G4 | without C2 | Hydrogen | 2.8 |
| R376 | A3 | without C2 | Salt bridge | 3.8 |
| R41 | A3 | without C2 | Hydrogen | 2.8 |
| R391 | U15 | without C2 | Hydrogen | 2.8 |
| R387 | A13 | without C2 | Hydrogen | 2.7 |
| R218 | C14 | without C2 | Hydrogen | 2.7 |
| S378 | U16 | without C2 | Hydrogen | 2.8 |
| R261 | G11 | without C2 | Hydrogen | 2.8 |
| R262 | G11 | without C2 | Salt bridge | 3.9 |
| - | U15 | R32 | π–π | 3.6 |
| - | A18 | K26 | Hydrogen | 2.7 |
| - | A18 | K26 | Salt bridge | 3.3 |
| - | G17 | K25 | Salt bridge | 4.0 |
| K15 | U5 | - (with) | Salt bridge | 4.0 |
| K15 | U5 | - (with) | Hydrogen | 2.7 |
| K10 | U5 | - (with) | Hydrogen | 2.8 |

| | | | | |
|---|---|---|---|---|
| K10 | U7 | - (with) | Salt bridge | 3.8 |
| K368 | C8 | - (with) | Salt bridge | 3.7 |
| G367 | G11 | - (with) | Hydrogen | 2.8 |
| S263 | G11 | - (with) | Hydrogen | 2.7 |
| R218 | C14 | - (with) | Hydrogen | 2.8 |
| R262 | A12 | - (with) | Hydrogen | 2.8 |
| N272 | A13 | - (with) | Hydrogen | 2.6 |

**Table S19** Molecular docking of ALKBH4-ssRNA interaction with CP protein or without

| **ALKBH4 binding residues** | **3'UTR-A14 binding sites** | **CP binding residues** | **Bond type** | **Distance (Å)** |
|---|---|---|---|---|
| R104 | C17 | without CP | Hydrogen | 2.7 |
| K106 | U16 | without CP | Salt bridge | 3.6 |
| K17 | C1 | without CP | Hydrogen | 2.7 |
| F115 | A18 | without CP | π–π | 5.3 |
| K212 | A6 | without CP | Hydrogen | 2.8 |
| - | C1 | R142 | π–π | 3.5 |
| - | C4 | R182 | Hydrogen | 2.7 |
| - | A6 | R20 | Hydrogen | 2.6 |
| - | A18 | Y51 | Hydrogen | 2.7 |
| - | U8 | Y51 | π–π | 5.0 |
| - | G19 | R43 | π–π | 3.6 |
| - | U16 | Q37 | Hydrogen | 2.7 |
| K106 | C24 | - (with) | Salt bridge | 3.6 |
| K119 | C21 | - (with) | Salt bridge | 2.7 |
| R118 | U8 | - (with) | π–π | 4.2 |
| K116 | C9 | - (with) | Hydrogen | 2.6 |
| K116 | U8 | - (with) | Hydrogen | 2.7 |
| **ALKBH4 binding residues** | **3'UTR-A138 binding sites** | **CP binding residues** | **Bond type** | **Distance (Å)** |
| F275 | G3 | without CP | π–π | 5.3 |
| F115 | C14 | without CP | Hydrogen | 2.8 |
| K106 | U15 | without CP | Salt bridge | 3.7 |
| K106 | C14 | without CP | Salt bridge | 3.5 |
| T14 | A12 | without CP | Hydrogen | 2.7 |
| D2 | U19 | without CP | Hydrogen | 2.7 |
| K17 | U19 | without CP | π–π | 5.0 |
| R162 | A22 | without CP | Hydrogen | 2.7 |
| - | U8 | K206 | Salt bridge | 3.8 |
| - | C7 | K113 | Salt bridge | 3.8 |
| - | U8 | K113 | Hydrogen | 2.7 |
| - | A11 | Y116 | Hydrogen | 2.5 |
| - | A12 | R249 | Hydrogen | 2.7 |
| - | U15 | K247 | Hydrogen | 2.7 |
| - | U15 | S77 | Hydrogen | 2.7 |
| - | G16 | K220 | Salt bridge | 3.9 |
| - | U17 | K220 | Salt bridge | 3.8 |
| - | U21 | K85 | Salt bridge | 3.9 |
| - | A24 | G130 | Hydrogen | 2.8 |
| Lys202 | C7 | - (with) | Salt bridge | 3.7 |
| **ALKBH4 binding residues** | **CDS-A621 binding sites** | **CP binding residues** | **Bond type** | **Distance (Å)** |

| | | | | |
|---|---|---|---|---|
| K10 | U21 | without CP | Salt bridge | 3.4 |
| R104 | G17 | without CP | Hydrogen | 2.8 |
| K106 | U18 | without CP | Salt bridge | 3.8 |
| K116 | G17 | without CP | Hydrogen | 2.7 |
| K116 | C19 | without CP | Hydrogen | 2.7 |
| D172 | G17 | without CP | Hydrogen | 2.7 |
| - | G11 | K202 | Hydrogen | 2.8 |
| - | A20 | N131 | Hydrogen | 2.9 |
| - | C22 | A238 | Hydrogen | 2.9 |
| - | G23 | D162 | Hydrogen | 2.9 |
| K202 | G3 | - (with) | Salt bridge | 3.4 |
| K213 | C1 | - (with) | Hydrogen | 2.9 |
| D199 | G12 | - (with) | Hydrogen | 2.9 |
| K195 | C14 | - (with) | Salt bridge | 3.7 |
| K66 | G17 | - (with) | Hydrogen | 3.0 |

| **ALKBH4 binding residues** | **CDS-A884 binding sites** | **CP binding residues** | **Bond type** | **Distance (Å)** |
|---|---|---|---|---|
| K121 | U5 | without CP | Salt bridge | 3.7 |
| K119 | G6 | without CP | Hydrogen | 2.7 |
| K119 | U7 | without CP | Salt bridge | 3.8 |
| K112 | A9 | without CP | Hydrogen | 2.7 |
| K110 | A9 | without CP | Salt bridge | 3.6 |
| K116 | U16 | without CP | Salt bridge | 4.0 |
| R3 | C14 | without CP | Hydrogen | 2.8 |
| R3 | U15 | without CP | Hydrogen | 2.7 |
| K106 | G11 | without CP | Salt bridge | 3.4 |
| K106 | G11 | without CP | Hydrogen | 2.8 |
| Q101 | A12 | without CP | Hydrogen | 2.7 |
| R104 | A12 | without CP | Hydrogen | 2.8 |
| - | C15 | T169 | Hydrogen | 2.6 |
| - | G17 | K206 | Salt bridge | 3.4 |
| - | U15 | R4 | Hydrogen | 2.7 |
| - | U15 | R20 | Hydrogen | 2.7 |
| - | A13 | R55 | Hydrogen | 2.7 |
| - | A13 | R58 | Hydrogen | 2.7 |
| - | A12 | K16 | π–π | 4.3 |

**Table S20** The mutant recombinant proteins used in this study

| Vector | Name | Mutation | Usage |
|---|---|---|---|
| pET28a | METTL14(MU) | Mutating enzyme domain | UPLS-MS、SELECT、enzyme activity |
| pET28a | ALKBH4(MU) | Mutating enzyme domain | UPLS-MS、SELECT、enzyme activity |
| pGBKT7 | METTL14$^{MU}$ | Mutating binding residues | Y2H: METTL14-C2 interaction |
| pGBKT7 | ALKBH4$^{MU}$ | Mutating binding residues | Y2H: ALKBH4-CP interaction |
| pGADT7 | C2$^{MU}$ | Mutating binding residues | Y2H: METTL14-C2 interaction |
| pGADT7 | CP$^{MU}$ | Mutating binding residues | Y2H: ALKBH4-CP interaction |
| pET28a | METTL14(mu) | Mutating binding residues of METTL14-C2 interaction | UPLS-MS: m$^6$A level on *Vg* |
| pET28a | ALKBH4(mu) | Mutating binding residues of ALKBH4-CP interaction | UPLS-MS: m$^6$A level on *Vg* |
| pET28a | C2(mu) | Mutating binding residues of METTL14-C2 interaction | UPLS-MS: m$^6$A level on *Vg* |
| pET28a | CP(mu) | Mutating binding residues of ALKBH4-CP interaction | UPLS-MS: m$^6$A level on *Vg* |
| pET28a | C2(MU) | Mutating binding residues of METTL14-ssRNAs-C2 interaction | UPLS-MS: m$^6$A level on *Vg*、EMSA: METTL14-RNA binding |
| pET28a | CP(MU) | Mutating binding residues of ALKBH4-ssRNAs-CP interaction | UPLS-MS: m$^6$A level on *Vg*、EMSA: ALKBH4-RNA binding |

**Table S21** Single-stranded RNAs (ssRNAs) used in $m^6A$ detection by UPLC-MS

| **$m^6A$ sites** | **ssRNA substrate** | **METTL3** | **METTL14** | **ALKBH4** |
|---|---|---|---|---|
| 5’UTR-A610 | AGCUC**AGACU**UCAAG | √ | √ | - |
| 5’UTR-A713 | AGCUC**GGACU**GGGGA | √ | √ | - |
| CDS-A621 | AAAAC**GGACA**AGUCA | √ | √ | |
| CDS-A621 | AAAAC**GGA**($m^6A$)**CA**AGUCA | - | - | √ |
| CDS-A884 | GUCAU**GAACU**UGACU | √ | √ | |
| CDS-A884 | GUCAU**GAA**($m^6A$)**CU**UGACU | - | - | √ |
| CDS-1494 | ACAAC**GGACA**CAACG | √ | √ | - |
| CDS-1512 | ACAAC**GGACA**CAACG | √ | √ | - |
| CDS-1530 | ACAAU**GGACA**CAACG | √ | √ | - |
| CDS-5538 | UCCAC**GGACA**CAAUG | √ | √ | - |
| CDS-5738 | CCUCA**AAACU**GUGUC | √ | √ | - |
| CDS-6360 | AAAAC**GGACC**CUCUA | √ | √ | - |
| 3’UTR-A14 | AAUCU**GGACU**UCAGC | √ | √ | - |
| 3’UTR-A14 | AAUCU**GGA**($m^6A$)**CU**UCAGC | - | - | √ |
| 3’UTR-A138 | CCCUCA**AAACU**GUGUC | √ | √ | - |
| 3’UTR-A138 | CCCUCA**AAA**($m^6A$)**CU**GUGUC | - | - | √ |
| 3UTR-A189 | GUCAU**UGACU**CUUCU | √ | √ | - |
| 3UTR-A426 | GACCA**GGACU**CCUCC | √ | √ | - |

**Table S22** The probes used in SELECT assays

| $m^6A$ sites | X/N | up/down | 5'-phos | Sequence (5'-3') |
|---|---|---|---|---|
| 3'UTR-A14 | X | up | - | tagccagtaccgtagtgcgtgAGCTCCGTCATAGATGGCTGAAG |
| | X | down | √ | CCAGATTGGGGGATTAGGCAAcagaggctgagtcgctgcat |
| | N | up | - | tagccagtaccgtagtgcgtgCGTCATAGATGGCTGAAGTCCAG |
| | N | down | √ | TTGGGGGATTAGGCAACACAGcagaggctgagtcgctgcat |
| 3'UTR-A138 | X | up | - | tagccagtaccgtagtgcgtgGTCTTCTGGGTTCTTGTAGACACAG |
| | X | down | √ | TTTGAGGGGCGGTGAAGTCcagaggctgagtcgctgcat |
| | N | up | - | tagccagtaccgtagtgcgtgTGGGTTCTTGTAGACACAGTTTTGA |
| | N | down | √ | GGGCGGTGAAGTCGTCAGcagaggctgagtcgctgcat |
| CDS-A621 | X | up | - | tagccagtaccgtagtgcgtgGGTCTTGACGATGTCGATGACTTG |
| | X | down | √ | CCGTTTTCACGGAGCTTGGcagaggctgagtcgctgcat |
| | N | up | - | tagccagtaccgtagtgcgtgGATGTCGATGACTTGTCCGTTTT |
| | N | down | √ | ACGGAGCTTGGGGAAAGGAcagaggctgagtcgctgcat |
| CDS-A884 | X | up | - | tagccagtaccgtagtgcgtgTGGAAGGAGGCCAAAGTCAAG |
| | X | down | √ | TCATGACACTGACAACCATTCCCTcagaggctgagtcgctgcat |
| | N | up | - | tagccagtaccgtagtgcgtgAGGAGGCCAAAGTCAAGTTCATG |
| | N | down | √ | CACTGACAACCATTCCCTTCTGTGcagaggctgagtcgctgcat |

Note: X, the $m^6A$ site; N, the negative non-$m^6A$ site. These SELECT probes were used for detection $m^6A$ modification at single-base resolution according to Yu *et al.,* (4).

**Table S23** The RNA probes used in EMSA, MST and SPR assays

| $m^6A$ sites | Sequence (5'-3') | 5'-labelled | 3'-labelled | Type |
|---|---|---|---|---|
| 3'UTR-A14 | CCCCCAAUCU**GGACU**UCAGCCAUCU | / | / | Unlabelled |
| | CCCCCAAUCU**GGACU**UCAGCCAUCU | Biotin | CY5 | Labelled |
| | CCCCCAAUCU**AAGUC**UCAGCCAUCU | Biotin | CY5 | Mutated |
| | CCCCCAAUCU**GGA(**$m^6A$**)CU**UCAGCCAUCU | Biotin | CY5 | $m^6A$-taged |
| 3'UTR-A138 | CCGCCCCUCA**AAACU**GUGUCUACAA | / | / | Unlabelled |
| | CCGCCCCUCA**AAACU**GUGUCUACAA | Biotin | CY5 | Labelled |
| | CCGCCCCUCA**GGGUC**GUGUCUACAA | Biotin | CY5 | Mutated |
| | CCGCCCCUCA**AAA(**$m^6A$**)CU**GUGUCUACAA | Biotin | CY5 | $m^6A$-taged |
| CDS-A621 | CCGUGAAAAC**GGACA**AGUCAUCGAC | / | / | Unlabelled |
| | CCGUGAAAAC**GGACA**AGUCAUCGAC | Biotin | CY5 | Labelled |
| | CCGUGAAAAC**AAGUG**AGUCAUCGAC | Biotin | CY5 | Mutated |
| | CCGUGAAAAC**GGA(**$m^6A$**)CA**AGUCAUCGAC | Biotin | CY5 | $m^6A$-taged |
| CDS-A884 | UCAGUGUCAU**GAACU**UGACUUUGGC | / | / | Unlabelled |
| | UCAGUGUCAU**GAACU**UGACUUUGGC | Biotin | CY5 | Labelled |
| | UCAGUGUCAU**AGGUC**UGACUUUGGC | Biotin | CY5 | Mutated |
| | UCAGUGUCAU**GAA(**$m^6A$**)CU**UGACUUUGGC | Biotin | CY5 | $m^6A$-taged |

**Table S24** MST assay of binding affinity of METTL14 to *Vg* m$^6$A sites

| **Fluor. Molecule** | **Ligand** | **$K_D$ [M]** | **$K_D$ Confidence [M]** | **Signal/Noise** |
|---|---|---|---|---|
| C2 | METTL14 | 2.5551E-07 | ± 5.5779E-08 | 14.860821 |
| A14 | METTL14 | 3.6125E-07 | ± 6.6754E-08 | 17.592963 |
| A14 + C2 | METTL14 | 9.7338E-08 | ± 1.4231E-08 | 22.783170 |
| A138 | METTL14 | 1.4070E-07 | ± 2.3075E-08 | 19.993261 |
| A138 + C2 | METTL14 | 4.1096E-08 | ± 4.1557E-09 | 36.123795 |
| A621 | METTL14 | 1.1336E-07 | ± 1.6257E-08 | 23.05557 |
| A621 + C2 | METTL14 | 2.7913E-08 | ± 3.3463E-09 | 31.058159 |
| A884 | METTL14 | 4.0130E-06 | ± 5.3257E-07 | 31.155962 |
| A884 + C2 | METTL14 | 7.3067E-07 | ± 1.7972E-07 | 13.603209 |

**Table S25** MST assay of binding affinity of ALKBH4 to *Vg* $m^6A$ sites

| Fluor. Molecule | Ligand | $K_D$ [M] | $K_D$ Confidence [M] | Signal/Noise |
|---|---|---|---|---|
| CP | ALKBH4 | 1.4768E-07 | ± 2.6179E-08 | 26.840393 |
| A14-$m^6A$ | ALKBH4 | 1.2279E-06 | ± 2.9949E-07 | 22.670097 |
| A14-$m^6A$ + CP | ALKBH4 | 5.1795E-06 | ± 1.0216E-06 | 77.726483 |
| A138-$m^6A$ | ALKBH4 | 2.0523E-06 | ± 4.3169E-07 | 34.251585 |
| A138-$m^6A$ + CP | ALKBH4 | 9.8631E-06 | ± 2.8325E-06 | 115.36537 |
| A621-$m^6A$ | ALKBH4 | 1.6908E-06 | ± 3.4542E-07 | 31.566604 |
| A621-$m^6A$ + CP | ALKBH4 | 8.6898E-06 | ± 4.7929E-06 | 50.823046 |
| A884-$m^6A$ | ALKBH4 | 3.0846E-06 | ± 8.6982E-07 | 33.914434 |
| A884-$m^6A$ + CP | ALKBH4 | 1.1266E-05 | ± 9.6756E-06 | 46.334749 |

**Table S26** SPR assay of binding affinities of METTL14 or ALBH4 to *Vg* $m^6A$ sites in the presence of C2 or CP protein

| Ligand | Analyte | $K_a$ (1/Ms) | $K_d$ (1/s) | $K_D$ (M) | Rmax |
|---|---|---|---|---|---|
| METTL14 | 3'UTR-A14 | 1.77e+04 | 1.74e-02 | 9.82e-07 | 99.90 |
| | C2 (+) | 1.86e+04 | 9.67e-03 | 5.19e-07 | 99.90 |
| | CDS-A621 | 2.31e+04 | 7.52e-03 | 3.26e-07 | 98.38 |
| | C2 (+) | 2.78e+04 | 2.37e-03 | 8.53e-08 | 98.38 |
| ALKBH4 | 3'UTR-A14 | 2.03e+04 | 4.20e-02 | 2.07e-06 | 99.73 |
| | CP (+) | 6.88e+03 | 5.17e-02 | 7.52e-06 | 99.73 |
| | CDS-A621 | 3.02e+03 | 1.35e-02 | 4.47e-06 | 98.22 |
| | CP (+) | 1.72e+03 | 1.68e-02 | 9.75e-06 | 98.22 |

### Other Supplementary Material for this manuscript includes the following:

**Videos S1-S5.** Outbreaks of the whitefly *B. tabaci* across China.

**Dataset S1.** Sequences of CDS of *Ex*, *Va*, *Bg* and *Vg*.

**Dataset S2.** 5’ UTR and 3’ UTR sequences of *Vg* gene by RACE cloning.

**Dataset S3.** Sequences of the TYLCV structure genes.
